# MicrobeMS - A MATLAB Toolbox for Microbial Identification Based on Mass Spectrometry

**DOI:** 10.64898/2026.05.08.723807

**Authors:** Peter Lasch

## Abstract

Over the last two decades, matrix-assisted laser desorption/ionization time-of-flight mass spectrometry (MALDI-ToF MS) has become the standard method for identifying bacteria and has found a wide range of applications, especially in clinical microbiology. The method’s high taxonomic resolution, minimal sample preparation, and complete, ready-to-use commercial systems, which include instrumentation, experimental protocols, spectral databases, and identification analysis software, were key factors in the success of MALDI-ToF MS as the standard for identifying microorganisms in routine diagnostic laboratories. However, despite the availability of these commercial solutions, there is also a growing need for efficient, cost-effective, vendor-neutral databases and analysis tools. These tools would enable the compilation of user-defined mass spectral databases and the testing of new analysis methods and algorithms, particularly in an academic context. To this end, MicrobeMS software has been developed to cover all stages of MALDI-ToF MS-based identification analysis. MicrobeMS is an easy-to-use desktop application for analyzing mass spectra from microorganisms and performing tasks related to spectrum database compilation. It includes routines for direct data import and export, biomarker peak searches, management of spectrum metadata, testing of spectrum quality, supervised and unsupervised identification analysis and intuitive result display. MicrobeMS is implemented in MATLAB and is freely available as MATLAB pcode for Windows and Linux, as well as a standalone application. Over the last fifteen years, the software has undergone continuous development and is now used routinely in various settings at the Centre for Biological Threats and Special Pathogens (ZBS) at the Robert Koch Institute (RKI) in Berlin, Germany, for example in supporting spectrum database compilation, to identify special or rare pathogenic bacteria by advanced identification analysis concepts, or to test *in silico* MALDI-ToF MS databases derived from microbial genomes.

In this software publication the versatility and capabilities of MicrobeMS are demonstrated using a test data set from highly pathogenic bacteria (HPB) which has been obtained as part of a published European Union (EU)-funded External Quality Assurance Exercise (EQAE). MicrobeMS and HPB test data can both be downloaded from https://wiki.microbe-ms.com/. The goal of this software publication is twofold: to raise awareness of MicrobeMS within the scientific community and to encourage the testing of the software and custom-developed MALDI-ToF MS databases of the RKI, which are published at the ZENODO data repository (https://doi.org/10.5281/zenodo.7702374).

## 2. INTRODUCTION

Matrix-assisted laser desorption/ionization time-of-flight (MALDI-ToF) mass spectrometry (MS) has become a standard method for identifying pathogenic bacteria in routine clinical diagnostics over the past 20 years [1, 2]. The method enables bacteria to be diagnosed quickly, reliably, and cost-effective with high sample throughput and only limited material and labour requirements. In addition to its widespread use in clinical microbiology, the MS-based workflow for bacteria has also been established for other areas of application. These include environmental microbiology, [3] analysis of drinking water, [4] food quality control, [5, 6] veterinary medicine, [7] and biosafety applications i.e. for the diagnosis of highly pathogenic bacteria (HPB) classified as biosafety level 3 (BSL-3). [8, 9]

The MALDI-ToF MS identification workflow involves systematically comparing the experimentally recorded mass spectra of a microbial sample with an *a priori* unknown taxonomic identity with reference spectra stored in databases whose taxonomic status has been carefully determined by established reference methods. Spectral distance or similarity measures, scores, are often calculated and used to create score ranking lists. Ideally, these lists, along with the scores and taxonomic assignments of the reference spectra, can be used to determine the genus and species identities of the unknown test organisms. [1]

In 2002, two units at the Robert Koch Institute (RKI) initiated a long-term project that aimed at evaluating the potential of MALDI-ToF MS for identifying highly pathogenic microorganisms. An *autofleX I* MALDI-ToF mass spectrometer from Bruker Daltonics was obtained, and in the following two units at the Center for Biological Threats and Special Pathogens (ZBS) — the *Highly Pathogenic Bacteria* Unit (ZBS2) and the *Proteomics and Spectroscopy* Unit (ZBS6) — collaborated closely to develop and test a reliable, MALDI-ToF MS-compatible method for inactivating highly pathogenic bacteria (HPB) [10, 11]. Upon completion of this initial development phase, the TFA-based inactivation method (TFA: trifluoroacetic acid) was widely adopted for collecting MALDI-ToF mass spectra from HPB and related bacteria [9, 12–15]. In the context of this long-term project, partly carried out in collaboration with the MALDI technology pioneers Anagnostec and Bruker Daltonics, we then compiled databases containing mass spectra of HPB including *Bacillus anthracis*, *Brucella* spp., *Yersinia pestis*, *Francisella tularensis*, *Burkholderia mallei* and *Burkholderia pseudomallei*, as well as spectra of their close and more distant relatives [16, 17]. The databases have been published freely and have undergone continuous development in subsequent years for the benefit of the general public [16].

In addition to the aforementioned activities, in-house tools for analyzing mass spectrum data were developed and implemented throughout the duration of the RKI’s HPB-MALDI project (see [10, 12, 18]). In the early years of the project, routines for the taxonomic identification of bacteria had to be developed, as integrated, ready-to-use solutions such as VITEK MS (bioMerieux, France) [19] or the MALDI Biotyper (MBT, Bruker Daltonics, Germany) [20] were still not available. Consequently, the necessity for software tools became apparent in three areas: firstly, to facilitate the identification of bacterial pathogens, secondly, to organize and manage the ever-growing in-house spectrum databases, and thirdly to identify discriminating spectral markers, i.e. taxon-specific mass peaks. Although developing a comprehensive, feature-complete software product for the mass spectrometric analysis of bacteria was initially not a declared goal, MicrobeMS has evolved over the years into a software solution that I believe could also be helpful outside the RKI.

MicrobeMS has been written in MATLAB and, in its current version, includes a variety of functions in addition to microbial identification analysis. These functions include, among others, various import and export routines, functions for spectral preprocessing, spectral quality tests, hierarchical cluster analysis as a way for unsupervised classification, and interfaces to neural network software [21]. Almost all of these functions have been developed for, and used in various studies with MALDI-ToF MS applications from ZBS6, the RKI’s *Proteomics and Spectroscopy* Unit [13, 16, 22–25].

It is important to emphasize at this point that the present software description is primarily intended for potential users in an academic environment. While the program has demonstrated its efficacy in the domain of MS-based microbiological diagnostics by the RKI’s consulting laboratories for HPB and regularly facilitates achieving accurate identification results from special, or rare, bacterial pathogens, neither the software nor the RKI databases have undergone formal validation. This distinguishes MicrobeMS from integrated complete solutions available from commercial suppliers. Nevertheless, apart from cost aspects, MicrobeMS’s emphasis on applications in academic settings offers distinct advantages. While commercial products must be optimized for traceability and high safety levels in use, must be validated, and should thus be adapted cautiously due to their intended use in the regulatory context of clinical diagnostics, research software is allowed to be designed with greater flexibility. For example, software for academia may offer a variety of different analysis options, enabling the systematic testing of algorithms and programme settings. Furthermore, the data exchange capabilities inherent in commercial diagnostics software products do not always align with the preferences of researchers. For obvious reasons, it is not desirable for manufacturers if users leave their software ecosystems. The respective vendors have invested heavily in method development and compiling spectral database, so data exchange is not a priority. Furthermore, also the users expect commercial diagnostics software to offer a different range of features. In high-throughput microbiological diagnostics, the ease of use, the focus on key functions, and the reliability of application are more important than the widest possible range of functionalities or the availability of different interfaces for data exchange.

MicrobeMS does not pursue commercial interests. The software is designed with a primary focus on academic users and facilitates straightforward access to spectral data, thereby supporting the development and evaluation of novel analysis methods. In addition, MicrobeMS offers the possibility to conduct spectrum analyses in a local, restricted environment. This characteristic sets MicrobeMS apart from open web-based resources for identifying microorganisms, such as MicrobeNet, [26] IDBac, [27] or MSI-2 [28] and is advantageous or required in cases where sending sample data to third parties is unwanted or not permitted.

This software description serves two primary purposes: first, it aims to inform potential users of MicrobeMS regarding the software’s range of functions, applications, and advantages and disadvantages in comparison to alternative solutions; and second, it seeks to elicit comprehensive feedback from users.

## 3. SOFTWARE DESCRIPTION

The MicrobeMS software package is specifically designed for analyzing mass spectra from microbial samples. It was developed over the last two decades as part of the RKI’s mass spectrometry projects for analyzing both, MALDI-ToF MS and later then LC-MS1 data. It features a graphical user interface (GUI) for controlling the program and entering program operation and analysis parameters. A text window is also included, in which additional program output and error messages are displayed. By default, the contents of the text window are stored in a log file for documentation purposes. MicrobeMS represents a comprehensive MATLAB-based package that can operate either as a standalone application or as MATLAB m-, or p-code under the Windows, or Linux operating system. An easy-to-use installer is available for the Windows stand-alone package, which automatically extracts the program, test data, and configuration files from the cabinet. It also performs all the necessary steps to make MicrobeMS operational.

MicrobeMS was initially developed to analyze MALDI-ToF MS [18] data and later then adapted for working with LC-MS1 data [29]. Currently, MicrobeMS’s primary application is analyzing MALDI-ToF mass spectra of pathogenic bacteria. Accordingly, this exposition of the software’s particulars refers to version 0.92 from January 2026 and is focused on its MALDI applications.

Original MALDI-ToF mass spectra in formats defined by Bruker Daltonics (BrukerRaw, *fid*) or Shimadzu (via the mzXML data format) can be imported and processed, and then converted into MicrobeMS-specific MATLAB data matrix formats. Additional MATLAB functions are available for importing LC-MS^1^ data in Bruker’s tims-ToF format (proprietary .*d* folder structure) or Thermo’s Orbitrap *(.raw*) format. However, these import functions are not part of the current program version.

MicrobeMS allows standard MALDI-ToF MS preprocessing manipulations such as smoothing, baseline correction, normalization, peak detection, auto-calibration to mention some of them. Furthermore, the software’s functionalities include spectrum quality testing (QT), microbial identification analysis based on inter-spectral distances and machine learning (ML) methods (e.g., artificial neural networks [ANN]), visualization of identification results, unsupervised hierarchical cluster analysis (UHCA), biomarker analysis, *pseudo-gel* view generation, and microbial mass spectra database management, which includes graphical user dialog-guided functionalities for adding, editing, and manipulating spectrum metadata. Since the software runs in a Windows, or Linux 64-bit environment, the number of spectra in the data sets is limited only by the amount of available computer memory (RAM) allowing analysis of thousands of mass spectra during a program session. A screenshot of the main gui is given in Figure 1 and an overview of MicrobeMS main functionalities is given by Figure 2.

**Figure 1.**
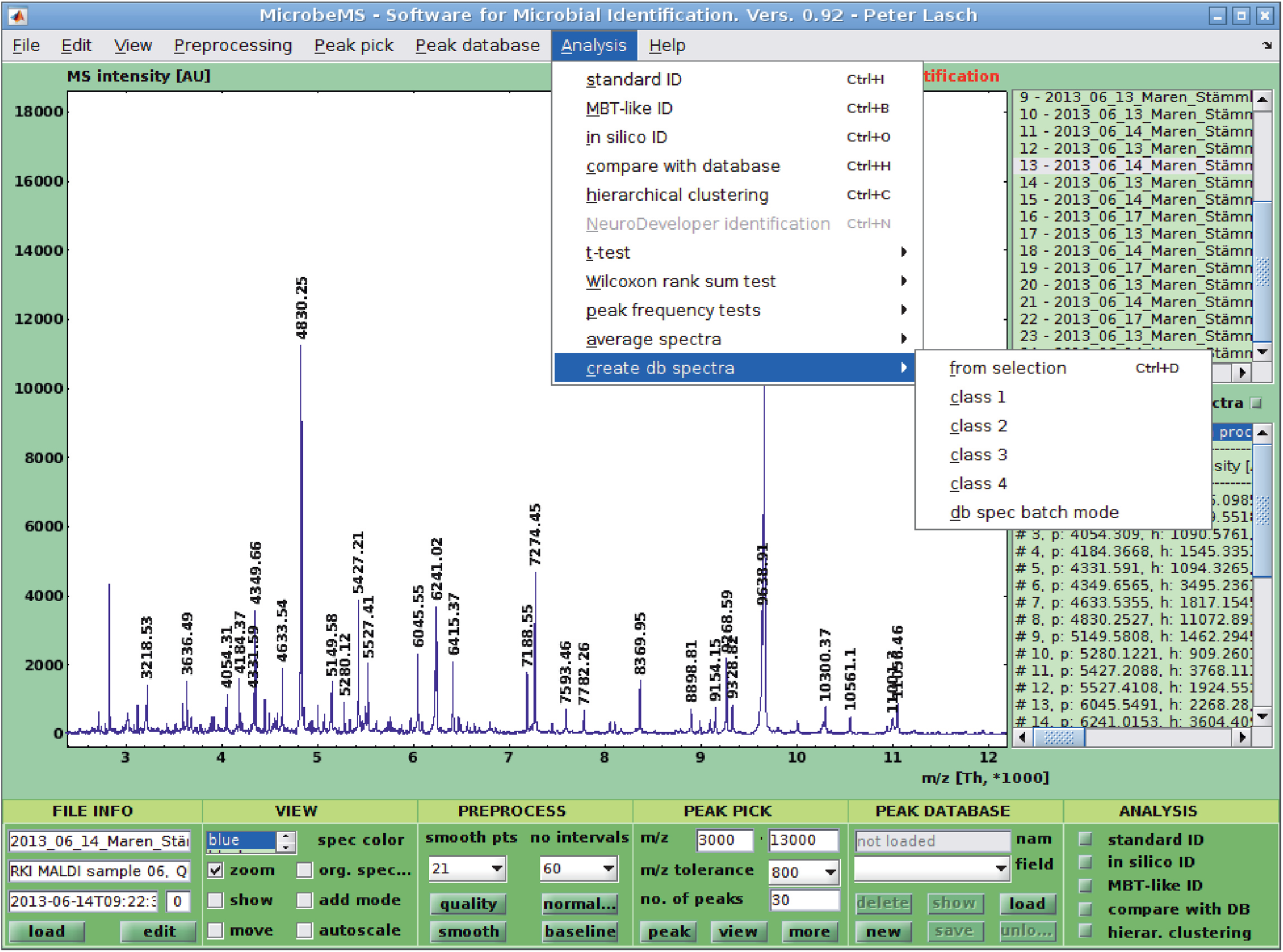
Main user interface of MicrobeMS version 0.92.

**Figure 2.**
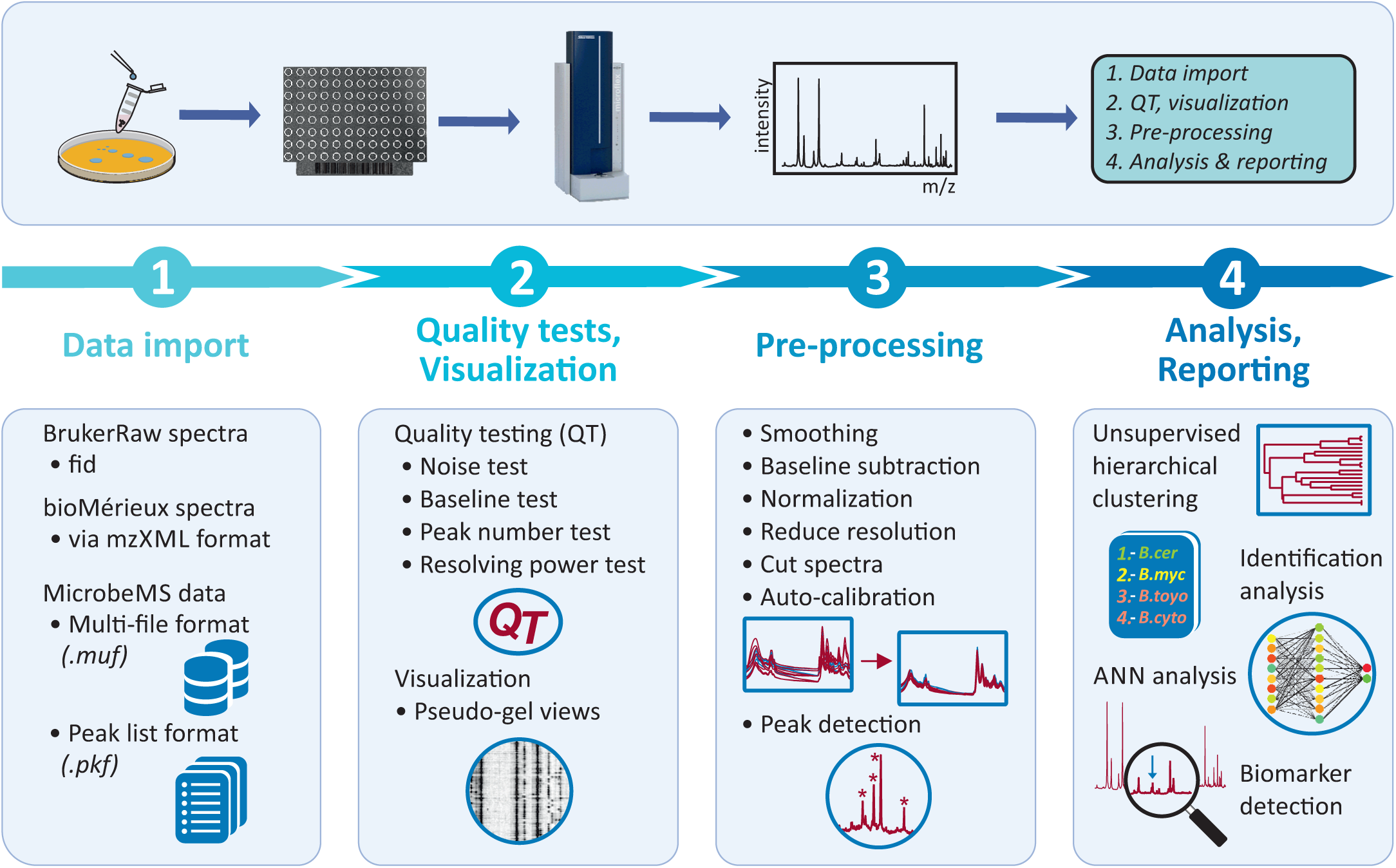
MicrobeMS’ main software options for analyzing bacterial MALDI-ToF mass spectra.

An important feature of the MicrobeMS software is its ability to identify bacteria through a comparison of experimental MALDI-ToF mass spectra with spectrum entries contained in a spectral database. Typically, replicates of MALDI-ToF mass spectra from microorganisms are recorded within the mass-to-charge ratio (m/z) range between 2,000 and 20,000 Th. Spectrum databases ideally contain a representative selection of validated spectrum entries (consensus spectra), referred to in MicrobeMS as *database* spectra.

The principle of bacterial identification using MALDI-ToF MS involves only very few basic steps: First, mass spectra from unknown, cultivated bacterial isolates are acquired by a dedicated instrument using standardized settings for cultivation, sample preparation and data acquisition. These spectra can then be queried against a database of MALDI-ToF mass spectra, the taxonomic identity of which is known. Through pattern matching, the specific MALDI fingerprints allow identifying the tested bacteria, with the degree of match defining the taxonomic level of identification [16].

This relatively simple principle for the spectrum-based identification of bacteria was proposed by Dieter Naumann more than 35 years ago, although FTIR spectroscopy instead of MALDI-ToF mass spectrometry was employed [30–32]. The described workflow was later used by commercial vendors for MALDI-ToF MS and recently adapted by this author’s RKI group to identify pathogenic bacteria using LC-MS^1^ data [29].

MicrobeMS contains a number of automated functions to make the most frequently used routines more user friendly and convenient. Routines such as QT, database compilation, and identification analyses based on spectral distances and experimental or *in silico* MALDI-ToF databases are largely automated, tough with modifiable default parameters. For example, the distance-based standard identification method runs the following software routines sequentially: QT, spectral preprocessing (smoothing, baseline correction, normalization), peak detection, and loading a standard peak list library. All parameters used by these automated routines are contained in an editable, documented text file (*microbems.opt*; see the Wiki for details) [33]. When MicrobeMS is initialized, it parses a plain text file containing a wide range of program-wide parameters for QT, auto-preprocessing, and auto-peak detection, among others.

## 4. MAIN SOFTWARE FEATURES

### 4.1 Data import and export

MicrobeMS supports importing Bruker MALDI-ToF mass spectral data in the form of the standard *BrukerRaw* data format. *BrukerRaw* structures compose a specific folder structure that contains time-of-flight (ToF) data in a binary format (*fid* files) and text-formatted acquisition files named *acqu, acqus, proc, procs,* etc., in which original and/or processed metadata, spectrum IDs, calibration information, etc. are contained. The *BrukerRaw* structure format is described with some detail in Schmidt-Santiago et al., 2025 [34]. To import *BrukerRaw* spectra into MicrobeMS, users can select single, or multiple spectrum directories via a custom dialog box, thus allowing importing large directory structures that may contain hundreds of individual MALDI-ToF mass spectra. Furthermore, VITEK MS (bioMerieux) spectrum data can be imported via the mzXML data format enabling cross-vendor studies. The respective MicrobeMS functionality relies on the MATLAB *importmzxml* function. Spectral data sets can furthermore be imported or exported via a custom-designed MATLAB data format (.*muf*, multi-spectrum format). These files are designed to contain sets of original and eventually existing pre-pre-processed MALDI-ToF MS intensity spectra, respective peak list data, and the complete set of metadata. For data exchange, .*muf* files can be directly imported in MATLAB by entering the following expression at the MATLAB prompt: *spec = load(’sample_set.muf’, ‘-mat’);*. The variable *spec* denotes a MATLAB structure array, and *sample_set.muf* is the *.muf*-file name. A detailed description of the *.muf* file format can be found at the MicrobeMS Wiki [33].

MicrobeMS permits furthermore storing MALDI-ToF spectrum data in one of the following formats: MicrobeMS multifile format (.*muf*, see above), Bruker’s raw spectrum format (experimental, vide supra) and in an ASCII data format (tabular data, space, or comma separated). The *so-called* peak list data format (.*pkf*) represents a second type of a custom MicrobeMS data format. Such files serve the purpose of exporting or importing peak list data derived from MALDI-ToF MS, or LC-MS^1^ data. Peak lists are required for hierarchical cluster analysis and identification analyses. The format of .*pkf* data files is also described at the MicrobeMS Wiki [33]. Peak list data files are accessible in MATLAB’s workspace by entering *peaks = load(‘peaklist_set.pkf’,’-mat’);* at MATLAB’s command line. The variable *peaks* represents a MATLAB structure array; its format is described in detail at the MicrobeMS Wiki [33]. With .xml formatted files MicrobeMS supports a second type of peak list files. Such files can be processed by the CDC’s MicrobeNet web platform, a free online resource and database that supports clinical and public health laboratorians to identify rare and unusual bacteria [26]. Based on pre-defined parameters stored in its parameter file *microbems.opt*, MicrobeMS enables the user-friendly, one-stop conversion of native Bruker-formatted spectrum data into the *.xml* format.

### 4.2 Add / edit spectrum metadata

MicrobeMS offers various methods for adding, modifying, and extracting metadata from microbial mass spectra. This metadata can include taxonomic information, cultivation parameters (e.g., time, temperature, and medium), the sample preparation method, the operators’ names, and other relevant information. Metadata can be stored directly with Bruker Daltonics’ original MALDI-ToF mass spectra files (*BrukerRaw*). Direct access to metadata is often important during spectral analysis and interpretation of results, for example, when interpreting dendrograms or *pseudo-gel* views (see below). The RKI workflow fulfills this requirement by leveraging a MicrobeMS capability that incorporates spectral metadata into *BrukerRaw* data. A detailed description of the procedure is beyond the scope of this article, but it can be found in the MicrobeMS Wiki [33]. Metadata stored in *BrukerRaw* spectrum files is automatically transferred into MicrobeMS upon spectrum import and can be modified manually later via the ‘e*dit metadata*’ menu item. Figure 3 shows a screenshot of the corresponding user dialog box. Additionally, selected metadata can be output to the MATLAB command window and the log file, which routinely records the events, activities, and results of a MicrobeMS analysis run (*view* → *display spectra entries* menu bar).

**Figure 3.**
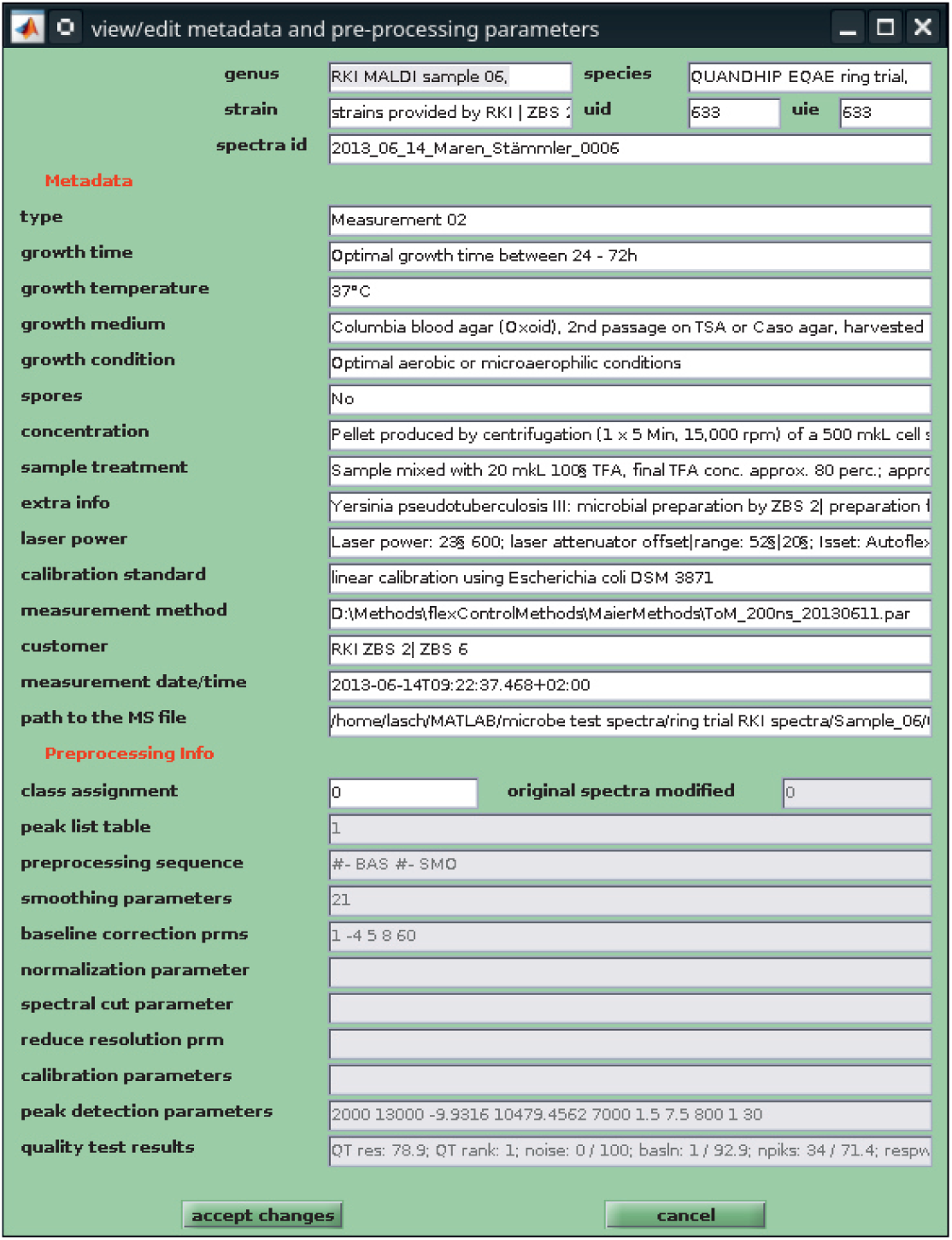
Screenshot of the user dialog box *View/edit metadata and preprocessing parameters.* During the early stages of MicrobeMS, the development of procedures and functions for managing spectrum metadata benefited greatly from discussions with staff at Anagnostec, a pioneer in the field of MALDI-ToF MS-based microbiology (Dr. M. Erhardt). From 2003 to 2006, the RKI and the then-startup company engaged in collaborative projects, which included intensive scientific exchange. In 2010, the company bioMerieux acquired Anagnostec and their mass spectrometry-based solutions, including the SARAMIS technology developed there.

A script that automates adding spectrum metadata from Excel table content is explained in detail in the MicrobeMS Wiki [33].

### 4.3 Quality tests

The MicrobeMS software’s quality testing function was introduced to provide guidance and support to less experienced experimenters that use the manual measurement mode. It allows such users to assess, based on objective criteria, whether the recorded mass spectra are suitable for identification analysis or to be included databases. The QT routine is also extremely helpful in automatic measurements because it helps to save time by identifying and sorting out MALDI-ToF mass spectra that are of insufficient quality.

Spectral quality tests (QT) can be used to evaluate spectra based on predefined quality criteria and, in the event of insufficient quality, exclude them from further analysis. MicrobeMS enables the automated execution of four different tests on chosen MALDI-ToF mass spectra, with a quality score being determined for each test, which is then incorporated into the calculation of an overall quality score. Specifically, the QT implemented in MicrobeMS can automatically evaluate the following MALDI-ToF mass spectrum quality criteria: (i) spectrum noise, (ii) number of peaks per spectrum, (iii) baseline elevation, and (iv) mean resolving power of the identified mass peaks. Quality testing involves the automated determination of individual scores for each of the four QT criteria. Each score ranges between values of zero (poor quality, test failed) and 100 (excellent quality). From these scores a weighted overall QT score is then determined using weightings of the underlying test scores. The overall quality score varies between zero and 100 and has proven useful for detecting outliers by means of user-adjustable objective test criteria. All QT parameters are adjustable through the software. The results of quality testing can be visualized in an HTML-formatted document whereas the spectral quality achieved is encoded using a traffic light scheme (see Figure 4). Furthermore, QT results can be stored in a MATLAB data format, opening possibilities for further statistical analyses of spectral quality parameters (used for example in ref. [16]).

**Figure 4.**
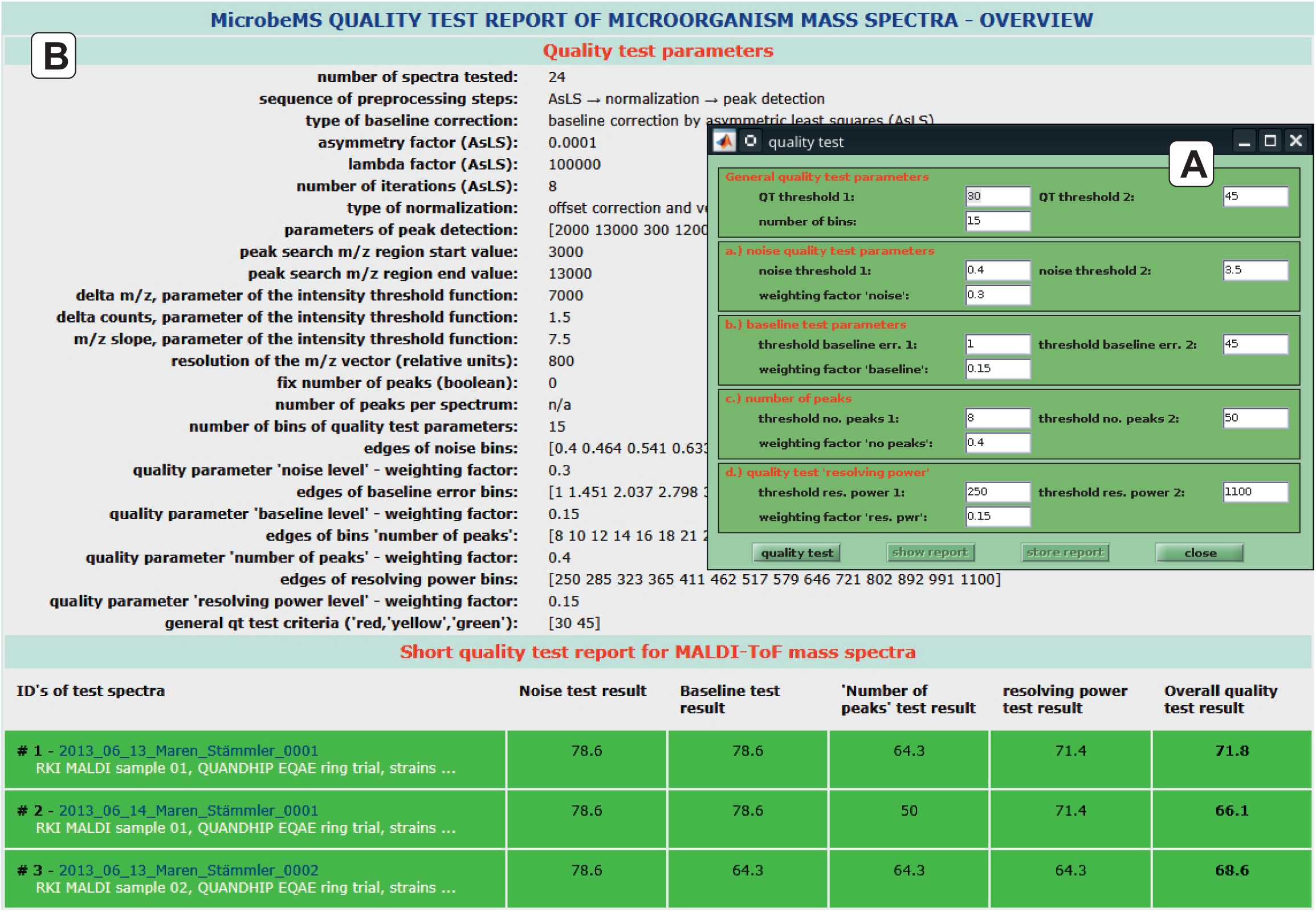
Quality test of MALDI-ToF mass spectra. Screenshot of the QT dialog box (A) and of the QT report (HTML formatted, B).

### 4.4 Preprocessing

Preprocessing is an important first step of spectrum analysis that aims, among others, at increasing the robustness and accuracy of subsequent analysis tasks, improving the data’s interpretability, and removing irrelevant and/or redundant information. [35] MicrobeMS allows the following preprocessing tasks to be performed: (i) autocalibration, (ii) baseline subtraction (minimum method, or asymmetric least square method [36]), (iii) smoothing (average smoothing, Savitzky-Golay smoothing [37]) (iv) normalization by a modified 1-norm method, (v), reduction of spectrum resolution, and (vi) cutting (truncating) spectra in the low and/or high m/z range. Autocalibration, the first preprocessing method, requires knowledge of the precise positions of at least two mass peaks of the analyte per spectrum, i.e. peak tables from adequately pre-processed mass spectra are needed in order to correct for inadequate spectrum calibration during data acquisition.

### 4.5. Peak picking

Peak picking is an essential and the final step of MicrobeMS’ preprocessing pipeline. The result of peak picking is a peak table attached to the mass spectra. In MicrobeMS peak picking can be initiated using the *Find Peaks* command from the *Peak Pick* menu bar.

MicrobeMS uses exclusively own algorithms for peak detection, i.e. existing peak tables generated by other software packages are ignored. The key idea behind peak picking in MicrobeMS is based on two observations: First, the analytical sensitivity in MALDI-ToF mass spectrometry is inversely related to the x, i.e. the m/z position: The method’s sensitivity is higher in the low m/z range and lower in the high m/z range. Second, it is often convenient to define a predefined (fixed) number of peaks per mass spectrum. These two observations are addressed by a procedure that uses an S-shaped curve as an intensity threshold function, defined as a special type of generalized logistic function. For peak detection, peaks exhibiting intensities that exceed the threshold function at the given m/z positions are considered, while those below the threshold are disregarded. The logistic function produces higher intensity threshold values in the low m/z region and lower values in the high m/z range. In MicrobeMS, the threshold function for peak picking is defined as the sum of the following m/z dependencies:

1. A baseline function obtained by baseline correction by asymmetric least squares, [36]
2. an S-shaped sigmoid function *T(x)* that defines MALDI-ToF MS threshold intensities as a function of x, i.e. the m/z values, and
3. a noise distribution function scaled by an adjustable factor.

The sigmoid function *T(x)* is considered a special type of a generalized logistic function, which is defined by the following equation:

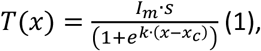

where *k* = *e*^−*l*^, with *l* defining the slope of the threshold function around a given point *x_c_*. For a given mass spectrum, the intensity *I_m_* is defined as the mean of the 13[-3] most intense mass peaks whereby the expression 13[-3] indicates that among the thirteen most intense mass peaks the top three mass peaks are ignored. This approach is intended to ensure that spectrum normalization is not dictated by the intensities of a few highly intense mass peaks. Furthermore, the variable *s* denotes a scaling factor of *I_m_*, and *x* represents the x-vector data in Thomson (Th) m/z units.

Peak picking should preferentially be carried out after preprocessing. MicrobeMS allows for peak picking by specifying a constant, i.e. fixed number of peaks per spectrum, usually 30, as the default setting. If the number of peaks to be determined is fixed, the threshold function component (ii), *T(x)*, is scaled iteratively by adjusting *s* until the specified number of peaks is reached. Otherwise, the peak tables will only contain peaks with an intensity greater than the non-scaled threshold function at a given factor *s*.

### 4.6. Data visualization by pseudo-gel view representations

Manually screening each spectrum within a reasonable time frame is often not feasible when dealing with large spectra databases, which sometimes comprise hundreds of individual mass spectra. In such cases, simulated gel views, also called *pseudo-gel* views, are a popular means of performing quick visual inspections. In these representations, mass spectral intensities are converted to color, or gray-scale and plotted as functions of the spectrum index (y-axis) and the m/z values (x-axis).

In the MicrobeMS implementation, *pseudo-gel* views are produced using log-transformed MS intensity values and are typically generated from pre-processed (i.e. normalized), internally recalibrated MALDI-ToF mass spectra. The MicrobeMS toolbox allows conveniently generating gel views from hundreds of mass spectra and offers features for manually selecting m/z regions, or defining the scale range that indicates how the intensity data is mapped to the colormap chosen. MATLAB’s *MouseOver* (WindowButtonMotionFcn) functionality is used programmatically for inspecting m/z peak positions and displaying metadata of the underlying mass spectra.

Figure 5 shows an instance of a *pseudo-gel* view. In this example, over 1,400 spectra from the closely related *Bacillus* species, *Bacillus cereus* sensu stricto (s.s.) and *Bacillus anthracis*, were first pre-processed, i.e. smoothed, normalized, baseline-corrected and then subjected to the *pseudo-gel* view visualization function. The vertical *gel bands* illustrate the consistent presence of mass peaks at defined m/z positions. Systematic differences in these bands among genera, species, or strains may indicate the existence of taxon-specific biomarker peaks. (cf. also the legend of Figure 5).

**Figure 5.**
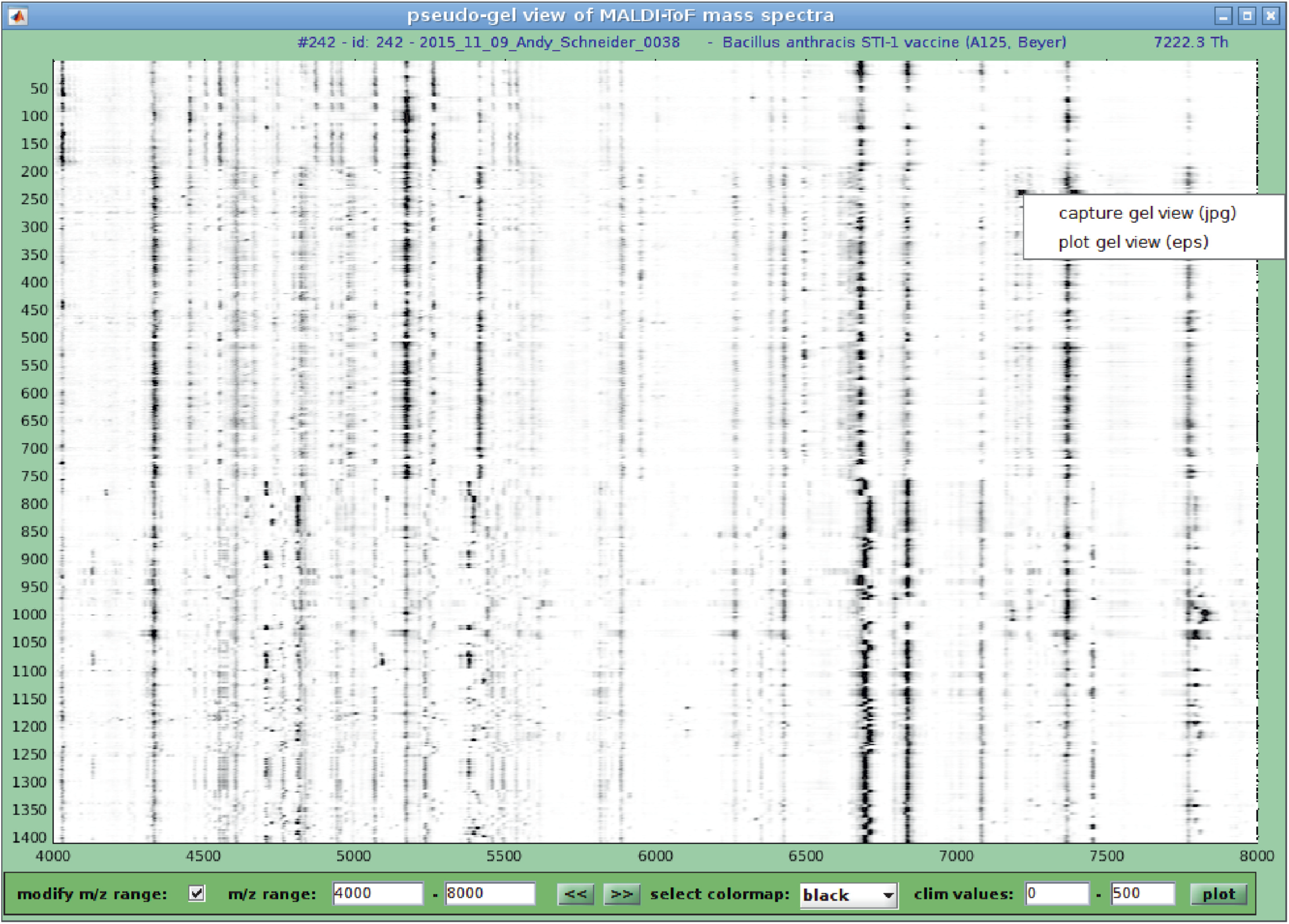
User interface to construct *pseudo-gel* views from pre-processed microbial mass spectra. In such representations, the spectral peak intensities are converted to grey, or color scales and plotted as a function of the spectrum indices and the m/z coordinates. The generation of gel views typically involves normalization, baseline correction and internal re-calibration of the microbial mass spectra. The *pseudo-gel* view presented here displays pre-processed experimental mass spectra from the Bacillus dataset, which is a part of the ZENODO dataset [17]. The x-axis records the m/z values, while the y-axis displays the running spectrum number. In the MicrobeMS implementation, log-scaled spectral intensities are expressed by a grey, or color code.

### 4.7. Creating consensus, or database spectra

Database spectra are the references entries in spectrum libraries of MicrobeMS and are ideally obtained through multiple measurements of a bacterial isolate under study. Technical replicate measurements from biological replicates, i.e. from independent microbial cultures, aim at covering of the spectral variability which is typically observed. Under optimal conditions, the number of individual mass spectra used for database spectrum generation should vary between 12 and 20, and should ideally include data from three biological replicates with at least four technical replicate spectra, respectively [16, 38].

The concept of consensus, or database spectra involves automated generation of peak tables from a series of spectrum measurements of a given microbial strain. These tables contain all the necessary information for subsequent microbial identification analysis. Technically, a database spectrum is represented by a mean mass spectrum linked to a specific type of peak table. MicrobeMS’s identification procedures only use peak table information. Unlike peak tables obtained from single experimental mass spectra, peak tables of database spectra contain peak abundance data, in addition to the m/z peak positions and intensity values. Peak abundance data is defined as the relative peak frequency information which is derived from peak tables of mass spectra used to compile a database spectrum.

The MicrobeMS toolbox enables the automated generation of database spectra, with the parameters for spectrum preprocessing and peak picking taken from an editable parameter file (*microbems.opt*). Database spectra can be obtained in different ways: From labeled sets of mass spectra, from manually selected spectra, or in *batch mode*. The latter mode enables the creation of large series of database spectra in a fully automated manner. The current RKI MALDI-ToF MS spectrum database, which contains more than 1,700 database spectra from highly pathogenic bacteria and their close and more distant relatives, [17] was created using this option.

### 4.8 Unsupervised hierarchical cluster analysis (UHCA)

The primary purpose of the UHCA function is to categorize patterns, i.e. spectra, or peak tables thereof, into groups that are both meaningful and useful, utilizing some type of similarity measure. UHCA is considered a data-driven (unsupervised) classification technique, which is helpful for extracting information from unclassified patterns during an exploratory phase of pattern recognition. MicrobeMS enables for the hierarchical clustering of mass spectra from peak table data; the procedure requires microbial mass spectra obtained via a standardized workflow encompassing reproducible cultivation and data acquisition, as well as utilization of identical preprocessing and peak detection parameters. The UHCA analysis procedure involves two basic steps: calculating a distance matrix (similarity assessment) followed by performing stepwise clustering, i.e. sequential merging of similar clusters. MicrobeMS’s UHCA default distance function uses *D-values* [39, 40], which are derived from Pearson’s product-moment correlation coefficients (PPMCC). Furthermore, MicrobeMS allows users to obtain distance matrices from peak table data based on Euclidean, standardized Euclidean, and City Block distances. The following methods are available for hierarchical clustering: Ward’s method (the default method), single linkage, average linkage, complete linkage, and centroid linkage.

Dendrograms, graphical representations of cluster analysis results, can be generated and visualized using the UHCA user dialog box (Figure 6). MicrobeMS facilitates interpreting and preparing publication-ready dendrograms by enabling users to export vector graphic dendrogram data in PDF or EPS format and output cluster membership lists at the MATLAB command prompt.

**Figure 6.**
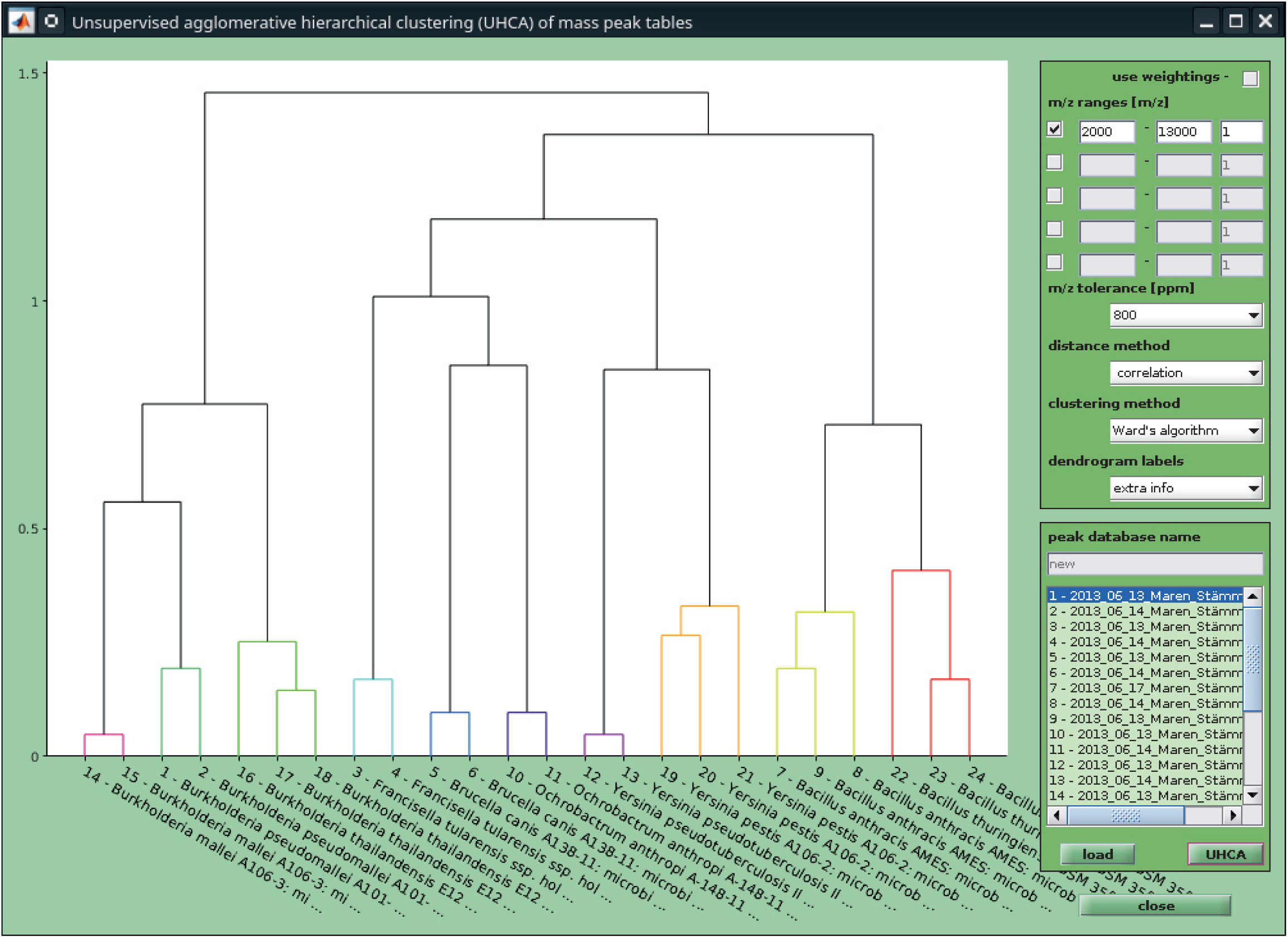
Dendrogram generated by unsupervised hierarchical cluster analysis (UHCA) from mass spectra of the HPB test panel [9] (see also section 5.1). In the present example, peak tables were obtained from pre-processed individual mass spectra. Spectra were first pre-processed before peak tables containing 30 peaks per spectrum were obtained. Technical replicate spectra from the microbial strains tested form strain-specific clusters. Furthermore, clusters of phylogenetically closely related species such as *Yersinia pestis* and *Yersinia pseudotuberculosis*, *Bacillus cereus* group species (*B. cereus s.s.*, *B. thuringiensis*, and *B. anthracis*), *Burkholderia pseudomallei* complex species (*B. thailandensis*, *B. mallei*, and *B. pseudomallei*), or *Brucella canis* and *Ochrobactrum anthropii* form higher order clusters. This supports the statement that MALDI-derived dendrograms may be helpful to study phylogenetic relationships of bacteria.

### 4.9 Identification analysis

Identification of unknown bacteria based on peak list data from reference bacteria represents a pivotal functionality of MicrobeMS. To this end, the software offers peak pattern recognition using artificial neural networks (ANNs, see section 7) as well as distance-based methods as the default approach. The latter group of methods was introduced into the field of microbial identification by Naumann et al. in their seminal work in the late 1980s [30, 39, 41]. In their studies, Naumann and colleagues utilized Fourier-transform infrared (FTIR) spectroscopy and were the first to propose spectral distances as measures of taxonomic distances between unknown bacteria and pathogens of which the taxonomic identities are known. As distance measures Naumann and colleagues suggested *so-called* distance, or *D-values*, which were calculated on the basis of PPMCCs between pairs of bacterial FTIR spectra [42].

During the development of MicrobeMS, the aforementioned principle of spectroscopy-based identification was adapted for use in MALDI-ToF MS. MALDI-ToF MS analyses differ from vibrational spectroscopic analyses in that peak tables rather than spectra are commonly used. This is in contrast with the processing of first or second derivatives obtained from IR absorbance or Raman intensity spectra, which are typically employed in vibrational spectroscopic analyses.

Pearson’s product-moment correlation coefficient *r(x,y)* can be calculated between two spectrum vectors *x* and *y* using equation (2). The PPMCC is then transformed into the *D-values* suggested by Naumann et al. using equation (3). *D-values* range from 0 (perfect correlation, identity) to 2000 (anti-correlation). In order to perform correlation analyses, peak tables have to be truncated to a user-specified window and then back-transformed into discretized low-resolution mass spectra in which broad discretized m/z bins not only account for mass accuracy limitations of the MALDI-ToF MS method but help also to reduce computational requirements.

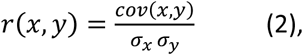

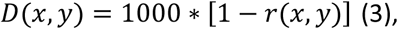

To calculate the PPMCC values *r(x,y)* between spectrum *x* and *y*, the covariance between the two vectors is first determined and then divided by the product of the respective standard deviations. *D-values* are then obtained using equation (3).

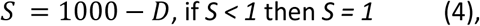

For further identification analyses, it was useful to convert *D-values* of matrix *D* into similarity, or *S-values* using equation (4). *S-values* may range from 1000 (identity) to -1000 (anti-correlation). However, as negative intensities cannot occur in MS, the latter case is of purely theoretical relevance. *S-values* less than 1 (no, negligible, or negative correlation) are set to 1:

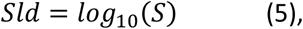

As equation (5) illustrates, *S-values* are subsequently log_10_ transformed (*Sld*). A *Sld* value of 3 indicates that peak tables *x* and *y* are identical, whereas a value of 0 points to the (almost) complete absence of positive correlations.

It is important to note at this place that a direct comparison between MicrobeMS *Sld* scores and Bruker’s score values is not attainable. The latter are calculated by means of alternative preprocessing methods and result from a significantly more complex computational procedure [43] that cannot be vectorized using MATLAB (see the following section for further details). Observations to date have indicated that, on average, MicrobeMS’ *Sld* values from identical data are marginally higher than the respective MBT scores, with an average discrepancy of approximately 0.3 units. This pertains to standard workflows and default parameter settings in both programs. Unlike Bruker Daltonics, this author will abstain from offering a general recommendation regarding generic *Sld* levels that should be reached to indicate highly probable, secure, or probable identification at the genus or species level, respectively.

One significant advantage of the MicrobeMS methodology described is that, as a higher-level programming language, MATLAB can efficiently perform the calculation of a complete set of spectral similarity values. However, this requires the use of a methodology referred to by The MathWorks as *vectorization*, a method that relies on matrix and vector operations instead of loop-based scalar-oriented programming. On the one hand, this configuration imposes greater demands on the RAM equipment, yet it concomitantly offers substantial advantages in terms of processing speed. For instance, the calculation of 800,000 correlation coefficients between, for example, 2,000 database peak tables and 400 tables from experimental microbial mass spectra is completed in less than one second on standard PC hardware from 2026.

The advanced computing capabilities of MicrobeMS enabled the incorporation of a brute force correction method for calibration errors. In this approach, the calibration constants of experimental mass spectra can be varied according to a user-adjustable scheme and involves simultaneously testing the entire array of peak tables derived from the calibration-varied mass spectra against database entries. From the *Sld* values obtained, either the highest, or mean values of *n* highest *Sld* values are selected for each combination of calibration-varied test entries and a given database entry. The procedure described proved to be effective in minimizing errors that may arise due to unavoidable calibration inaccuracies. Despite the method’s computational demands, the implementation of vectorized MATLAB code was helpful to prevent extended execution times.

#### Interpretation of score ranking lists

The method described in the following section was inspired by a publication by Harju et al. (2017), co-authored by Thomas Maier and Markus Kostrzewa from Bruker Daltonics [44]. Score ranking lists are not always unambiguously interpretable, particularly in the following cases: a) the scores (*Sld* values) of entries of closely related species are similar, b) The rank lists comprise disordered arrangements of two or more representatives of different, often closely related species, or c) a mix of a) and b). The idea outlined by Harju et al. aims at objectively evaluating rank lists in cases where a clear interpretation is difficult. To this end, the species assignment of each of the listed databases entry, as well as their list position and score values, are automatically analyzed and summarized as a *so-called* rank list score for each of the species involved. The result of such a rank list analysis, the logarithmic rank list score Sld_rl,j_, is calculated using equations (6) through (9) which are indicated below:

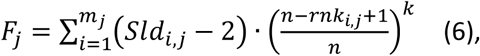

In equation (6) *F_j_* denotes an absolute rank list parameter for species *j*; *rnk_i,j_* and *Sld_i,j_* are the rank and the corresponding log score of the *i-th* strain entry of the *j-th* species present in the given rank list, respectively. Furthermore, *n* equals the total number of entries in the rank list, usually 20, while *k* denotes an exponent (*k* = 9.0). Log scores below 2 are set to 2; furthermore, *mj* equals the number of strains from species *j* present in the database of MALDI-ToF mass peak tables.

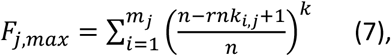

In equation (7) *F_j,max_* is the maximum theoretical value, *F_j_* can reach. This is achieved when all strains of species *j* are listed on top positions and all log scores equal 3.

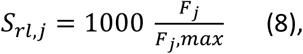

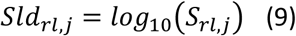

*Sr_l,j_* is the rank list score of species *j*, with 0 ≤ *Srl,j* ≤ 1000. *Sld_rl,j_* denotes the respective log score. Logarithmic rank list scores can vary between values of 0 and 3.

Rank analyses are considered reliable when the database contains more than five, preferably over eight, strains of the species in question. If this condition is not met, the results should be interpreted with greater caution. This is especially important when database spectra are available from only a few strains, or even a single strain of a given species. However, equations (6) through (9) and their parameter settings ensure that the top entry in the score ranking list cannot be overturned from its position if this entry is represented in the database by only one strain.

In addition to distances based on PPMCCs, MicrobeMS features a variety of alternative distance calculation options. These include Euclidean distances, Pareto-scaled distances, as well as covariance-based distances. It is evident that the selection of these distance metrics necessitates the utilization of equations other than those listed in (2) through (9). For more information, see the MicrobeMS Wiki [33].

For identification of bacteria by means of MALDI-ToF MS, i.e. mass spectrum databases and interspectral distances, MicrobeMS offers two distinct workflows. A standardized workflow requiring only minimal user interaction allows semi-automated identification analysis. The second option is characterized by its ability to accommodate a wider range of parameter selection, making it particularly well suited for users with a more extensive experiential background. More information is given below, see section 6.3., and in the MicrobeMS Wiki [33].

## 5. DESCRIPTION OF THE TEST DATA SETS

### 5.1 The HPB test panel

The HPB test panel, also known as the HPB test panel, was compiled in 2014 in the context of an external quality assurance exercise (EQAE) to identify HPB using MALDI-ToF MS. The exercise was conducted as part of the EU-funded project, “*Quality Assurance Exercises and Networking on the Detection of Highly Infectious Pathogens*” (QUANDHIP). Its aim was to build up a consortium of specialized European laboratories to guarantee the exchange of diagnostic strategies to support a coordinated response to outbreaks of highly pathogenic infectious agents. This EQAE was designed as a blinded laboratory comparison study and involved ten different bacterial isolates, including five HPB and five non-HPB test strains. Some of these strains were taxonomically closely related (cf. Table 1). For example, the EQAE test strains from *Burkholderia mallei*, *B. pseudomallei*, and *B. thailandensis* are members of the *B. pseudomallei* group, making differentiation between these strains difficult. Eleven European QUANDHIP project partners, including RKI, participated in the exercise. While the RKI unit *Highly Pathogenic Bacteria* (ZBS2) organized the interlaboratory test, and was responsible for strain selection, sample preparation, inactivation, and shipment, the *Proteomics and Spectroscopy* unit (ZBS6) was involved as a regular ring trial participant. Consequently, microbial samples were provided blindly to ZBS6. Table 1 provides an overview of the taxonomic status and initial concentration of bacterial suspensions, as well as the quality test results obtained by the recent reanalysis.

**Table 1.** The HPB test panel spectra were acquired by the RKI partner as part of a European inter-laboratory ring trial. Prior to shipment and MALDI-ToF-compatible sample preparation, the bacterial material was inactivated using high-dose γ-irradiation (30 kGy) [9].

| Sample number | Genus / Species / Strain | Number of spectra | Concentration (cfu/mL) * | Quality test results |
| --- | --- | --- | --- | --- |
| 1 | *Burkholderia pseudomallei* A101-10 | 2 | $1.1 \times 10^9$ | 71.8 / 66.1 |
| 2 | *Francisella tularensis* ssp. *holarctica* Ft 32 | 2 | $1.7 \times 10^{10}$ | 68.6 / 63.6 |
| 3 | *Brucella canis* A183-5 | 2 | $1.9 \times 10^{10}$ | 71.1 / 66.1 |
| 4 | *Bacillus anthracis* AMES | 3 | $6.4 \times 10^7$ | 65.4 / 46.1 / 68.2 |
| 5 | *Ochrobactrum (Brucella) anthropi* A148-11 | 2 | $2.0 \times 10^{10}$ | 70.7 / 70.0 |
| 6 | *Yersinia pseudotuberculosis* type III | 2 | $1.3 \times 10^9$ | 75.4 / 72.5 |
| 7 | *Burkholderia mallei* A106-3 | 2 | $1.0 \times 10^9$ | 72.5 / 73.2 |
| 8 | *Burkholderia thailandensis* E125 | 3 | $5.6 \times 10^{10}$ | 60.4 / 73.6 / 67.1 |
| 9 | *Yersinia pestis* A106-2 | 3 | $1.3 \times 10^7$ | 47.1 / 53.2 / 60.7 |
| 10 | *Bacillus thuringiensis* DSM 350 | 3 | $8.6 \times 10^8$ | 58.2 / 57.1 / 45.0 |

\* cfu – colony forming units

The TFA method [10] was used to prepare samples from the *a priori* unknown test strains. MALDI-ToF mass spectra were subsequently recorded from these samples using an *autofleX* I mass spectrometer from Bruker Daltonics. More specific details on sample preparation, data acquisition and test results are available from Lasch et al., 2025 [9]. The HPB test dataset is available from the MicrobeMS Wiki.

### 5.2 The ZENODO database

MicrobeMS was designed, *inter alia*, as a tool for the development and management of MALDI-ToF MS spectrum databases. These databases focus on highly pathogenic bacterial species belonging to the genera *Bacillus*, *Brucella*, *Burkholderia*, *Francisella* and *Yersinia*, and have been made available to the public free of charge at irregular intervals since 2016 on the ZENODO website hosted by CERN. The most recent version, v. 4.2, was released in June 2023 and contains a total of 11,055 MALDI-ToF mass spectra from 1,601 bacterial strains of biosafety levels 1–3 (BSL1–3) [17]. The spectrum data in the original *BrukerRaw* MALDI-ToF MS file format, complete metadata and a database peak lists (MicrobeMS *.pkf* and Bruker’s .*btmsp* file format) are available to download from the ZENODO website. A comprehensive description of the RKI database and its contents was published in 2025 [16]. The publication of the next database version, v. 5.0 is planned for the near future.

### 5.3 The RKI in silico MALDI-ToF MS database

The RKI *in silico* database has been under active development for several years and is used for diagnosing rare and special bacterial pathogens. The latest version of this database contains altogether 17,850 pseudo-MALDI spectra which are derived from UniprotKB, i.e. ultimately from bacterial genomes. The 2024 version of this *in silico* database is still very experimental in nature; the identification results based on it do not achieve the accuracy of results based on spectrum databases that rely on mass spectrometric measurements of bacterial samples. The *Bacteria-May-30-2024-sprot+trembl-50724-standard.pkf* database (*.pkf* format) contains 80 pseudo-peaks per *in silico* pseudo-spectrum. It can be used by MicrobeMS without restrictions or technical limitations in the same way as experimental databases. This database is available for download on the MicrobeMS Wiki website [33] but is not intended for diagnostic purposes. It is provided exclusively for testing and scientific applications.

A publication with more detailed information on database creation and an improved version of the *in silico* database is currently being prepared.

## 6. WORKFLOW EXAMPLES

### 6.1. Quality Tests carried out on the HPB Test Panel

In MicrobeMS, performing a QT is quite straightforward: First, previously loaded mass spectra are selected in the list box labelled with ‘*MicrobeMS spectra IDs*’ (top right position of the main gui, see Figure 1). After selecting the ‘*Quality Test*’ option, a dialog box titled ‘*Quality Test*’ will appear (see Figure 4A), which allows modifying default parameters of the QT routine. After clicking the ‘*Quality Test*’ button, the QTs of all the selected spectra are executed one after the other, based on the input parameters chosen. A progress bar illustrates the percentage of completed computations. Once all calculations have been completed, the buttons that were previously greyed out can be used to either save the QT results as an Excel table (button ‘*Store Report*’) or display them in HTML format via the system web browser (button ‘*Show Report*’). The latter displays the results in a color-coded format using a predefined traffic light scheme of red, yellow or green (cf. Figure 4B).

Executing the QT using the HPB test set yielded satisfactory results overall. With an average QT result of 64.3 (range 45.0–75.4) overall, sufficiently good results could be achieved. However, it was noticeable that low pathogen concentrations resulted in only moderate QT results (cf. lines 4, 9 and 10 of Table 1 for *B. anthracis*, *Y. pestis* and *B. thuringiensis*, respectively). Below-average QT can negatively affect the correctness of identification results, especially when genetically very closely related species are tested together. One such example is the bioterrorism-relevant anthrax pathogen *B. anthracis*, which exhibits only minor genetic and, consequently, only subtle MALDI-ToF MS differences compared to *B. thuringiensis*. Both species belong to the *B. cereus* group, which some authors consider to be a single species [45–47]. A similar situation exists for *Y. pestis* and *Y. pseudotuberculosis*, both of which are also part of the HPB test panel: *Y. pestis*, the plague pathogen, is considered a recently evolved clone of *Y. pseudotuberculosis* [48–50]. Nevertheless, due to the very different clinical courses, and the severity of infections caused by *Y. pestis*, this pathogen is continued to be classified as a separate species.

Therefore, to exploit the full potential of the MALDI-ToF MS-based method in these and similar cases, and to achieve accurate identification results, it is important to record high-quality mass spectra. In the case of low-concentration HPB samples, acceptable-quality mass spectra were obtained through manual mode measurements, whereby weak MS signals were summed up over an extended period of time. Objective tools for assessing spectrum quality can assist experimenters in such cases.

The detailed results of the QT test on the HPB test set can be found in the Supporting Information in the form of an HTML-formatted quality test report (*report-quality-07-Jan-2026-11-53-47.html, see SI*).

### 6.2. Unsupervised hierarchical cluster analysis (UHCA) of the HPB test panel

When exploring unknown data sets, the UHCA approach is a good starting point. This approach is relatively simple and requires no prior knowledge of internal data structures. For hierarchical clustering, the MALDI-ToF mass spectra must be adequately pre-processed beforehand. In the given UHCA application example, this involves smoothing, baseline correction, and normalization. In MicrobeMS this can be preferably done by applying predefined standard parameters, followed by extracting peak lists from the pretreated mass spectra. As illustrated in Figure 6, peak extraction was carried out in this example by using predefined standard peak pick parameters, which include, among others, a setting of 30 peaks per spectrum within the m/z range between 2000 and 13,000 Th units. Once the peak lists have been generated, they can be either saved as *.pkf* files via the ‘*save*’ option (see file pull-down menu) and then loaded by pressing the ‘*load*’ button available in the dialog box ‘*UHCA of mass peak tables*’ that opens after pressing a button labeled as ‘*hierar. clustering*’ in the ‘*ANALYSIS*’ tab (cf. Figure 6). Alternatively, peak lists can be exported by marking spectra in the *’MicrobeMS Spectra ID’s*’ list box and clicking then the ‘*new*’ button in the ‘*PEAK DATABASE*’ tab. The next step is to open the UHCA dialog box using the ‘*hierar. clustering*’ button. Peak lists saved in this way will then be available for UHCA. In the UHCA dialog box the most critical UHCA settings pertain to the methods for calculating interspectral distances and the clustering algorithm. Furthermore, MicrobeMS allows users to choose between intensity data or barcoded data as inputs to obtain distances. In the latter case, the ‘*use weightings*’ option should be deactivated. A full description of UHCA’s functionality goes beyond the scope of this publication but is available from the MicrobeMS Wiki [33].

Figure 6 exemplifies the dendrogram created with MicrobeMS based on extracted peak tables from the HPB test data set. For preprocessing, the original MALDI-ToF mass spectra were first loaded in the *BrukerRaw* format and then processed using the following sequence of routines: Smoothing (Savitzky-Golay smoothing filter with 21 smoothing points), asymmetric least square baseline correction, normalization, in both instances with the default parameter set. Subsequently, 30 peaks were extracted in the m/z range from 2000 to 13,000 per spectrum. For UHCA, D-values (correlation) were selected for distance calculation and Ward’s method was chosen as the cluster algorithm. Additional parameters were inactive settings for ‘*use weightings*’ and a relative width of m/z intervals of 800 ppm.

The dendrogram generated by means of these settings from the HPB test data demonstrates that spectra of the technical replicates invariably form separate strain clusters, despite the very high degrees of taxonomic relatedness they may express. However, the fusion levels within these strain-specific clusters differ. Clusters of bacterial strains for which only suspensions with low cell concentrations were available show a higher degree of heterogeneity than clusters from bacterial suspensions of higher concentrations (cf. Figure 6, Clusters of *B. anthracis*, *B. thuringiensis* and *Y. pestis* and Table 1). This can certainly be attributed to the reduced quality of some of these spectra from the species. Figure 6 also illustrates that strain-specific clusters of closely related species form higher-order clusters. For example, spectra of both *Bacillus* species, *Bacillus anthracis* and *B. thuringiensis*, can be found in the same clade. The same holds true for representatives of *Burkholderia* and *Yersinia*: Three representatives of the *Burkholderia pseudomallei* group (*B. mallei*, *B. thailandensis* and *B. pseudomallei*) form a higher-level clade, similarly the two *Yersinia* species, *Y. pestis* and *Y. pseudotuberculosis* (cf. Figure 6). Furthermore, the dendrogram suggests the similarity of peak patterns of *Brucella canis* and *Ochrobactrum anthropii*. Although the taxonomic status of *Ochrobactrum* (*Brucella*) *anthropii* is currently the subject of considerable controversy in the literature, the close relationship between these two species is widely undisputed [51, 52]. In summary, the selected example nicely illustrates the usefulness of UHCA for analyzing bacterial mass spectra. It is re-emphasized, however, that such type of analysis requires controlling the spectrum quality and assuring adequate preprocessing of the microbial mass spectra tested.

### 6.3. Identification analysis using MALDI-ToF mass spectra of the HPB test panel

There are two ways to perform identification analyses in MicrobeMS. The first option is intended for experienced users because all preprocessing, peak detection, and identification analysis steps must be performed manually in a step-by-step process. This workflow requires selecting a large number of specific parameters for each step; each parameter affects the identification results. While this approach allows for a highly flexible response to specific analysis requirements, it requires the operator to have specific knowledge and special experience. A detailed description of the manual workflow procedure can be found in the MicrobeMS Wiki [33].

The second approach to identification analyses requires significantly less effort and experience because both the sequence and the parameters of the individual analysis steps are predefined. The following describes the process of an automated standard identification analysis using the HPB test dataset.

After loading and selecting the MALDI-ToF mass spectra to be analyzed, click the ‘*standard ID*’ button in the ‘*ANALYSIS*’ tab. The selected spectra will then be subjected to QT and pretreated according to the pre-defined specifications in the file *microbems.opt*. After peak extraction, a dialog box titled ‘*Identification analysis based on interspectral distances*’ will open. There, one can change the parameters for peak list comparison, but this is only recommended for experienced users.

If the name and path of the *.pkf* database file to be used have been indicated correctly in the *microbems.opt* configuration file, the corresponding database will be loaded automatically when the dialog box opens. In such cases, users can initiate the identification analysis by selecting the ‘compare’ button (see Figure 7A). When testing experimental peak tables against corresponding database entries using correlation or distance methods, it is essential that the number of peaks in the experimental tables is identical to the number of peaks in the database entries. For this reason, there are different versions of the RKI databases with 30, 45, or 78 peaks per entry. These can be downloaded in the *.pkf* format from the ZENODO website or obtained from the author of this study. While peak lists with 30 peaks have proven advantageous for identification analyses with experimental databases, peak lists with 78 entries were found to be effective for analyses with the *in silico* database.

**Figure 7.**
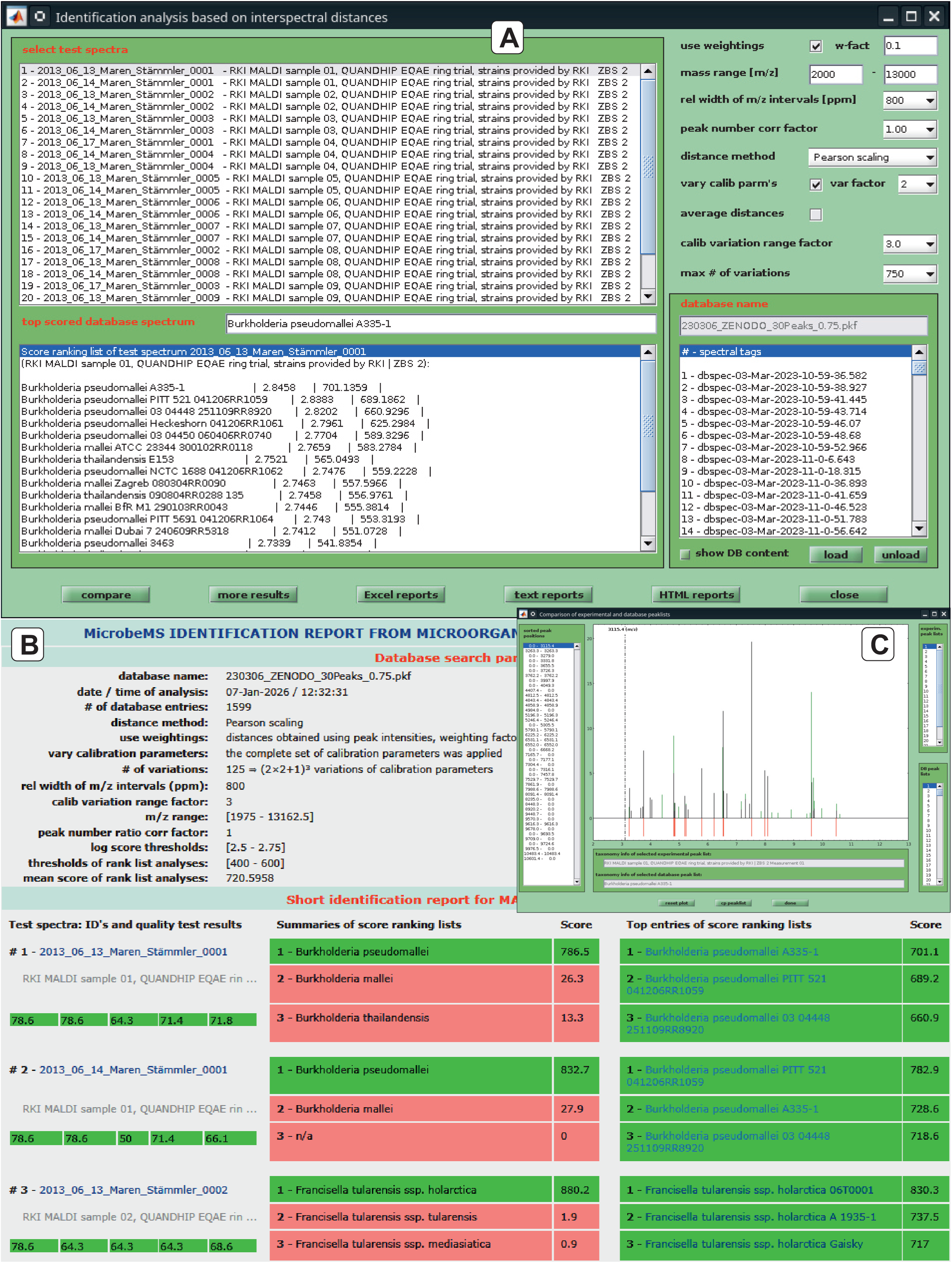
Identification analysis of MALDI-ToF mass spectra from the HPB test panel (see section 5.1) **A.** User dialog box ‘*Identification analysis based on inter-spectral distances*’. **B.** Snapshot of a pop-up window allowing visual comparison of experimental and database peak lists. **C.** Identification report (HTML formatted) with results of MALDI-ToF MS identification analysis. The top section lists important details of the analysis: name and contents of the peak list database used, parameters of identification analysis, etc. The lower left column provides test spectra info such as the spectrum ID, taxonomic data, and QT results. Taxonomic information of three topmost ranked database spectra and the respective scores are given in the right column. Note the traffic light scheme. A summary of the corresponding score ranking list analysis is given by the lower middle column.

Once the analyses are complete, the identification results can be output by pressing the ‘*HTML reports*’ button, for example (see Figure 7B).

The HTML document (*report-maldi-07-Jan-2026-12-32-36.html, see SI*) contained in the supporting information details the results of the identification analyses with spectra from the HPB test panel. As the results overview show, the correct taxonomic identification at the genus, species, and subspecies levels was found for most samples, with only one exception. In addition to this, the correct subspecies information is provided for sample no. 2 (*Francisella tularensis* subsp. *holarctica*, see Table 1 and *report-maldi-07-Jan-2026-12-32-36.html* (see SI). Out of 24 cases and despite the high degree of relatedness among samples within the HPB test panel, only one incorrect assignment occurred (cf. spectrum #21, *Y. pestis*, misidentified as *Y. similis*). As a member of the *Yersinia pseudotuberculosis* complex, *Y. similis* is genetically highly related to *Y. pseudotuberculosis* and *Y. pestis*. [53, 54] Since high genetic similarity is usually accompanied by similar peak patterns, experts should perform a detailed manual peak analysis in these and similar cases before making a diagnosis. A typical MALDI feature for strains of *Y. pestis* carrying one of the two *Y. pestis*-specific plasmids (pPCP1) is an intense mass peak at m/z 3065 [13, 15]. In the present example, visual inspection of the MALDI-ToF mass spectra of sample #9 confirmed the presence of an intense MS signal at this m/z position. Therefore, despite the misidentification of spectrum #21 by a peak list correlation-based approach, manual follow-up examination suggested identification of sample #9 as *Y. pestis*.

In MicrobeMS the results of identification analyses can be saved as plain text or in an MS Excel data format. The latter option considerably helps to streamline subsequent statistical analysis of identification results, for instance, for monitoring analyses of the complete RKI MALDI-ToF MS spectrum database.

## 7. FURTHER FUNCTIONALITIES

For the sake of completeness, this section will briefly mention some additional functionalities of MicrobeMS. Due to space limitations, these will only be referenced; no comprehensive description will be provided.

Aside from the correlation, or distance-based options for identifying pathogenic microorganisms, MicrobeMS offers also an interfaces to ANN software (NeuroDeveloper [21]) and a routine that allows experimental mass spectra to be queried against an *in silico* calculated MALDI database, derived from UniprotKB data [55].

The ANN interface was originally developed because highly pathogenic BSL-3 bacteria, such as *B. anthracis* and *Y. pestis*, are closely related to some BSL-2 pathogens, such as *B. cereus* sensu stricto (*B. cereus* s.s.) and *Y. pseudotuberculosis*, respectively, and therefore only minor spectrometric differences can be observed between the BSL-3 and BSL-2 representatives in either example [24]. This makes it difficult to identify the pathogens by standard spectral distance methods and was a key motivation to develop the ANN-based identification function, and MicrobeMS as a whole. High standards in cultivation, sample preparation, and data acquisition are essential for successfully applying ANNs to accurately identify the aforementioned, or to discriminate other BSL-3 representatives (e.g. *Burkholderia mallei* vs. *Burkholderia pseudomallei*). Additionally, a sufficiently large number of well-characterized microbial isolates of BSL-3 pathogens and related species must be available from which a sufficient number of biological and technical replicate spectra should be recorded. The ANN identification function has been used successfully in the early years of MicrobeMS, [12, 13] but it has not been actively developed further in recent years.

The function ‘*average spectra*’ can be used to obtain mean MALDI-ToF mass spectra. This function generates new spectrum files and is helpful to increase the spectral signal-to-noise ratio (SNR). Another possible purpose of averaging spectra is to reduce unwanted spectral variability, e.g. when measuring spatially inhomogeneous samples. No weighting by the spectra’ individual laser shots is performed. Furthermore, executing the ‘*average spectra*’ function does not include preprocessing or calculating peak tables.

The ‘*peak frequency test*’ function can be employed to systematically obtain and to visualize the frequency of mass peaks from ensembles of mass spectra. For this purpose, mass spectra are first discretized into m/z segments (bins) of a defined width which is controlled by a ‘*m/z tolerance*’ value. This program-wide parameter is defined in relative ppm units which means that bins at low m/z positions are narrow and broad in the high m/z range. Then, peak abundance values are obtained for each of the m/z segments in a systematic manner. The ‘*peak frequency test*’ function is useful for biomarker screening and requires pre-processed spectra with peak tables. The output of the peak frequency test series is a table listing the ranked absolute and relative peak frequencies. Additionally, the relative peak frequency values are plotted as a curve against the m/z segment positions.

Two-sample *t-tests* and *Wilcoxon rank-sum tests* are two further potentially useful methods for biomarker peak screening. Both tests require labeled subsets of spectra and peak lists thereof, preferentially obtained from the pre-processed spectra. Again, tests are systematically performed using peak data extracted consecutively from the previously mentioned m/z bins within a spectrum range of interest. The results of these tests are provided at the program’s command line and as curves showing the dependence either of the H(0) hypothesis test output (boolean), p-values, or alternative test parameters as functions of the bins’ m/z center positions. A more detailed description of these functionalities can be found on the MicrobeMS Wiki [33].

## 8. INDEPENDENT TESTING

### 8.1. Matthias Mailander

Developer at Lablicate GmbH, the driving force to create OpenChrom®, an Open Source multi-vendor cross-platform chromatography data system. Martin-Luther-King-Platz 6, 20146 Hamburg, Germany.

Every functionality of MicrobeMS 0.92 that was accessible via the graphical user interface was systematically tested except NeuroDeveloper integration and LC-MS *in-silico* spectra. MicrobeMS loads MALDI-ToF MS instrument vendor data flawlessly, contains an effective preprocessing pipeline, and its identification as well as classification with built-in quality control tests is robust. The software is extensively documented, including its file formats to produce FAIR data.

### 8.1. Jorg Rau

The software was independently tested by Dr. Jorg Rau, Chemical and Veterinary Analysis Agency Stuttgart, CVUAS, Germany.

MicrobeMS v. 0.92 was installed and tested on a Windows 10 PC. The software was tested in a real laboratory environment, where all Bruker MALDI-Biotyper software programs were running in parallel. Before installing MicrobeMS 0.92, it was important for us to keep the MATLAB environment compatible with the MALDI Biotyper Compass Explorer 4.1 (Bruker Daltonik, GmbH, Bremen). This can be ensured by installing MATLAB R2023b 1.0 and MATLAB v75 in parallel. The accompanying comprehensive and detailed manual in wiki format (https://wiki-ms.microbe-ms.com/) is particularly helpful and necessary.

The interface of MicrobeMS v. 0.92 is simple and clearly organized. Correctly loading your own spectra for analysis only makes sense once the metadata has been converted to the format specified by MicrobeMS. An auxiliary table is provided in the Wiki for this purpose. The metadata record for ‘MALDI-Fields.xls’ can be filtered out almost completely from the MALDI-UP catalog (https://maldi-up.ua-bw.de) [56] and transferred. In the medium term, the outdated Excel file format (.xls) could prove problematic in the application. Equipped with this metadata, individual spectra can be used in various ways for identification. Our test spectrum examples came from an independent validation study for the *Corynebacterium diphtheriae* species complex (Rau et al., 2026). The report options generated for identification with MicrobeMS (text, Excel, html) are easy to interpret. The identification analysis based on interspectral distances (MBT-like ID) * provides us, as Bruker users, with the familiar score values. The success of identification naturally depends on the database used. The calculations performed with MicrobeMS are fast and reliable, even for more complex cluster analysis. The software from the world’s leading MALDI-ToF MS manufacturers, Bruker and Shimadzu, lacks compatibility between the formats of the MALDI-ToF mass spectra. This has prevented the combination of data until now. MicrobeMS, as a freeware analytic tool, could be helpful here.

* Comment from the author: This relatively new feature has not been described in the current manuscript.

## Supporting information

QT report of the HPB data set

Identification report of the HPB data set

## Abbreviations

EQAE: external quality assurance exercise
HPB: highly pathogenic bacteria
LC-MS: liquid chromatography - mass spectrometry
MALDI-ToF: matrix-assisted laser desorption/ionization - time–of–flight
MBT: MALDI Biotyper
MSP: main spectrum profile or main spectral projection
m/z: mass-to-charge ratio
HCCA: α-cyano-4-hydroxycinnamic acid
MS: mass spectrometry
PPMCC: Pearson’s product-moment correlation coefficient
QT: quality test
RKI: Robert Koch Institute
TFA: trifluoroacetic acid
ZBS: Centre for Biological Threats and Special Pathogens

## 10. CODE AVAILABILITY

The MATLAB-based software MicrobeMS used for this work is freely available as a pcode version for Windows or LINUX, or as a standalone version (Windows only) from https://wiki-ms.microbe-ms.com.

## 11. ARTIFICIAL INTELLIGENCE (AI) USAGE STATEMENT

For the creation of this publication, AI was solely employed for linguistic revision of the manuscript text. For this purpose, the commercial version of DeepL (https://www.deepl.com) was used. No other AI methods or tools were employed.

## 12. ACKNOWLEDGEMENTS

I thank Nico Fischer, Elke Westhauser, Charlyn John, Jeremias Meiß, Maren Stammler and Andy Schneider (all RKI, Berlin) for carefully performing microbial sample preparation and MALDI-ToF MS measurements. Furthermore, the author is grateful to Dr. Joerg Doellinger (RKI), Dr. Alejandra Bosch (CINDEFI, La Plata, Argentina), Dr. Marcel Erhardt (former Anagnostec) and Dr. Thomas Maier (Bruker Daltonics) for fruitful discussions and support.

