## Supplementary material for "MicrobeMS - A MATLAB Toolbox for Microbial Identification Based on Mass Spectrometry": QT report of the HPB data set

### MicrobeMS QUALITY TEST REPORT OF MICROORGANISM MASS SPECTRA - OVERVIEW

#### Quality test parameters

| | |
|---|---|
| **number of spectra tested:** | 24 |
| **sequence of preprocessing steps:** | AsLS → normalization → peak detection |
| **type of baseline correction:** | baseline correction by asymmetric least squares (AsLS) |
| **asymmetry factor (AsLS):** | 0.0001 |
| **lambda factor (AsLS):** | 100000 |
| **number of iterations (AsLS):** | 8 |
| **type of normalization:** | offset correction and vector-norm of mass spectra such that their standard deviation finally equals 1000 |
| **parameters of peak detection:** | [2000 13000 300 12000 7000 1.5 7.5 800 1 40] |
| **peak search m/z region start value:** | 3000 |
| **peak search m/z region end value:** | 13000 |
| **delta m/z, parameter of the intensity threshold function:** | 7000 |
| **delta counts, parameter of the intensity threshold function:** | 1.5 |
| **m/z slope, parameter of the intensity threshold function:** | 7.5 |
| **resolution of the m/z vector (relative units):** | 800 |
| **fix number of peaks (boolean):** | 0 |
| **number of peaks per spectrum:** | n/a |
| **number of bins of quality test parameters:** | 15 |
| **edges of noise bins:** | [0.4 0.464 0.541 0.633 0.743 0.876 1.035 1.226 1.455 1.73 2.059 2.455 2.93 3.5] |
| **quality parameter 'noise level' - weighting factor:** | 0.3 |
| **edges of baseline error bins:** | [1 1.451 2.037 2.798 3.789 5.076 6.749 8.925 11.753 15.429 20.209 26.422 34.499 45] |
| **quality parameter 'baseline level' - weighting factor:** | 0.15 |
| **edges of bins 'number of peaks':** | [8 10 12 14 16 18 21 24 28 31 35 40 45 50] |
| **quality parameter 'number of peaks' - weighting factor:** | 0.4 |
| **edges of resolving power bins:** | [250 285 323 365 411 462 517 579 646 721 802 892 991 1100] |
| **quality parameter 'resolving power level' - weighting factor:** | 0.15 |
| **general qt test criteria ('red,'yellow','green'):** | [30 45] |

#### Short quality test report for MALDI-ToF mass spectra

| ID's of test spectra | Noise test result | Baseline test result | 'Number of peaks' test result | resolving power test result | Overall quality test result |
|---|---|---|---|---|---|
| **# 1** - 2013_06_13_Maren_Stämmler_0001<br>RKI MALDI sample 01, QUANDHIP EQAE ring trial, strains ... | 78.6 | 78.6 | 64.3 | 71.4 | **71.8** |
| **# 2** - 2013_06_14_Maren_Stämmler_0001<br>RKI MALDI sample 01, QUANDHIP EQAE ring trial, strains ... | 78.6 | 78.6 | 50 | 71.4 | **66.1** |
| **# 3** - 2013_06_13_Maren_Stämmler_0002<br>RKI MALDI sample 02, QUANDHIP EQAE ring trial, strains ... | 78.6 | 64.3 | 64.3 | 64.3 | **68.6** |
| **# 4** - 2013_06_14_Maren_Stämmler_0002<br>RKI MALDI sample 02, QUANDHIP EQAE ring trial, strains ... | 71.4 | 64.3 | 57.1 | 64.3 | **63.6** |
| **# 5** - 2013_06_13_Maren_Stämmler_0003<br>RKI MALDI sample 03, QUANDHIP EQAE ring trial, strains ... | 85.7 | 78.6 | 57.1 | 71.4 | **71.1** |
| **# 6** - 2013_06_14_Maren_Stämmler_0003<br>RKI MALDI sample 03, QUANDHIP EQAE ring trial, strains ... | 78.6 | 71.4 | 50 | 78.6 | **66.1** |
| **# 7** - 2013_06_17_Maren_Stämmler_0001<br>RKI MALDI sample 04, QUANDHIP EQAE ring trial, strains ... | 78.6 | 50 | 64.3 | 57.1 | **65.4** |
| **# 8** - 2013_06_14_Maren_Stämmler_0004<br>RKI MALDI sample 04, QUANDHIP EQAE ring trial, strains ... | 57.1 | 28.6 | 42.9 | 50 | **46.1** |
| **# 9** - 2013_06_13_Maren_Stämmler_0004<br>RKI MALDI sample 04, QUANDHIP EQAE ring trial, strains ... | 78.6 | 50 | 71.4 | 57.1 | **68.2** |
| **# 10** - 2013_06_13_Maren_Stämmler_0005<br>RKI MALDI sample 05, QUANDHIP EQAE ring trial, strains ... | 78.6 | 71.4 | 64.3 | 71.4 | **70.7** |
| **# 11** - 2013_06_14_Maren_Stämmler_0005<br>RKI MALDI sample 05, QUANDHIP EQAE ring trial, strains ... | 71.4 | 50 | 78.6 | 64.3 | **70** |
| **# 12** - 2013_06_13_Maren_Stämmler_0006<br>RKI MALDI sample 06, QUANDHIP EQAE ring trial, strains ... | 78.6 | 71.4 | 78.6 | 64.3 | **75.4** |
| **# 13** - 2013_06_14_Maren_Stämmler_0006<br>RKI MALDI sample 06, QUANDHIP EQAE ring trial, strains ... | 78.6 | 71.4 | 71.4 | 64.3 | **72.5** |
| **# 14** - 2013_06_13_Maren_Stämmler_0007<br>RKI MALDI sample 07, QUANDHIP EQAE ring trial, strains ... | 78.6 | 64.3 | 71.4 | 71.4 | **72.5** |
| **# 15** - 2013_06_14_Maren_Stämmler_0007<br>RKI MALDI sample 07, QUANDHIP EQAE ring trial, strains ... | 71.4 | 64.3 | 78.6 | 71.4 | **73.2** |
| **# 16** - 2013_06_17_Maren_Stämmler_0002<br>RKI MALDI sample 08, QUANDHIP EQAE ring trial, strains ... | 71.4 | 50 | 57.1 | 57.1 | **60.4** |
| **# 17** - 2013_06_13_Maren_Stämmler_0008<br>RKI MALDI sample 08, QUANDHIP EQAE ring trial, strains ... | 78.6 | 71.4 | 71.4 | 71.4 | **73.6** |

| | | | | | |
|---|---|---|---|---|---|
| **# 18** - 2013_06_14_Maren_Stämmler_0008<br>RKI MALDI sample 08, QUANDHIP EQAE ring trial, strains ... | 71.4 | 50 | 71.4 | 64.3 | **67.1** |
| **# 19** - 2013_06_17_Maren_Stämmler_0003<br>RKI MALDI sample 09, QUANDHIP EQAE ring trial, strains ... | 78.6 | 57.1 | 21.4 | 42.9 | **47.1** |
| **# 20** - 2013_06_13_Maren_Stämmler_0009<br>RKI MALDI sample 09, QUANDHIP EQAE ring trial, strains ... | 78.6 | 78.6 | 28.6 | 42.9 | **53.2** |
| **# 21** - 2013_06_14_Maren_Stämmler_0009<br>RKI MALDI sample 09, QUANDHIP EQAE ring trial, strains ... | 78.6 | 64.3 | 50 | 50 | **60.7** |
| **# 22** - 2013_06_17_Maren_Stämmler_0004<br>RKI MALDI sample 10, QUANDHIP EQAE ring trial, strains ... | 78.6 | 28.6 | 57.1 | 50 | **58.2** |
| **# 23** - 2013_06_13_Maren_Stämmler_0010<br>RKI MALDI sample 10, QUANDHIP EQAE ring trial, strains ... | 71.4 | 35.7 | 57.1 | 50 | **57.1** |
| **# 24** - 2013_06_14_Maren_Stämmler_0010<br>RKI MALDI sample 10, QUANDHIP EQAE ring trial, strains ... | 71.4 | 7.1 | 42.9 | 35.7 | **45** |

---

#### # 1 - QUALITY TEST REPORT FOR MALDI-TOF MS SPECTRUM *'2013_06_13_Maren_Stämmler_0001'*

##### Metadata of actual MALDI-ToF test spectrum

| | |
|---|---|
| **genus / species / strain:** | RKI MALDI sample 01, QUANDHIP EQAE ring trial, strains provided by RKI \| ZBS 2 |
| **file id:** | 2013_06_13_Maren_Stämmler_0001 |
| **type:** | Measurement 01 |
| **Bruker ID:** | 732DF236-D7F4-4EE7-99BF-FD7E225F3A6F |
| **NCBI ID (primary):** | 28450 |
| **NCBI ID (secondary):** | 28450 |
| **growth time:** | Optimal growth time between 24 - 72h |
| **growth temperature:** | 37°C |
| **growth conditions:** | Optimal aerobic or microaerophilic conditions |
| **growth medium:** | Columbia blood agar (Oxoid), 2nd passage on TSA or Caso agar, harvested by the 2nd passage |
| **sample treatment:** | Sample mixed with 20 mkL 100Â§ TFA, final TFA conc. approx. 80 perc.; approx. 30 Min treatment time; diluted 1:10 (vol); mixed with 1:1 HCCA\|TA2(A) |
| **spores:** | No |
| **concentration:** | Pellet produced by centrifugation (1 x 5 Min, 15,000 rpm) of a 500 mkL cell suspension, gamma ray irradiated (30 kGy) |
| **extra info:** | Burkholderia pseudomallei A101-10: microbial preparation by ZBS 2\| preparation for MALDI-ToF MS: M. Stämmler |
| **calibration standard:** | linear calibration using Escherichia coli DSM 3871 |
| **measurement method:** | D:\Methods\flexControlMethods\MaierMethods\ToM_200ns_20130611.par |
| **customer:** | RKI ZBS 2\| ZBS 6 |
| **measurement date/time:** | 2013-06-13T13:52:45.062+02:00 |
| **path to MS file:** | C:\Users\LaschP\Documents\MATLAB\Microbe MS testdata\ring trial RKI spectra\Sample_01\0_H3\1\1SLin |

##### Detailed quality test report

| Parameter | Noise | Baseline error | Number of peaks | Mean resolving power | General Quality |
|---|---|---|---|---|---|
| **absolut qt results** | 0.55098 | 2.0564 | 28 | 765.854 | |
| **rank** (out of 24 spectra) | 2 | 1 | 12 | 6 | 6 |
| **quality test result** | 78.6 | 78.6 | 64.3 | 71.4 | 71.8 |

---

#### # 2 - QUALITY TEST REPORT FOR MALDI-TOF MS SPECTRUM *'2013_06_14_Maren_Stämmler_0001'*

##### Metadata of actual MALDI-ToF test spectrum

| | |
|---|---|
| **genus / species / strain:** | RKI MALDI sample 01, QUANDHIP EQAE ring trial, strains provided by RKI \| ZBS 2 |
| **file id:** | 2013_06_14_Maren_Stämmler_0001 |
| **type:** | Measurement 02 |
| **Bruker ID:** | C5344CB2-A106-46AA-A51A-64817DBD40F2 |
| **NCBI ID (primary):** | 28450 |
| **NCBI ID (secondary):** | 28450 |
| **growth time:** | Optimal growth time between 24 - 72h |
| **growth temperature:** | 37°C |
| **growth conditions:** | Optimal aerobic or microaerophilic conditions |
| **growth medium:** | Columbia blood agar (Oxoid), 2nd passage on TSA or Caso agar, harvested by the 2nd passage |
| **sample treatment:** | Sample mixed with 20 mkL 100Â§ TFA, final TFA conc. approx. 80 perc.; approx. 30 Min treatment time; diluted 1:10 (vol); mixed with 1:1 HCCA\|TA2(A) |
| **spores:** | No |
| **concentration:** | Pellet produced by centrifugation (1 x 5 Min, 15,000 rpm) of a 500 mkL cell suspension, gamma ray irradiated (30 kGy) |
| **extra info:** | Burkholderia pseudomallei A101-10: microbial preparation by ZBS 2\| preparation for MALDI-ToF MS: M. Stämmler |
| **calibration standard:** | linear calibration using Escherichia coli DSM 3871 |
| **measurement method:** | D:\Methods\flexControlMethods\MaierMethods\ToM_200ns_20130611.par |
| **customer:** | RKI ZBS 2\| ZBS 6 |
| **measurement date/time:** | 2013-06-14T08:27:14.015+02:00 |
| **path to MS file:** | C:\Users\LaschP\Documents\MATLAB\Microbe MS testdata\ring trial RKI spectra\Sample_01\0_H4\1\1SLin |

#### Detailed quality test report

| Parameter | Noise | Baseline error | Number of peaks | Mean resolving power | General Quality |
|---|---|---|---|---|---|
| **absolut qt results** | 0.60156 | 2.5164 | 23 | 794.0641 | |
| **rank** (out of 24 spectra) | 14 | 4 | 18 | 3 | 13 |
| **quality test result** | 78.6 | 78.6 | 50 | 71.4 | 66.1 |

---

### # 3 - QUALITY TEST REPORT FOR MALDI-TOF MS SPECTRUM *'2013_06_13_Maren_Stämmler_0002'*

#### Metadata of actual MALDI-ToF test spectrum

| | |
|---|---|
| **genus / species / strain:** | RKI MALDI sample 02, QUANDHIP EQAE ring trial, strains provided by RKI \| ZBS 2 |
| **file id:** | 2013_06_13_Maren_Stämmler_0002 |
| **type:** | Measurement 01 |
| **Bruker ID:** | B1409699-0EFE-4C28-AAD4-FB638EF4774B |
| **NCBI ID (primary):** | 119857 |
| **NCBI ID (secondary):** | 119857 |
| **growth time:** | Optimal growth time between 24 - 72h |
| **growth temperature:** | 37°C |
| **growth conditions:** | Optimal aerobic or microaerophilic conditions |
| **growth medium:** | Heart cysteine agar (HCA) |
| **sample treatment:** | Sample mixed with 20 mkL 100Â§ TFA, final TFA conc. approx. 80 perc.; approx. 30 Min treatment time; diluted 1:10 (vol); mixed with 1:1 HCCA\|TA2(A) |
| **spores:** | No |
| **concentration:** | Pellet produced by centrifugation (1 x 5 Min, 15,000 rpm) of a 500 mkL cell suspension, gamma ray irradiated (30 kGy) |
| **extra info:** | Francisella tularensis ssp. holarctica Ft 32: microbial preparation by ZBS 2\| preparation for MALDI-ToF MS: M. Stämmler |
| **calibration standard:** | linear calibration using Escherichia coli DSM 3871 |
| **measurement method:** | D:\Methods\flexControlMethods\MaierMethods\ToM_200ns_20130611.par |
| **customer:** | RKI ZBS 2\| ZBS 6 |
| **measurement date/time:** | 2013-06-13T14:04:41.578+02:00 |
| **path to MS file:** | C:\Users\LaschP\Documents\MATLAB\Microbe MS testdata\ring trial RKI spectra\Sample_02\0_H5\1\1SLin |

#### Detailed quality test report

| Parameter | Noise | Baseline error | Number of peaks | Mean resolving power | General Quality |
|---|---|---|---|---|---|
| **absolut qt results** | 0.5603 | 4.3239 | 30 | 697.5869 | |
| **rank** (out of 24 spectra) | 5 | 13 | 10 | 12 | 10 |
| **quality test result** | 78.6 | 64.3 | 64.3 | 64.3 | 68.6 |

---

### # 4 - QUALITY TEST REPORT FOR MALDI-TOF MS SPECTRUM *'2013_06_14_Maren_Stämmler_0002'*

#### Metadata of actual MALDI-ToF test spectrum

| | |
|---|---|
| **genus / species / strain:** | RKI MALDI sample 02, QUANDHIP EQAE ring trial, strains provided by RKI \| ZBS 2 |
| **file id:** | 2013_06_14_Maren_Stämmler_0002 |
| **type:** | Measurement 02 |
| **Bruker ID:** | 4F9079BF-2237-4052-A736-259FD2ABA28A |
| **NCBI ID (primary):** | 119857 |
| **NCBI ID (secondary):** | 119857 |
| **growth time:** | Optimal growth time between 24 - 72h |
| **growth temperature:** | 37°C |
| **growth conditions:** | Optimal aerobic or microaerophilic conditions |
| **growth medium:** | Heart cysteine agar (HCA) |
| **sample treatment:** | Sample mixed with 20 mkL 100Â§ TFA, final TFA conc. approx. 80 perc.; approx. 30 Min treatment time; diluted 1:10 (vol); mixed with 1:1 HCCA\|TA2(A) |
| **spores:** | No |
| **concentration:** | Pellet produced by centrifugation (1 x 5 Min, 15,000 rpm) of a 500 mkL cell suspension, gamma ray irradiated (30 kGy) |
| **extra info:** | Francisella tularensis ssp. holarctica Ft 32: microbial preparation by ZBS 2\| preparation for MALDI-ToF MS: M. Stämmler |
| **calibration standard:** | linear calibration using Escherichia coli DSM 3871 |
| **measurement method:** | D:\Methods\flexControlMethods\MaierMethods\ToM_200ns_20130611.par |
| **customer:** | RKI ZBS 2\| ZBS 6 |
| **measurement date/time:** | 2013-06-14T08:33:44.531+02:00 |
| **path to MS file:** | C:\Users\LaschP\Documents\MATLAB\Microbe MS testdata\ring trial RKI spectra\Sample_02\0_H6\1\1SLin |

#### Detailed quality test report

| Parameter | Noise | Baseline error | Number of peaks | Mean resolving power | General Quality |
|---|---|---|---|---|---|
| **absolut qt results** | 0.6944 | 4.1334 | 27 | 662.5247 | |
| **rank** (out of 24 spectra) | 21 | 11 | 14 | 14 | 16 |
| **quality test result** | 71.4 | 64.3 | 57.1 | 64.3 | 63.6 |

---

### # 5 - QUALITY TEST REPORT FOR MALDI-TOF MS SPECTRUM *'2013_06_13_Maren_Stämmler_0003'*

#### Metadata of actual MALDI-ToF test spectrum

| | |
|---|---|
| **genus / species / strain:** | RKI MALDI sample 03, QUANDHIP EQAE ring trial, strains provided by RKI \| ZBS 2 |
| **file id:** | 2013_06_13_Maren_Stämmler_0003 |
| **type:** | Measurement 01 |
| **Bruker ID:** | 76182EF3-AD43-4157-A768-0D20D4A2FDF6 |
| **NCBI ID (primary):** | 36855 |
| **NCBI ID (secondary):** | 36855 |
| **growth time:** | Optimal growth time between 24 - 72h |
| **growth temperature:** | 37°C |
| **growth conditions:** | Optimal aerobic or microaerophilic conditions |
| **growth medium:** | Columbia blood agar (Oxoid), 2nd passage on TSA or Caso agar, harvested by the 2nd passage |
| **sample treatment:** | Sample mixed with 20 mkL 100Â§ TFA, final TFA conc. approx. 80 perc.; approx. 30 Min treatment time; diluted 1:10 (vol); mixed with 1:1 HCCA\|TA2(A) |
| **spores:** | No |
| **concentration:** | Pellet produced by centrifugation (1 x 5 Min, 15,000 rpm) of a 500 mkL cell suspension, gamma ray irradiated (30 kGy) |
| **extra info:** | Brucella canis A138-11: microbial preparation by ZBS 2\| preparation for MALDI-ToF MS: M. Stämmler |
| **calibration standard:** | linear calibration using Escherichia coli DSM 3871 |
| **measurement method:** | D:\Methods\flexControlMethods\MaierMethods\ToM_200ns_20130611.par |
| **customer:** | RKI ZBS 2\| ZBS 6 |
| **measurement date/time:** | 2013-06-13T14:11:32.906+02:00 |
| **path to MS file:** | C:\Users\LaschP\Documents\MATLAB\Microbe MS testdata\ring trial RKI spectra\Sample_03\0_H7\1\1SLin |

#### Detailed quality test report

| **Parameter** | **Noise** | **Baseline error** | **Number of peaks** | **Mean resolving power** | **General Quality** |
|---|---|---|---|---|---|
| **absolut qt results** | 0.52864 | 2.0835 | 25 | 795.0915 | |
| **rank** (out of 24 spectra) | 1 | 2 | 17 | 2 | 7 |
| **quality test result** | 85.7 | 78.6 | 57.1 | 71.4 | 71.1 |

---

### # 6 - QUALITY TEST REPORT FOR MALDI-TOF MS SPECTRUM *'2013_06_14_Maren_Stämmler_0003'*

#### Metadata of actual MALDI-ToF test spectrum

| | |
|---|---|
| **genus / species / strain:** | RKI MALDI sample 03, QUANDHIP EQAE ring trial, strains provided by RKI \| ZBS 2 |
| **file id:** | 2013_06_14_Maren_Stämmler_0003 |
| **type:** | Measurement 02 |
| **Bruker ID:** | 22710A07-8E7C-49C6-8EB2-00460D2E50DB |
| **NCBI ID (primary):** | 36855 |
| **NCBI ID (secondary):** | 36855 |
| **growth time:** | Optimal growth time between 24 - 72h |
| **growth temperature:** | 37°C |
| **growth conditions:** | Optimal aerobic or microaerophilic conditions |
| **growth medium:** | Columbia blood agar (Oxoid), 2nd passage on TSA or Caso agar, harvested by the 2nd passage |
| **sample treatment:** | Sample mixed with 20 mkL 100Â§ TFA, final TFA conc. approx. 80 perc.; approx. 30 Min treatment time; diluted 1:10 (vol); mixed with 1:1 HCCA\|TA2(A) |
| **spores:** | No |
| **concentration:** | Pellet produced by centrifugation (1 x 5 Min, 15,000 rpm) of a 500 mkL cell suspension, gamma ray irradiated (30 kGy) |
| **extra info:** | Brucella canis A138-11: microbial preparation by ZBS 2\| preparation for MALDI-ToF MS: M. Stämmler |
| **calibration standard:** | linear calibration using Escherichia coli DSM 3871 |
| **measurement method:** | D:\Methods\flexControlMethods\MaierMethods\ToM_200ns_20130611.par |
| **customer:** | RKI ZBS 2\| ZBS 6 |
| **measurement date/time:** | 2013-06-14T08:42:40.468+02:00 |
| **path to MS file:** | C:\Users\LaschP\Documents\MATLAB\Microbe MS testdata\ring trial RKI spectra\Sample_03\0_H8\1\1SLin |

#### Detailed quality test report

| **Parameter** | **Noise** | **Baseline error** | **Number of peaks** | **Mean resolving power** | **General Quality** |
|---|---|---|---|---|---|
| **absolut qt results** | 0.56969 | 3.4129 | 22 | 809.2989 | |
| **rank** (out of 24 spectra) | 8 | 8 | 19 | 1 | 14 |
| **quality test result** | 78.6 | 71.4 | 50 | 78.6 | 66.1 |

---

### # 7 - QUALITY TEST REPORT FOR MALDI-TOF MS SPECTRUM *'2013_06_17_Maren_Stämmler_0001'*

#### Metadata of actual MALDI-ToF test spectrum

| | |
|---|---|
| **genus / species / strain:** | RKI MALDI sample 04, QUANDHIP EQAE ring trial, strains provided by RKI \| ZBS 2 |
| **file id:** | 2013_06_17_Maren_Stämmler_0001 |
| **type:** | Measurement 01 |
| **Bruker ID:** | 7DEDDBEA-CB5D-49B9-B905-F8942788B5EC |
| **NCBI ID (primary):** | 1392 |
| **NCBI ID (secondary):** | 1392 |
| **growth time:** | Optimal growth time between 24 - 72h |

| | |
|---|---|
| **growth temperature:** | 37°C |
| **growth conditions:** | Optimal aerobic or microaerophilic conditions |
| **growth medium:** | Columbia blood agar (Oxoid), 2nd passage on TSA or Caso agar, harvested by the 2nd passage |
| **sample treatment:** | Sample mixed with 20 mkL 100Â§ TFA, final TFA conc. approx. 80 perc.; approx. 30 Min treatment time; diluted 1:10 (vol); mixed with 1:1 HCCA\|TA2(A) |
| **spores:** | No |
| **concentration:** | Pellet produced by centrifugation (1 x 5 Min, 15,000 rpm) of a 500 mkL cell suspension, gamma ray irradiated (30 kGy) |
| **extra info:** | Bacillus anthracis AMES: microbial preparation by ZBS 2\| preparation for MALDI-ToF MS: M. Stämmler |
| **calibration standard:** | linear calibration using Escherichia coli DSM 3871 |
| **measurement method:** | D:\Methods\flexControlMethods\MaierMethods\ToM_200ns_20130611.par |
| **customer:** | RKI ZBS 2\| ZBS 6 |
| **measurement date/time:** | 2013-06-17T09:21:01.468+02:00 |
| **path to MS file:** | C:\Users\LaschP\Documents\MATLAB\Microbe MS testdata\ring trial RKI spectra\Sample_04\0_E19\1\1SLin |

#### Detailed quality test report

| **Parameter** | **Noise** | **Baseline error** | **Number of peaks** | **Mean resolving power** | **General Quality** |
|---|---|---|---|---|---|
| **absolut qt results** | 0.59026 | 6.8788 | 28 | 586.4747 | |
| **rank** (out of 24 spectra) | 11 | 18 | 11 | 17 | 15 |
| **quality test result** | 78.6 | 50 | 64.3 | 57.1 | 65.4 |

---

### # 8 - QUALITY TEST REPORT FOR MALDI-TOF MS SPECTRUM *'2013_06_14_Maren_Stämmler_0004'*

#### Metadata of actual MALDI-ToF test spectrum

| | |
|---|---|
| **genus / species / strain:** | RKI MALDI sample 04, QUANDHIP EQAE ring trial, strains provided by RKI \| ZBS 2 |
| **file id:** | 2013_06_14_Maren_Stämmler_0004 |
| **type:** | Measurement 02 |
| **Bruker ID:** | 0AFF63F5-F910-4924-A353-4FAA9835969D |
| **NCBI ID (primary):** | 1392 |
| **NCBI ID (secondary):** | 1392 |
| **growth time:** | Optimal growth time between 24 - 72h |
| **growth temperature:** | 37°C |
| **growth conditions:** | Optimal aerobic or microaerophilic conditions |
| **growth medium:** | Columbia blood agar (Oxoid), 2nd passage on TSA or Caso agar, harvested by the 2nd passage |
| **sample treatment:** | Sample mixed with 25 mkL 100Â§ TFA, final TFA conc. approx. 80 perc.; 10 mkL 80 perc. TFA added; approx. 30 Min treatment time; diluted 1:10 (vol); mixed with 1:1 HCCA\|TA2(A) |
| **spores:** | No |
| **concentration:** | Pellet produced by centrifugation (1 x 5 Min, 15,000 rpm) of a 500 mkL cell suspension, gamma ray irradiated (30 kGy) |
| **extra info:** | Bacillus anthracis AMES: microbial preparation by ZBS 2\| preparation for MALDI-ToF MS: M. Stämmler |
| **calibration standard:** | linear calibration using Escherichia coli DSM 3871 |
| **measurement method:** | D:\Methods\flexControlMethods\MaierMethods\ToM_200ns_20130611.par |
| **customer:** | RKI ZBS 2\| ZBS 6 |
| **measurement date/time:** | 2013-06-14T08:52:26.953+02:00 |
| **path to MS file:** | C:\Users\LaschP\Documents\MATLAB\Microbe MS testdata\ring trial RKI spectra\Sample_04\0_H10\1\1SLin |

#### Detailed quality test report

| **Parameter** | **Noise** | **Baseline error** | **Number of peaks** | **Mean resolving power** | **General Quality** |
|---|---|---|---|---|---|
| **absolut qt results** | 0.89672 | 19.1088 | 20 | 548.6465 | |
| **rank** (out of 24 spectra) | 24 | 23 | 21 | 18 | 23 |
| **quality test result** | 57.1 | 28.6 | 42.9 | 50 | 46.1 |

---

### # 9 - QUALITY TEST REPORT FOR MALDI-TOF MS SPECTRUM *'2013_06_13_Maren_Stämmler_0004'*

#### Metadata of actual MALDI-ToF test spectrum

| | |
|---|---|
| **genus / species / strain:** | RKI MALDI sample 04, QUANDHIP EQAE ring trial, strains provided by RKI \| ZBS 2 |
| **file id:** | 2013_06_13_Maren_Stämmler_0004 |
| **type:** | Measurement 03 |
| **Bruker ID:** | 27965397-8164-4463-BB40-0FD314995834 |
| **NCBI ID (primary):** | 1392 |
| **NCBI ID (secondary):** | 1392 |
| **growth time:** | Optimal growth time between 24 - 72h |
| **growth temperature:** | 37°C |
| **growth conditions:** | Optimal aerobic or microaerophilic conditions |
| **growth medium:** | Columbia blood agar (Oxoid), 2nd passage on TSA or Caso agar, harvested by the 2nd passage |
| **sample treatment:** | Sample mixed with 25 mkL 100Â§ TFA, final TFA conc. approx. 80 perc.; 10 mkL 80 perc. TFA added; approx. 30 Min treatment time; diluted 1:10 (vol); mixed with 1:1 HCCA\|TA2(A) |
| **spores:** | No |
| **concentration:** | Pellet produced by centrifugation (1 x 5 Min, 15,000 rpm) of a 500 mkL cell suspension, gamma ray irradiated (30 kGy) |
| **extra info:** | Bacillus anthracis AMES: microbial preparation by ZBS 2\| preparation for MALDI-ToF MS: M. Stämmler |
| **calibration standard:** | linear calibration using Escherichia coli DSM 3871 |
| **measurement method:** | D:\Methods\flexControlMethods\MaierMethods\ToM_200ns_20130611.par |
| **customer:** | RKI ZBS 2\| ZBS 6 |
| **measurement date/time:** | 2013-06-13T14:19:29.671+02:00 |

| | |
|---|---|
| **path to MS file:** | C:\Users\LaschP\Documents\MATLAB\Microbe MS testdata\ring trial RKI spectra\Sample_04\0_H9\1\1SLin |

**Detailed quality test report**

| Parameter | Noise | Baseline error | Number of peaks | Mean resolving power | General Quality |
|---|---|---|---|---|---|
| **absolut qt results** | 0.56098 | 8.0194 | 32 | 600.1801 | |
| **rank** (out of 24 spectra) | 6 | 20 | 7 | 16 | 11 |
| **quality test result** | 78.6 | 50 | 71.4 | 57.1 | 68.2 |

---

#### # 10 - QUALITY TEST REPORT FOR MALDI-TOF MS SPECTRUM *'2013_06_13_Maren_Stämmler_0005'*

**Metadata of actual MALDI-ToF test spectrum**

| | |
|---|---|
| **genus / species / strain:** | RKI MALDI sample 05, QUANDHIP EQAE ring trial, strains provided by RKI \| ZBS 2 |
| **file id:** | 2013_06_13_Maren_Stämmler_0005 |
| **type:** | Measurement 01 |
| **Bruker ID:** | DB87AC50-0ED5-4ACA-B2F9-4B34DE757E1B |
| **NCBI ID (primary):** | 529 |
| **NCBI ID (secondary):** | 529 |
| **growth time:** | Optimal growth time between 24 - 72h |
| **growth temperature:** | 37°C |
| **growth conditions:** | Optimal aerobic or microaerophilic conditions |
| **growth medium:** | Columbia blood agar (Oxoid), 2nd passage on TSA or Caso agar, harvested by the 2nd passage |
| **sample treatment:** | Sample mixed with 20 mkL 100Â§ TFA, final TFA conc. approx. 80 perc.; approx. 30 Min treatment time; diluted 1:10 (vol); mixed with 1:1 HCCA\|TA2(A) |
| **spores:** | No |
| **concentration:** | Pellet produced by centrifugation (1 x 5 Min, 15,000 rpm) of a 500 mkL cell suspension, gamma ray irradiated (30 kGy) |
| **extra info:** | Ochrobactrum anthropi A-148-11: microbial preparation by ZBS 2\| preparation for MALDI-ToF MS: M. Stämmler |
| **calibration standard:** | linear calibration using Escherichia coli DSM 3871 |
| **measurement method:** | D:\Methods\flexControlMethods\MaierMethods\ToM_200ns_20130611.par |
| **customer:** | RKI ZBS 2\| ZBS 6 |
| **measurement date/time:** | 2013-06-13T14:25:30.359+02:00 |
| **path to MS file:** | C:\Users\LaschP\Documents\MATLAB\Microbe MS testdata\ring trial RKI spectra\Sample_05\0_H11\1\1SLin |

**Detailed quality test report**

| Parameter | Noise | Baseline error | Number of peaks | Mean resolving power | General Quality |
|---|---|---|---|---|---|
| **absolut qt results** | 0.6001 | 3.7571 | 30 | 756.5342 | |
| **rank** (out of 24 spectra) | 13 | 9 | 9 | 8 | 8 |
| **quality test result** | 78.6 | 71.4 | 64.3 | 71.4 | 70.7 |

---

#### # 11 - QUALITY TEST REPORT FOR MALDI-TOF MS SPECTRUM *'2013_06_14_Maren_Stämmler_0005'*

**Metadata of actual MALDI-ToF test spectrum**

| | |
|---|---|
| **genus / species / strain:** | RKI MALDI sample 05, QUANDHIP EQAE ring trial, strains provided by RKI \| ZBS 2 |
| **file id:** | 2013_06_14_Maren_Stämmler_0005 |
| **type:** | Measurement 02 |
| **Bruker ID:** | B8441899-48FF-4802-B5F8-4491FED91EF2 |
| **NCBI ID (primary):** | 529 |
| **NCBI ID (secondary):** | 529 |
| **growth time:** | Optimal growth time between 24 - 72h |
| **growth temperature:** | 37°C |
| **growth conditions:** | Optimal aerobic or microaerophilic conditions |
| **growth medium:** | Columbia blood agar (Oxoid), 2nd passage on TSA or Caso agar, harvested by the 2nd passage |
| **sample treatment:** | Sample mixed with 20 mkL 100Â§ TFA, final TFA conc. approx. 80 perc.; approx. 30 Min treatment time; diluted 1:10 (vol); mixed with 1:1 HCCA\|TA2(A) |
| **spores:** | No |
| **concentration:** | Pellet produced by centrifugation (1 x 5 Min, 15,000 rpm) of a 500 mkL cell suspension, gamma ray irradiated (30 kGy) |
| **extra info:** | Ochrobactrum anthropi A-148-11: microbial preparation by ZBS 2\| preparation for MALDI-ToF MS: M. Stämmler |
| **calibration standard:** | linear calibration using Escherichia coli DSM 3871 |
| **measurement method:** | D:\Methods\flexControlMethods\MaierMethods\ToM_200ns_20130611.par |
| **customer:** | RKI ZBS 2\| ZBS 6 |
| **measurement date/time:** | 2013-06-14T09:00:58.531+02:00 |
| **path to MS file:** | C:\Users\LaschP\Documents\MATLAB\Microbe MS testdata\ring trial RKI spectra\Sample_05\0_H12\1\1SLin |

**Detailed quality test report**

| Parameter | Noise | Baseline error | Number of peaks | Mean resolving power | General Quality |
|---|---|---|---|---|---|
| **absolut qt results** | 0.63783 | 6.784 | 37 | 712.0333 | |
| **rank** (out of 24 spectra) | 18 | 16 | 2 | 10 | 9 |
| **quality test result** | 71.4 | 50 | 78.6 | 64.3 | 70 |

### # 12 - QUALITY TEST REPORT FOR MALDI-TOF MS SPECTRUM *'2013_06_13_Maren_Stämmler_0006'*

#### Metadata of actual MALDI-ToF test spectrum

| | |
|---|---|
| **genus / species / strain:** | RKI MALDI sample 06, QUANDHIP EQAE ring trial, strains provided by RKI \| ZBS 2 |
| **file id:** | 2013_06_13_Maren_Stämmler_0006 |
| **type:** | Measurement 01 |
| **Bruker ID:** | A91F9B76-08DA-40E6-9154-5F7CE8EF3789 |
| **NCBI ID (primary):** | 633 |
| **NCBI ID (secondary):** | 633 |
| **growth time:** | Optimal growth time between 24 - 72h |
| **growth temperature:** | 37°C |
| **growth conditions:** | Optimal aerobic or microaerophilic conditions |
| **growth medium:** | Columbia blood agar (Oxoid), 2nd passage on TSA or Caso agar, harvested by the 2nd passage |
| **sample treatment:** | Sample mixed with 20 mkL 100Â§ TFA, final TFA conc. approx. 80 perc.; approx. 30 Min treatment time; diluted 1:10 (vol); mixed with 1:1 HCCA\|TA2(A) |
| **spores:** | No |
| **concentration:** | Pellet produced by centrifugation (1 x 5 Min, 15,000 rpm) of a 500 mkL cell suspension, gamma ray irradiated (30 kGy) |
| **extra info:** | Yersinia pseudotuberculosis III: microbial preparation by ZBS 2\| preparation for MALDI-ToF MS: M. Stämmler |
| **calibration standard:** | linear calibration using Escherichia coli DSM 3871 |
| **measurement method:** | D:\Methods\flexControlMethods\MaierMethods\ToM_200ns_20130611.par |
| **customer:** | RKI ZBS 2\| ZBS 6 |
| **measurement date/time:** | 2013-06-13T14:36:26.328+02:00 |
| **path to MS file:** | C:\Users\LaschP\Documents\MATLAB\Microbe MS testdata\ring trial RKI spectra\Sample_06\0_H13\1\1SLin |

#### Detailed quality test report

| **Parameter** | **Noise** | **Baseline error** | **Number of peaks** | **Mean resolving power** | **General Quality** |
|---|---|---|---|---|---|
| **absolut qt results** | 0.55874 | 2.8589 | 36 | 720.8583 | |
| **rank** (out of 24 spectra) | 4 | 5 | 3 | 9 | 1 |
| **quality test result** | 78.6 | 71.4 | 78.6 | 64.3 | 75.4 |

### # 13 - QUALITY TEST REPORT FOR MALDI-TOF MS SPECTRUM *'2013_06_14_Maren_Stämmler_0006'*

#### Metadata of actual MALDI-ToF test spectrum

| | |
|---|---|
| **genus / species / strain:** | RKI MALDI sample 06, QUANDHIP EQAE ring trial, strains provided by RKI \| ZBS 2 |
| **file id:** | 2013_06_14_Maren_Stämmler_0006 |
| **type:** | Measurement 02 |
| **Bruker ID:** | 0D79F381-905B-4F07-8C28-4E0E02288EDC |
| **NCBI ID (primary):** | 633 |
| **NCBI ID (secondary):** | 633 |
| **growth time:** | Optimal growth time between 24 - 72h |
| **growth temperature:** | 37°C |
| **growth conditions:** | Optimal aerobic or microaerophilic conditions |
| **growth medium:** | Columbia blood agar (Oxoid), 2nd passage on TSA or Caso agar, harvested by the 2nd passage |
| **sample treatment:** | Sample mixed with 20 mkL 100Â§ TFA, final TFA conc. approx. 80 perc.; approx. 30 Min treatment time; diluted 1:10 (vol); mixed with 1:1 HCCA\|TA2(A) |
| **spores:** | No |
| **concentration:** | Pellet produced by centrifugation (1 x 5 Min, 15,000 rpm) of a 500 mkL cell suspension, gamma ray irradiated (30 kGy) |
| **extra info:** | Yersinia pseudotuberculosis III: microbial preparation by ZBS 2\| preparation for MALDI-ToF MS: M. Stämmler |
| **calibration standard:** | linear calibration using Escherichia coli DSM 3871 |
| **measurement method:** | D:\Methods\flexControlMethods\MaierMethods\ToM_200ns_20130611.par |
| **customer:** | RKI ZBS 2\| ZBS 6 |
| **measurement date/time:** | 2013-06-14T09:22:37.468+02:00 |
| **path to MS file:** | C:\Users\LaschP\Documents\MATLAB\Microbe MS testdata\ring trial RKI spectra\Sample_06\0_H14\1\1SLin |

#### Detailed quality test report

| **Parameter** | **Noise** | **Baseline error** | **Number of peaks** | **Mean resolving power** | **General Quality** |
|---|---|---|---|---|---|
| **absolut qt results** | 0.6042 | 3.3495 | 34 | 708.8133 | |
| **rank** (out of 24 spectra) | 15 | 7 | 5 | 11 | 4 |
| **quality test result** | 78.6 | 71.4 | 71.4 | 64.3 | 72.5 |

### # 14 - QUALITY TEST REPORT FOR MALDI-TOF MS SPECTRUM *'2013_06_13_Maren_Stämmler_0007'*

#### Metadata of actual MALDI-ToF test spectrum

| | |
|---|---|
| **genus / species / strain:** | RKI MALDI sample 07, QUANDHIP EQAE ring trial, strains provided by RKI \| ZBS 2 |
| **file id:** | 2013_06_13_Maren_Stämmler_0007 |
| **type:** | Measurement 01 |
| **Bruker ID:** | 9E2A8556-22E3-4402-90AF-098AB16AC043 |

| | |
|---|---|
| **NCBI ID (primary):** | 13373 |
| **NCBI ID (secondary):** | 13373 |
| **growth time:** | Optimal growth time between 24 - 72h |
| **growth temperature:** | 37°C |
| **growth conditions:** | Optimal aerobic or microaerophilic conditions |
| **growth medium:** | Columbia blood agar (Oxoid), 2nd passage on TSA or Caso agar, harvested by the 2nd passage |
| **sample treatment:** | Sample mixed with 20 mkL 100Â§ TFA, final TFA conc. approx. 80 perc.; approx. 30 Min treatment time; diluted 1:10 (vol); mixed with 1:1 HCCA\|TA2(A) |
| **spores:** | No |
| **concentration:** | Pellet produced by centrifugation (1 x 5 Min, 15,000 rpm) of a 500 mkL cell suspension, gamma ray irradiated (30 kGy) |
| **extra info:** | Burkholderia mallei A106-3: microbial preparation by ZBS 2\| preparation for MALDI-ToF MS: M. Stämmler |
| **calibration standard:** | linear calibration using Escherichia coli DSM 3871 |
| **measurement method:** | D:\Methods\flexControlMethods\MaierMethods\ToM_200ns_20130611.par |
| **customer:** | RKI ZBS 2\| ZBS 6 |
| **measurement date/time:** | 2013-06-13T14:39:43.625+02:00 |
| **path to MS file:** | C:\Users\LaschP\Documents\MATLAB\Microbe MS testdata\ring trial RKI spectra\Sample_07\0_H15\1\1SLin |

##### Detailed quality test report

| **Parameter** | **Noise** | **Baseline error** | **Number of peaks** | **Mean resolving power** | **General Quality** |
|---|---|---|---|---|---|
| **absolut qt results** | 0.5829 | 4.8456 | 31 | 765.58 | |
| **rank** (out of 24 spectra) | 10 | 14 | 8 | 7 | 5 |
| **quality test result** | 78.6 | 64.3 | 71.4 | 71.4 | 72.5 |

---

#### # 15 - QUALITY TEST REPORT FOR MALDI-TOF MS SPECTRUM *'2013_06_14_Maren_Stämmler_0007'*

##### Metadata of actual MALDI-ToF test spectrum

| | |
|---|---|
| **genus / species / strain:** | RKI MALDI sample 07, QUANDHIP EQAE ring trial, strains provided by RKI \| ZBS 2 |
| **file id:** | 2013_06_14_Maren_Stämmler_0007 |
| **type:** | Measurement 02 |
| **Bruker ID:** | 92FCB042-DB6D-4DCC-9783-56CB2B43BCD5 |
| **NCBI ID (primary):** | 13373 |
| **NCBI ID (secondary):** | 13373 |
| **growth time:** | Optimal growth time between 24 - 72h |
| **growth temperature:** | 37°C |
| **growth conditions:** | Optimal aerobic or microaerophilic conditions |
| **growth medium:** | Columbia blood agar (Oxoid), 2nd passage on TSA or Caso agar, harvested by the 2nd passage |
| **sample treatment:** | Sample mixed with 20 mkL 100Â§ TFA, final TFA conc. approx. 80 perc.; approx. 30 Min treatment time; diluted 1:10 (vol); mixed with 1:1 HCCA\|TA2(A) |
| **spores:** | No |
| **concentration:** | Pellet produced by centrifugation (1 x 5 Min, 15,000 rpm) of a 500 mkL cell suspension, gamma ray irradiated (30 kGy) |
| **extra info:** | Burkholderia mallei A106-3: microbial preparation by ZBS 2\| preparation for MALDI-ToF MS: M. Stämmler |
| **calibration standard:** | linear calibration using Escherichia coli DSM 3871 |
| **measurement method:** | D:\Methods\flexControlMethods\MaierMethods\ToM_200ns_20130611.par |
| **customer:** | RKI ZBS 2\| ZBS 6 |
| **measurement date/time:** | 2013-06-14T09:30:51.015+02:00 |
| **path to MS file:** | C:\Users\LaschP\Documents\MATLAB\Microbe MS testdata\ring trial RKI spectra\Sample_07\0_H16\1\1SLin |

##### Detailed quality test report

| **Parameter** | **Noise** | **Baseline error** | **Number of peaks** | **Mean resolving power** | **General Quality** |
|---|---|---|---|---|---|
| **absolut qt results** | 0.64934 | 4.2132 | 37 | 787.9516 | |
| **rank** (out of 24 spectra) | 19 | 12 | 1 | 4 | 3 |
| **quality test result** | 71.4 | 64.3 | 78.6 | 71.4 | 73.2 |

---

#### # 16 - QUALITY TEST REPORT FOR MALDI-TOF MS SPECTRUM *'2013_06_17_Maren_Stämmler_0002'*

##### Metadata of actual MALDI-ToF test spectrum

| | |
|---|---|
| **genus / species / strain:** | RKI MALDI sample 08, QUANDHIP EQAE ring trial, strains provided by RKI \| ZBS 2 |
| **file id:** | 2013_06_17_Maren_Stämmler_0002 |
| **type:** | Measurement 01 |
| **Bruker ID:** | 8111BE2B-254B-4723-A815-87E8BF9DAC5F |
| **NCBI ID (primary):** | 57975 |
| **NCBI ID (secondary):** | 57975 |
| **growth time:** | Optimal growth time between 24 - 72h |
| **growth temperature:** | 37°C |
| **growth conditions:** | Optimal aerobic or microaerophilic conditions |
| **growth medium:** | Columbia blood agar (Oxoid), 2nd passage on TSA or Caso agar, harvested by the 2nd passage |
| **sample treatment:** | Sample mixed with 30 mkL 100Â§ TFA, final TFA conc. approx. 80 perc.; 10 mkL 80 perc. TFA added; approx. 30 Min treatment time; diluted 1:10 (vol); mixed with 1:1 HCCA\|TA2(A) |
| **spores:** | No |
| **concentration:** | Pellet produced by centrifugation (1 x 5 Min, 15,000 rpm) of a 500 mkL cell suspension, gamma ray irradiated (30 kGy) |
| **extra info:** | Burkholderia thailandensis E125: microbial preparation by ZBS 2\| preparation for MALDI-ToF MS: M. Stämmler |
| **calibration standard:** | linear calibration using Escherichia coli DSM 3871 |

| | |
|---|---|
| **measurement method:** | D:\Methods\flexControlMethods\MaierMethods\ToM_200ns_20130611.par |
| **customer:** | RKI ZBS 2\| ZBS 6 |
| **measurement date/time:** | 2013-06-17T09:23:57.843+02:00 |
| **path to MS file:** | C:\Users\LaschP\Documents\MATLAB\Microbe MS testdata\ring trial RKI spectra\Sample_08\0_E20\1\1SLin |

**Detailed quality test report**

| **Parameter** | **Noise** | **Baseline error** | **Number of peaks** | **Mean resolving power** | **General Quality** |
|---|---|---|---|---|---|
| **absolut qt results** | 0.67939 | 7.7174 | 27 | 643.9615 | |
| **rank** (out of 24 spectra) | 20 | 19 | 13 | 15 | 18 |
| **quality test result** | 71.4 | 50 | 57.1 | 57.1 | 60.4 |

---

#### # 17 - QUALITY TEST REPORT FOR MALDI-TOF MS SPECTRUM *'2013_06_13_Maren_Stämmler_0008'*

**Metadata of actual MALDI-ToF test spectrum**

| | |
|---|---|
| **genus / species / strain:** | RKI MALDI sample 08, QUANDHIP EQAE ring trial, strains provided by RKI \| ZBS 2 |
| **file id:** | 2013_06_13_Maren_Stämmler_0008 |
| **type:** | Measurement 02 |
| **Bruker ID:** | 70156A73-5473-4C86-A9AA-061FA9ACF803 |
| **NCBI ID (primary):** | 57975 |
| **NCBI ID (secondary):** | 57975 |
| **growth time:** | Optimal growth time between 24 - 72h |
| **growth temperature:** | 37°C |
| **growth conditions:** | Optimal aerobic or microaerophilic conditions |
| **growth medium:** | Columbia blood agar (Oxoid), 2nd passage on TSA or Caso agar, harvested by the 2nd passage |
| **sample treatment:** | Sample mixed with 20 mkL 100Â§ TFA, final TFA conc. approx. 80 perc.; approx. 30 Min treatment time; diluted 1:10 (vol); mixed with 1:1 HCCA\|TA2(A) |
| **spores:** | No |
| **concentration:** | Pellet produced by centrifugation (1 x 5 Min, 15,000 rpm) of a 500 mkL cell suspension, gamma ray irradiated (30 kGy) |
| **extra info:** | Burkholderia thailandensis E125: microbial preparation by ZBS 2\| preparation for MALDI-ToF MS: M. Stämmler |
| **calibration standard:** | linear calibration using Escherichia coli DSM 3871 |
| **measurement method:** | D:\Methods\flexControlMethods\MaierMethods\ToM_200ns_20130611.par |
| **customer:** | RKI ZBS 2\| ZBS 6 |
| **measurement date/time:** | 2013-06-13T14:48:34.328+02:00 |
| **path to MS file:** | C:\Users\LaschP\Documents\MATLAB\Microbe MS testdata\ring trial RKI spectra\Sample_08\0_H17\1\1SLin |

**Detailed quality test report**

| **Parameter** | **Noise** | **Baseline error** | **Number of peaks** | **Mean resolving power** | **General Quality** |
|---|---|---|---|---|---|
| **absolut qt results** | 0.62943 | 3.2104 | 32 | 768.9188 | |
| **rank** (out of 24 spectra) | 16 | 6 | 6 | 5 | 2 |
| **quality test result** | 78.6 | 71.4 | 71.4 | 71.4 | 73.6 |

---

#### # 18 - QUALITY TEST REPORT FOR MALDI-TOF MS SPECTRUM *'2013_06_14_Maren_Stämmler_0008'*

**Metadata of actual MALDI-ToF test spectrum**

| | |
|---|---|
| **genus / species / strain:** | RKI MALDI sample 08, QUANDHIP EQAE ring trial, strains provided by RKI \| ZBS 2 |
| **file id:** | 2013_06_14_Maren_Stämmler_0008 |
| **type:** | Measurement 03 |
| **Bruker ID:** | BE203424-ED2C-4A25-BA68-10D562636EDE |
| **NCBI ID (primary):** | 57975 |
| **NCBI ID (secondary):** | 57975 |
| **growth time:** | Optimal growth time between 24 - 72h |
| **growth temperature:** | 37°C |
| **growth conditions:** | Optimal aerobic or microaerophilic conditions |
| **growth medium:** | Columbia blood agar (Oxoid), 2nd passage on TSA or Caso agar, harvested by the 2nd passage |
| **sample treatment:** | Sample mixed with 20 mkL 100Â§ TFA, final TFA conc. approx. 80 perc.; approx. 30 Min treatment time; diluted 1:10 (vol); mixed with 1:1 HCCA\|TA2(A) |
| **spores:** | No |
| **concentration:** | Pellet produced by centrifugation (1 x 5 Min, 15,000 rpm) of a 500 mkL cell suspension, gamma ray irradiated (30 kGy) |
| **extra info:** | Burkholderia thailandensis E125: microbial preparation by ZBS 2\| preparation for MALDI-ToF MS: M. Stämmler |
| **calibration standard:** | linear calibration using Escherichia coli DSM 3871 |
| **measurement method:** | D:\Methods\flexControlMethods\MaierMethods\ToM_200ns_20130611.par |
| **customer:** | RKI ZBS 2\| ZBS 6 |
| **measurement date/time:** | 2013-06-14T09:36:55.281+02:00 |
| **path to MS file:** | C:\Users\LaschP\Documents\MATLAB\Microbe MS testdata\ring trial RKI spectra\Sample_08\0_H18\1\1SLin |

**Detailed quality test report**

| **Parameter** | **Noise** | **Baseline error** | **Number of peaks** | **Mean resolving power** | **General Quality** |
|---|---|---|---|---|---|
| **absolut qt results** | 0.69997 | 6.7848 | 34 | 694.5404 | |

| | | | | | |
|---|---|---|---|---|---|
| **rank** (out of 24 spectra) | 22 | 17 | 4 | 13 | 12 |
| **quality test result** | 71.4 | 50 | 71.4 | 64.3 | 67.1 |

---

#### # 19 - QUALITY TEST REPORT FOR MALDI-TOF MS SPECTRUM *'2013_06_17_Maren_Stämmler_0003'*

##### Metadata of actual MALDI-ToF test spectrum

| | |
|---|---|
| **genus / species / strain:** | RKI MALDI sample 09, QUANDHIP EQAE ring trial, strains provided by RKI \| ZBS 2 |
| **file id:** | 2013_06_17_Maren_Stämmler_0003 |
| **type:** | Measurement 01 |
| **Bruker ID:** | 8EEA47D8-C42B-4229-9813-6DBBF397112D |
| **NCBI ID (primary):** | 632 |
| **NCBI ID (secondary):** | 632 |
| **growth time:** | Optimal growth time between 24 - 72h |
| **growth temperature:** | 37°C |
| **growth conditions:** | Optimal aerobic or microaerophilic conditions |
| **growth medium:** | Columbia blood agar (Oxoid), 2nd passage on TSA or Caso agar, harvested by the 2nd passage |
| **sample treatment:** | Sample mixed with 30 mkL 100Â§ TFA, final TFA conc. approx. 80 perc.; approx. 30 Min treatment time; diluted 1:10 (vol); mixed with 1:1 HCCA\|TA2(A) |
| **spores:** | No |
| **concentration:** | Pellet produced by centrifugation (1 x 5 Min, 15,000 rpm) of a 500 mkL cell suspension, gamma ray irradiated (30 kGy) |
| **extra info:** | Yersinia pestis A106-2: microbial preparation by ZBS 2\| preparation for MALDI-ToF MS: M. Stämmler |
| **calibration standard:** | linear calibration using Escherichia coli DSM 3871 |
| **measurement method:** | D:\Methods\flexControlMethods\MaierMethods\ToM_200ns_20130611.par |
| **customer:** | RKI ZBS 2\| ZBS 6 |
| **measurement date/time:** | 2013-06-17T09:29:00.578+02:00 |
| **path to MS file:** | C:\Users\LaschP\Documents\MATLAB\Microbe MS testdata\ring trial RKI spectra\Sample_09\0_E21\1\1SLin |

##### Detailed quality test report

| Parameter | Noise | Baseline error | Number of peaks | Mean resolving power | General Quality |
|---|---|---|---|---|---|
| **absolut qt results** | 0.56131 | 5.4022 | 12 | 474.4032 | |
| **rank** (out of 24 spectra) | 7 | 15 | 24 | 23 | 22 |
| **quality test result** | 78.6 | 57.1 | 21.4 | 42.9 | 47.1 |

---

#### # 20 - QUALITY TEST REPORT FOR MALDI-TOF MS SPECTRUM *'2013_06_13_Maren_Stämmler_0009'*

##### Metadata of actual MALDI-ToF test spectrum

| | |
|---|---|
| **genus / species / strain:** | RKI MALDI sample 09, QUANDHIP EQAE ring trial, strains provided by RKI \| ZBS 2 |
| **file id:** | 2013_06_13_Maren_Stämmler_0009 |
| **type:** | Measurement 02 |
| **Bruker ID:** | 70E47FB9-207D-4ABA-83A5-5463D6F150AA |
| **NCBI ID (primary):** | 632 |
| **NCBI ID (secondary):** | 632 |
| **growth time:** | Optimal growth time between 24 - 72h |
| **growth temperature:** | 37°C |
| **growth conditions:** | Optimal aerobic or microaerophilic conditions |
| **growth medium:** | Columbia blood agar (Oxoid), 2nd passage on TSA or Caso agar, harvested by the 2nd passage |
| **sample treatment:** | Sample mixed with 15 mkL 100Â§ TFA, final TFA conc. approx. 80 perc.; approx. 30 Min treatment time; diluted 1:10 (vol); mixed with 1:1 HCCA\|TA2(A) |
| **spores:** | No |
| **concentration:** | Pellet produced by centrifugation (1 x 5 Min, 15,000 rpm) of a 500 mkL cell suspension, gamma ray irradiated (30 kGy) |
| **extra info:** | Yersinia pestis A106-2: microbial preparation by ZBS 2\| preparation for MALDI-ToF MS: M. Stämmler |
| **calibration standard:** | linear calibration using Escherichia coli DSM 3871 |
| **measurement method:** | D:\Methods\flexControlMethods\MaierMethods\ToM_200ns_20130611.par |
| **customer:** | RKI ZBS 2\| ZBS 6 |
| **measurement date/time:** | 2013-06-13T14:57:12.250+02:00 |
| **path to MS file:** | C:\Users\LaschP\Documents\MATLAB\Microbe MS testdata\ring trial RKI spectra\Sample_09\0_H19\1\1SLin |

##### Detailed quality test report

| Parameter | Noise | Baseline error | Number of peaks | Mean resolving power | General Quality |
|---|---|---|---|---|---|
| **absolut qt results** | 0.57448 | 2.2808 | 15 | 510.5523 | |
| **rank** (out of 24 spectra) | 9 | 3 | 23 | 22 | 21 |
| **quality test result** | 78.6 | 78.6 | 28.6 | 42.9 | 53.2 |

---

#### # 21 - QUALITY TEST REPORT FOR MALDI-TOF MS SPECTRUM *'2013_06_14_Maren_Stämmler_0009'*

##### Metadata of actual MALDI-ToF test spectrum

| | |
|---|---|
| **genus / species / strain:** | RKI MALDI sample 09, QUANDHIP EQAE ring trial, strains provided by RKI \| ZBS 2 |

| | |
|---|---|
| **file id:** | 2013_06_14_Maren_Stämmler_0009 |
| **type:** | Measurement 03 |
| **Bruker ID:** | 144E04DA-E6C7-4A7F-BD46-568E8747ADE2 |
| **NCBI ID (primary):** | 632 |
| **NCBI ID (secondary):** | 632 |
| **growth time:** | Optimal growth time between 24 - 72h |
| **growth temperature:** | 37°C |
| **growth conditions:** | Optimal aerobic or microaerophilic conditions |
| **growth medium:** | Columbia blood agar (Oxoid), 2nd passage on TSA or Caso agar, harvested by the 2nd passage |
| **sample treatment:** | Sample mixed with 15 mkL 100Â§ TFA, final TFA conc. approx. 80 perc.; approx. 30 Min treatment time; diluted 1:10 (vol); mixed with 1:1 HCCA\|TA2(A) |
| **spores:** | No |
| **concentration:** | Pellet produced by centrifugation (1 x 5 Min, 15,000 rpm) of a 500 mkL cell suspension, gamma ray irradiated (30 kGy) |
| **extra info:** | Yersinia pestis A106-2: microbial preparation by ZBS 2\| preparation for MALDI-ToF MS: M. Stämmler |
| **calibration standard:** | linear calibration using Escherichia coli DSM 3871 |
| **measurement method:** | D:\Methods\flexControlMethods\MaierMethods\ToM_200ns_20130611.par |
| **customer:** | RKI ZBS 2\| ZBS 6 |
| **measurement date/time:** | 2013-06-14T09:44:17.781+02:00 |
| **path to MS file:** | C:\Users\LaschP\Documents\MATLAB\Microbe MS testdata\ring trial RKI spectra\Sample_09\0_H20\1\1SLin |

##### Detailed quality test report

| Parameter | Noise | Baseline error | Number of peaks | Mean resolving power | General Quality |
|---|---|---|---|---|---|
| **absolut qt results** | 0.55511 | 3.9614 | 21 | 522.1378 | |
| **rank** (out of 24 spectra) | 3 | 10 | 20 | 21 | 17 |
| **quality test result** | 78.6 | 64.3 | 50 | 50 | 60.7 |

---

#### # 22 - QUALITY TEST REPORT FOR MALDI-TOF MS SPECTRUM *'2013_06_17_Maren_Stämmler_0004'*

##### Metadata of actual MALDI-ToF test spectrum

| | |
|---|---|
| **genus / species / strain:** | RKI MALDI sample 10, QUANDHIP EQAE ring trial, strains provided by RKI \| ZBS 2 |
| **file id:** | 2013_06_17_Maren_Stämmler_0004 |
| **type:** | Measurement 01 |
| **Bruker ID:** | BF74B46D-3F52-4707-A3D6-5DC37A88756C |
| **NCBI ID (primary):** | 1428 |
| **NCBI ID (secondary):** | 1428 |
| **growth time:** | Optimal growth time between 24 - 72h |
| **growth temperature:** | 37°C |
| **growth conditions:** | Optimal aerobic or microaerophilic conditions |
| **growth medium:** | Columbia blood agar (Oxoid), 2nd passage on TSA or Caso agar, harvested by the 2nd passage |
| **sample treatment:** | Sample mixed with 80 mkL 100Â§ TFA, final TFA conc. approx. 80 perc.; approx. 30 Min treatment time; diluted 1:10 (vol); mixed with 1:1 HCCA\|TA2(A) |
| **spores:** | No |
| **concentration:** | Pellet produced by centrifugation (1 x 5 Min, 15,000 rpm) of a 500 mkL cell suspension, gamma ray irradiated (30 kGy) |
| **extra info:** | Bacillus thuringiensis DSM 350: microbial preparation by ZBS 2\| preparation for MALDI-ToF MS: M. Stämmler |
| **calibration standard:** | linear calibration using Escherichia coli DSM 3871 |
| **measurement method:** | D:\Methods\flexControlMethods\MaierMethods\ToM_200ns_20130611.par |
| **customer:** | RKI ZBS 2\| ZBS 6 |
| **measurement date/time:** | 2013-06-17T09:39:08.984+02:00 |
| **path to MS file:** | C:\Users\LaschP\Documents\MATLAB\Microbe MS testdata\ring trial RKI spectra\Sample_10\0_E22\1\1SLin |

##### Detailed quality test report

| Parameter | Noise | Baseline error | Number of peaks | Mean resolving power | General Quality |
|---|---|---|---|---|---|
| **absolut qt results** | 0.59604 | 18.3112 | 25 | 534.9501 | |
| **rank** (out of 24 spectra) | 12 | 22 | 16 | 19 | 19 |
| **quality test result** | 78.6 | 28.6 | 57.1 | 50 | 58.2 |

---

#### # 23 - QUALITY TEST REPORT FOR MALDI-TOF MS SPECTRUM *'2013_06_13_Maren_Stämmler_0010'*

##### Metadata of actual MALDI-ToF test spectrum

| | |
|---|---|
| **genus / species / strain:** | RKI MALDI sample 10, QUANDHIP EQAE ring trial, strains provided by RKI \| ZBS 2 |
| **file id:** | 2013_06_13_Maren_Stämmler_0010 |
| **type:** | Measurement 02 |
| **Bruker ID:** | 31336184-977D-4562-99B6-1F28627C6218 |
| **NCBI ID (primary):** | 1428 |
| **NCBI ID (secondary):** | 1428 |
| **growth time:** | Optimal growth time between 24 - 72h |
| **growth temperature:** | 37°C |
| **growth conditions:** | Optimal aerobic or microaerophilic conditions |
| **growth medium:** | Columbia blood agar (Oxoid), 2nd passage on TSA or Caso agar, harvested by the 2nd passage |
| **sample treatment:** | Sample mixed with 25 mkL 100Â§ TFA, final TFA conc. approx. 80 perc.; approx. 30 Min treatment time; diluted 1:10 (vol); mixed with 1:1 HCCA\|TA2(A) |
| **spores:** | No |

| | |
|---|---|
| **concentration:** | Pellet produced by centrifugation (1 x 5 Min, 15,000 rpm) of a 500 mkL cell suspension, gamma ray irradiated (30 kGy) |
| **extra info:** | Bacillus thuringiensis DSM 350: microbial preparation by ZBS 2\| preparation for MALDI-ToF MS: M. Stämmler |
| **calibration standard:** | linear calibration using Escherichia coli DSM 3871 |
| **measurement method:** | D:\Methods\flexControlMethods\MaierMethods\ToM_200ns_20130611.par |
| **customer:** | RKI ZBS 2\| ZBS 6 |
| **measurement date/time:** | 2013-06-13T15:05:30.796+02:00 |
| **path to MS file:** | C:\Users\LaschP\Documents\MATLAB\Microbe MS testdata\ring trial RKI spectra\Sample_10\0_H21\1\1SLin |

##### Detailed quality test report

| **Parameter** | **Noise** | **Baseline error** | **Number of peaks** | **Mean resolving power** | **General Quality** |
|---|---|---|---|---|---|
| **absolut qt results** | 0.63582 | 15.2517 | 25 | 530.1463 | |
| **rank** (out of 24 spectra) | 17 | 21 | 15 | 20 | 20 |
| **quality test result** | 71.4 | 35.7 | 57.1 | 50 | 57.1 |

---

#### # 24 - QUALITY TEST REPORT FOR MALDI-TOF MS SPECTRUM *'2013_06_14_Maren_Stämmler_0010'*

##### Metadata of actual MALDI-ToF test spectrum

| | |
|---|---|
| **genus / species / strain:** | RKI MALDI sample 10, QUANDHIP EQAE ring trial, strains provided by RKI \| ZBS 2 |
| **file id:** | 2013_06_14_Maren_Stämmler_0010 |
| **type:** | Measurement 03 |
| **Bruker ID:** | FCF8970F-FA71-4773-839A-DB9AB0101324 |
| **NCBI ID (primary):** | 1428 |
| **NCBI ID (secondary):** | 1428 |
| **growth time:** | Optimal growth time between 24 - 72h |
| **growth temperature:** | 37°C |
| **growth conditions:** | Optimal aerobic or microaerophilic conditions |
| **growth medium:** | Columbia blood agar (Oxoid), 2nd passage on TSA or Caso agar, harvested by the 2nd passage |
| **sample treatment:** | Sample mixed with 25 mkL 100Â§ TFA, final TFA conc. approx. 80 perc.; approx. 30 Min treatment time; diluted 1:10 (vol); mixed with 1:1 HCCA\|TA2(A) |
| **spores:** | No |
| **concentration:** | Pellet produced by centrifugation (1 x 5 Min, 15,000 rpm) of a 500 mkL cell suspension, gamma ray irradiated (30 kGy) |
| **extra info:** | Bacillus thuringiensis DSM 350: microbial preparation by ZBS 2\| preparation for MALDI-ToF MS: M. Stämmler |
| **calibration standard:** | linear calibration using Escherichia coli DSM 3871 |
| **measurement method:** | D:\Methods\flexControlMethods\MaierMethods\ToM_200ns_20130611.par |
| **customer:** | RKI ZBS 2\| ZBS 6 |
| **measurement date/time:** | 2013-06-14T09:51:48.343+02:00 |
| **path to MS file:** | C:\Users\LaschP\Documents\MATLAB\Microbe MS testdata\ring trial RKI spectra\Sample_10\0_H22\1\1SLin |

##### Detailed quality test report

| **Parameter** | **Noise** | **Baseline error** | **Number of peaks** | **Mean resolving power** | **General Quality** |
|---|---|---|---|---|---|
| **absolut qt results** | 0.71045 | 35.1452 | 19 | 440.4483 | |
| **rank** (out of 24 spectra) | 23 | 24 | 22 | 24 | 24 |
| **quality test result** | 71.4 | 7.1 | 42.9 | 35.7 | 45 |

---
