## Supplementary material for "MicrobeMS - A MATLAB Toolbox for Microbial Identification Based on Mass Spectrometry": Identification report of the HPB data set

### MicrobeMS IDENTIFICATION REPORT FROM MICROORGANISM MALDI-TOF MASS SPECTRA - OVERVIEW

#### Database search parameters

| | |
|---|---|
| **database name:** | 230306_ZENODO_30Peaks_0.75.pkf |
| **date / time of analysis:** | 07-Jan-2026 / 12:32:31 |
| **# of database entries:** | 1599 |
| **distance method:** | Pearson scaling |
| **use weightings:** | distances obtained using peak intensities, weighting factor: 0.1 |
| **vary calibration parameters:** | the complete set of calibration parameters was applied |
| **# of variations:** | 125 ⇒ $(2\times2+1)^3$ variations of calibration parameters |
| **rel width of m/z intervals (ppm):** | 800 |
| **calib variation range factor:** | 3 |
| **m/z range:** | [1975 - 13162.5] |
| **peak number ratio corr factor:** | 1 |
| **log score thresholds:** | [2.5 - 2.75] |
| **thresholds of rank list analyses:** | [400 - 600] |
| **mean score of rank list analyses:** | 720.5958 |

#### Short identification report for MALDI-ToF test spectra

| Test spectra: ID's and quality test results | Summaries of score ranking lists | Score | Top entries of score ranking lists | Score |
|---|---|---|---|---|
| **# 1** - 2013_06_13_Maren_Stämmler_0001 | **1 -** Burkholderia pseudomallei | 786.5 | **1 -** Burkholderia pseudomallei A335-1 | 701.1 |
| RKI MALDI sample 01, QUANDHIP EQAE rin ... | **2 -** Burkholderia mallei | 26.3 | **2 -** Burkholderia pseudomallei PITT 521 041206RR1059 | 689.2 |
| 78.6 78.6 64.3 71.4 71.8 | **3 -** Burkholderia thailandensis | 13.3 | **3 -** Burkholderia pseudomallei 03 04448 251109RR8920 | 660.9 |
| **# 2** - 2013_06_14_Maren_Stämmler_0001 | **1 -** Burkholderia pseudomallei | 832.7 | **1 -** Burkholderia pseudomallei PITT 521 041206RR1059 | 782.9 |
| RKI MALDI sample 01, QUANDHIP EQAE rin ... | **2 -** Burkholderia mallei | 27.9 | **2 -** Burkholderia pseudomallei A335-1 | 728.6 |
| 78.6 78.6 50 71.4 66.1 | **3 -** n/a | 0 | **3 -** Burkholderia pseudomallei 03 04448 251109RR8920 | 718.6 |
| **# 3** - 2013_06_13_Maren_Stämmler_0002 | **1 -** Francisella tularensis ssp. holarctica | 880.2 | **1 -** Francisella tularensis ssp. holarctica 06T0001 | 830.3 |
| RKI MALDI sample 02, QUANDHIP EQAE rin ... | **2 -** Francisella tularensis ssp. tularensis | 1.9 | **2 -** Francisella tularensis ssp. holarctica A 1935-1 | 737.5 |
| 78.6 64.3 64.3 64.3 68.6 | **3 -** Francisella tularensis ssp. mediasiatica | 0.9 | **3 -** Francisella tularensis ssp. holarctica Gaisky | 717 |
| **# 4** - 2013_06_14_Maren_Stämmler_0002 | **1 -** Francisella tularensis ssp. holarctica | 880.9 | **1 -** Francisella tularensis ssp. holarctica 06T0001 | 799.4 |
| RKI MALDI sample 02, QUANDHIP EQAE rin ... | **2 -** Francisella tularensis ssp. tularensis | 1.9 | **2 -** Francisella tularensis ssp. holarctica A 1935-3 | 766.3 |
| 71.4 64.3 57.1 64.3 63.6 | **3 -** Francisella tularensis ssp. mediasiatica | 0.9 | **3 -** Francisella tularensis ssp. holarctica A 1935-1 | 737.7 |
| **# 5** - 2013_06_13_Maren_Stämmler_0003 | **1 -** Brucella canis | 696.8 | **1 -** Brucella canis A183-5 | 801.2 |
| RKI MALDI sample 03, QUANDHIP EQAE rin ... | **2 -** Brucella abortus | 212.3 | **2 -** Brucella canis A446-3 | 728.4 |
| 85.7 78.6 57.1 71.4 71.1 | **3 -** Brucella melitensis | 3.2 | **3 -** Brucella abortus A104-10 | 720.9 |
| **# 6** - 2013_06_14_Maren_Stämmler_0003 | **1 -** Brucella canis | 616.4 | **1 -** Brucella canis A183-5 | 801.9 |
| RKI MALDI sample 03, QUANDHIP EQAE rin ... | **2 -** Brucella abortus | 290.1 | **2 -** Brucella abortus A104-10 | 747.8 |
| 78.6 71.4 50 78.6 66.1 | **3 -** Brucella melitensis | 6.7 | **3 -** Brucella canis A446-3 | 728.3 |
| **# 7** - 2013_06_17_Maren_Stämmler_0001 | **1 -** Bacillus anthracis | 870.3 | **1 -** Bacillus anthracis unknown origin (A18, Beyer) | 756.2 |
| RKI MALDI sample 04, QUANDHIP EQAE rin ... | **2 -** Bacillus cereus | 0 | **2 -** Bacillus anthracis unknown origin (A61, Beyer) | 755.8 |
| 78.6 50 64.3 57.1 65.4 | **3 -** n/a | 0 | **3 -** Bacillus anthracis unknown origin (A27, Beyer) | 726.7 |
| **# 8** - 2013_06_14_Maren_Stämmler_0004 | **1 -** Bacillus anthracis | 867.6 | **1 -** Bacillus anthracis unknown origin (A10, Beyer) | 786.6 |
| RKI MALDI sample 04, QUANDHIP EQAE rin ... | **2 -** Bacillus cereus | 14.1 | **2 -** Bacillus anthracis unknown origin (A7, Beyer) | 750.2 |
| 57.1 28.6 42.9 50 46.1 | **3 -** n/a | 0 | **3 -** Bacillus anthracis unknown origin (A61, Beyer) | 749.4 |

| | | | | |
|---|---|---|---|---|
| # 9 - 2013_06_13_Maren_Stämmler_0004 | 1 - Bacillus anthracis | 847.7 | 1 - Bacillus anthracis ATCC 4229 (RKI) | 754.6 |
| RKI MALDI sample 04, QUANDHIP EQAE rin ... | 2 - Bacillus cereus | 3.4 | 2 - Bacillus anthracis unknown origin (A27, Beyer) | 692.4 |
| 78.6 50 71.4 57.1 68.2 | 3 - Bacillus cereus s.l. | 0 | 3 - Bacillus anthracis unknown origin (A61, Beyer) | 691.3 |
| # 10 - 2013_06_13_Maren_Stämmler_0005 | 1 - Ochrobactrum anthropi | 828.8 | 1 - Ochrobactrum anthropi DSM 20150, ATCC 11425 | 697.9 |
| RKI MALDI sample 05, QUANDHIP EQAE rin ... | 2 - Brucella intermedia | 128.5 | 2 - Ochrobactrum anthropi Ring trial A-269 08 A | 638.2 |
| 78.6 71.4 64.3 71.4 70.7 | 3 - Burkholderia plantarii | 50.2 | 3 - Brucella intermedia A1309 | 347.5 |
| # 11 - 2013_06_14_Maren_Stämmler_0005 | 1 - Ochrobactrum anthropi | 827.6 | 1 - Ochrobactrum anthropi DSM 20150, ATCC 11425 | 696.7 |
| RKI MALDI sample 05, QUANDHIP EQAE rin ... | 2 - Brucella intermedia | 128.3 | 2 - Ochrobactrum anthropi Ring trial A-269 08 A | 635.4 |
| 71.4 50 78.6 64.3 70 | 3 - Burkholderia plantarii | 49.7 | 3 - Brucella intermedia A1309 | 346.8 |
| # 12 - 2013_06_13_Maren_Stämmler_0006 | 1 - Yersinia pseudotuberculosis | 748.8 | 1 - Yersinia pseudotuberculosis 29827 | 698.9 |
| RKI MALDI sample 06, QUANDHIP EQAE rin ... | 2 - Yersinia similis | 67.7 | 2 - Yersinia pseudotuberculosis VI | 694.5 |
| 78.6 71.4 78.6 64.3 75.4 | 3 - Yersinia pestis | 44.6 | 3 - Yersinia pseudotuberculosis 25743 | 674 |
| # 13 - 2013_06_14_Maren_Stämmler_0006 | 1 - Yersinia pseudotuberculosis | 743.8 | 1 - Yersinia pseudotuberculosis VI | 692.9 |
| RKI MALDI sample 06, QUANDHIP EQAE rin ... | 2 - Yersinia similis | 67.8 | 2 - Yersinia pseudotuberculosis 25743 | 672.2 |
| 78.6 71.4 71.4 64.3 72.5 | 3 - Yersinia pestis | 44 | 3 - Yersinia pseudotuberculosis 29490 | 672.1 |
| # 14 - 2013_06_13_Maren_Stämmler_0007 | 1 - Burkholderia mallei | 707.5 | 1 - Burkholderia mallei Dubai 7 | 626.7 |
| RKI MALDI sample 07, QUANDHIP EQAE rin ... | 2 - Burkholderia pseudomallei | 83.7 | 2 - Burkholderia mallei Dubai 7 240609RR5318 | 621.2 |
| 78.6 64.3 71.4 71.4 72.5 | 3 - n/a | 0 | 3 - Burkholderia mallei ATCC 23344 251109RR8925 | 620.2 |
| # 15 - 2013_06_14_Maren_Stämmler_0007 | 1 - Burkholderia mallei | 482.5 | 1 - Burkholderia pseudomallei PITT 5691 041206RR1064 | 660.2 |
| RKI MALDI sample 07, QUANDHIP EQAE rin ... | 2 - Burkholderia pseudomallei | 324.4 | 2 - Burkholderia mallei Dubai 7 | 655.4 |
| 71.4 64.3 78.6 71.4 73.2 | 3 - Burkholderia thailandensis | 0 | 3 - Burkholderia mallei Dubai 7 240609RR5318 | 617.9 |
| # 16 - 2013_06_17_Maren_Stämmler_0002 | 1 - Burkholderia thailandensis | 798.7 | 1 - Burkholderia thailandensis E125 | 654.7 |
| RKI MALDI sample 08, QUANDHIP EQAE rin ... | 2 - Burkholderia oklahomensis | 0 | 2 - Burkholderia thailandensis DSM 13276 | 620.4 |
| 71.4 50 57.1 57.1 60.4 | 3 - Burkholderia pseudomallei | 0 | 3 - Burkholderia thailandensis E184 | 617.6 |
| # 17 - 2013_06_13_Maren_Stämmler_0008 | 1 - Burkholderia thailandensis | 879.2 | 1 - Burkholderia thailandensis DSM 13276 | 783.8 |
| RKI MALDI sample 08, QUANDHIP EQAE rin ... | 2 - Burkholderia pseudomallei | 0 | 2 - Burkholderia thailandensis E184 | 752.6 |
| 78.6 71.4 71.4 71.4 73.6 | 3 - Burkholderia mallei | 0 | 3 - Burkholderia thailandensis E131 | 750.8 |
| # 18 - 2013_06_14_Maren_Stämmler_0008 | 1 - Burkholderia thailandensis | 878.4 | 1 - Burkholderia thailandensis E131 | 774.6 |
| RKI MALDI sample 08, QUANDHIP EQAE rin ... | 2 - Burkholderia mallei | 0 | 2 - Burkholderia thailandensis E125 | 748.6 |
| 71.4 50 71.4 64.3 67.1 | 3 - Burkholderia pseudomallei | 0 | 3 - Burkholderia thailandensis E184 | 746.5 |
| # 19 - 2013_06_17_Maren_Stämmler_0003 | 1 - Yersinia pestis | 396.4 | 1 - Yersinia pestis O3-01501 | 320.5 |
| RKI MALDI sample 09, QUANDHIP EQAE rin ... | 2 - Yersinia pseudotuberculosis | 72.6 | 2 - Yersinia pestis O3-01500 | 313.2 |
| 78.6 57.1 21.4 42.9 47.1 | 3 - Bacillus anthracis | 5.8 | 3 - Yersinia pestis CCUG EV 76, CCUG 32133 | 304.2 |
| # 20 - 2013_06_13_Maren_Stämmler_0009 | 1 - Yersinia pestis | 432.2 | 1 - Yersinia pestis O3-01501 | 362.2 |
| RKI MALDI sample 09, QUANDHIP EQAE rin ... | 2 - Yersinia similis | 73.5 | 2 - Yersinia pestis CCUG EV 76, CCUG 32133 | 344.5 |
| 78.6 78.6 28.6 42.9 53.2 | 3 - Yersinia pseudotuberculosis | 48.6 | 3 - Yersinia pestis O3-01500 | 325.4 |
| # 21 - 2013_06_14_Maren_Stämmler_0009 | 1 - Yersinia similis | 378.7 | 1 - Yersinia similis DSM 18211, LMG 23763, CCUG 52882 | 413.4 |

| | | | | |
|---|---|---|---|---|
| RKI MALDI sample 09, QUANDHIP EQAE rin ... | 2 - Yersinia pestis | 240.1 | 2 - Yersinia pestis CCUG EV 76, CCUG 32133 | 402 |
| 78.6 64.3 50 50 60.7 | 3 - Yersinia pseudotuberculosis | 114.5 | 3 - Yersinia pestis O3-01501 | 387.7 |
| **# 22** - 2013_06_17_Maren_Stämmler_0004 | 1 - Bacillus thuringiensis | 763.2 | 1 - Bacillus thuringiensis DSM 2046 (B188, Beyer) | 647.5 |
| RKI MALDI sample 10, QUANDHIP EQAE rin ... | 2 - Bacillus cereus | 5.8 | 2 - Bacillus thuringiensis DSM 350 | 589.8 |
| 78.6 28.6 57.1 50 58.2 | 3 - Bacillus anthracis | 1.7 | 3 - Bacillus thuringiensis DSM 2046 | 555.4 |
| **# 23** - 2013_06_13_Maren_Stämmler_0010 | 1 - Bacillus thuringiensis | 789.3 | 1 - Bacillus thuringiensis DSM 2046 (B188, Beyer) | 649.3 |
| RKI MALDI sample 10, QUANDHIP EQAE rin ... | 2 - Bacillus cereus | 5.6 | 2 - Bacillus thuringiensis DSM 6890 | 648.9 |
| 71.4 35.7 57.1 50 57.1 | 3 - Bacillus cereus | 1.3 | 3 - Bacillus thuringiensis DSM 350 | 628.1 |
| **# 24** - 2013_06_14_Maren_Stämmler_0010 | 1 - Bacillus thuringiensis | 738.8 | 1 - Bacillus thuringiensis DSM 350 | 613.1 |
| RKI MALDI sample 10, QUANDHIP EQAE rin ... | 2 - Bacillus anthracis | 3 | 2 - Bacillus thuringiensis DSM 6890 | 544.7 |
| 71.4 7.1 42.9 35.7 45 | 3 - Bacillus toyonensis | 1.1 | 3 - Bacillus thuringiensis DSM 2046 (B188, Beyer) | 539.5 |

---

#### # 1 - IDENTIFICATION REPORT FOR MALDI-TOF MS SPECTRUM *'2013_06_13_Maren_Stämmler_0001'*

##### Metadata of actual MALDI-ToF test spectrum

| | |
|---|---|
| **genus / species / strain:** | RKI MALDI sample 01, QUANDHIP EQAE ring trial, strains provided by RKI \| ZBS 2 |
| **file id:** | 2013_06_13_Maren_Stämmler_0001 |
| **type:** | Measurement 01 |
| **Bruker ID:** | 732DF236-D7F4-4EE7-99BF-FD7E225F3A6F |
| **NCBI ID (primary):** | 28450 |
| **NCBI ID (secondary):** | 28450 |
| **growth time:** | Optimal growth time between 24 - 72h |
| **growth temperature:** | 37°C |
| **growth conditions:** | Optimal aerobic or microaerophilic conditions |
| **growth medium:** | Columbia blood agar (Oxoid), 2nd passage on TSA or Caso agar, harvested by the 2nd passage |
| **sample treatment:** | Sample mixed with 20 mkL 100Â§ TFA, final TFA conc. approx. 80 perc.; approx. 30 Min treatment time; diluted 1:10 (vol); mixed with 1:1 HCCA\|TA2(A) |
| **spores:** | No |
| **concentration:** | Pellet produced by centrifugation (1 x 5 Min, 15,000 rpm) of a 500 mkL cell suspension, gamma ray irradiated (30 kGy) |
| **extra info:** | Burkholderia pseudomallei A101-10: microbial preparation by ZBS 2\| preparation for MALDI-ToF MS: M. Stämmler |
| **calibration standard:** | linear calibration using Escherichia coli DSM 3871 |
| **measurement method:** | D:\Methods\flexControlMethods\MaierMethods\ToM_200ns_20130611.par |
| **customer:** | RKI ZBS 2\| ZBS 6 |
| **measurement date/time:** | 2013-06-13T13:52:45.062+02:00 |
| **path to MS file:** | C:\Users\LaschP\Documents\MATLAB\Microbe MS testdata\ring trial RKI spectra\Sample_01\0_H3\1\1SLin |

##### Identification results: analysis of score ranking list

| No. | Genus/Species | Score | Log Score | UniProt Identifier | No. of strains in DB |
|---|---|---|---|---|---|
| 1 | Burkholderia pseudomallei | 786.5474 | 2.8957 | 28450 | 20 |
| 2 | Burkholderia mallei | 26.2781 | 1.4196 | 13373 | 33 |
| 3 | Burkholderia thailandensis | 13.3201 | 1.1245 | 57975 | 15 |

##### Score ranking list: best matches with test spectrum

| No. | Genus/Species/Strain | Score | Log Score | Spectrum identifier | Customer, or UniprotKB link |
|---|---|---|---|---|---|
| 1 | Burkholderia pseudomallei A335-1 | 701.1359 | 2.8458 | dbspec-03-Mar-2023-12-51-15.574 | RKI ZBS2 |
| 2 | Burkholderia pseudomallei PITT 521 041206RR1059 | 689.1862 | 2.8383 | dbspec-03-Mar-2023-12-52-34.948 | Dr. Tomaso (FLI Jena) |
| 3 | Burkholderia pseudomallei 03 04448 251109RR8920 | 660.9296 | 2.8202 | dbspec-03-Mar-2023-12-50-25.2 | Dr. Tomaso (FLI Jena) |
| 4 | Burkholderia pseudomallei Heckeshorn 041206RR1061 | 625.2984 | 2.7961 | dbspec-03-Mar-2023-12-51-45.996 | Dr. Tomaso (FLI Jena) |
| 5 | Burkholderia pseudomallei 03 04450 060406RR0740 | 589.3296 | 2.7704 | dbspec-03-Mar-2023-12-50-45.809 | Dr. Tomaso (FLI Jena) |
| 6 | Burkholderia mallei ATCC 23344 300102RR0118 | 583.2784 | 2.7659 | dbspec-03-Mar-2023-12-41-29.653 | Dr. Tomaso (FLI Jena) |
| 7 | Burkholderia thailandensis E153 | 565.0493 | 2.7521 | dbspec-03-Mar-2023-12-58-32.096 | RKI ZBS6 |
| 8 | Burkholderia pseudomallei NCTC 1688 041206RR1062 | 559.2228 | 2.7476 | dbspec-03-Mar-2023-12-52-10.62 | Dr. Tomaso (FLI Jena) |
| 9 | Burkholderia mallei Zagreb 080304RR0090 | 557.5966 | 2.7463 | dbspec-03-Mar-2023-12-40-32.655 | Dr. Tomaso (FLI Jena) |
| 10 | Burkholderia thailandensis 090804RR0288 135 | 556.9761 | 2.7458 | dbspec-03-Mar-2023-12-57-39.41 | Dr. Tomaso (FLI Jena) |
| 11 | Burkholderia mallei BfR M1 290103RR0043 | 555.3814 | 2.7446 | dbspec-03-Mar-2023-12-44-39.196 | Dr. Tomaso (FLI Jena) |
| 12 | Burkholderia pseudomallei PITT 5691 041206RR1064 | 553.3193 | 2.743 | dbspec-03-Mar-2023-12-52-51.791 | Dr. Tomaso (FLI Jena) |
| 13 | Burkholderia mallei Dubai 7 240609RR5318 | 551.0728 | 2.7412 | dbspec-03-Mar-2023-12-39-40.093 | Dr. Tomaso (FLI Jena) |

| | | | | | |
|---|---|---|---|---|---|
| 14 | Burkholderia pseudomallei 3463 | 541.8354 | 2.7339 | dbspec-03-Mar-2023-12-51-39.871 | RKI ZBS6 |
| 15 | Burkholderia thailandensis E143 | 534.5962 | 2.728 | dbspec-03-Mar-2023-12-58-28.784 | RKI ZBS6 |
| 16 | Burkholderia mallei BfR 242 041206RR1050 | 533.3423 | 2.727 | dbspec-03-Mar-2023-12-42-38.496 | Dr. Tomaso (FLI Jena) |
| 17 | Burkholderia mallei BfR M2 041206RR1057 | 529.0283 | 2.7235 | dbspec-03-Mar-2023-12-44-57.446 | Dr. Tomaso (FLI Jena) |
| 18 | Burkholderia mallei NCTC 10247 041206RR1056 | 528.5896 | 2.7231 | dbspec-03-Mar-2023-12-46-40.553 | Dr. Tomaso (FLI Jena) |
| 19 | Burkholderia mallei NCTC 10230 041206RR1052 | 528.4998 | 2.723 | dbspec-03-Mar-2023-12-46-16.366 | Dr. Tomaso (FLI Jena) |
| 20 | Burkholderia mallei EQADeBa | 528.4849 | 2.723 | dbspec-03-Mar-2023-12-45-53.726 | Dr. Tomaso (FLI Jena) |

---

#### # 2 - IDENTIFICATION REPORT FOR MALDI-TOF MS SPECTRUM *'2013_06_14_Maren_Stämmler_0001'*

##### Metadata of actual MALDI-ToF test spectrum

| | |
|---|---|
| **genus / species / strain:** | RKI MALDI sample 01, QUANDHIP EQAE ring trial, strains provided by RKI \| ZBS 2 |
| **file id:** | 2013_06_14_Maren_Stämmler_0001 |
| **type:** | Measurement 02 |
| **Bruker ID:** | C5344CB2-A106-46AA-A51A-64817DBD40F2 |
| **NCBI ID (primary):** | 28450 |
| **NCBI ID (secondary):** | 28450 |
| **growth time:** | Optimal growth time between 24 - 72h |
| **growth temperature:** | 37°C |
| **growth conditions:** | Optimal aerobic or microaerophilic conditions |
| **growth medium:** | Columbia blood agar (Oxoid), 2nd passage on TSA or Caso agar, harvested by the 2nd passage |
| **sample treatment:** | Sample mixed with 20 mkL 100Â§ TFA, final TFA conc. approx. 80 perc.; approx. 30 Min treatment time; diluted 1:10 (vol); mixed with 1:1 HCCA\|TA2(A) |
| **spores:** | No |
| **concentration:** | Pellet produced by centrifugation (1 x 5 Min, 15,000 rpm) of a 500 mkL cell suspension, gamma ray irradiated (30 kGy) |
| **extra info:** | Burkholderia pseudomallei A101-10: microbial preparation by ZBS 2\| preparation for MALDI-ToF MS: M. Stämmler |
| **calibration standard:** | linear calibration using Escherichia coli DSM 3871 |
| **measurement method:** | D:\Methods\flexControlMethods\MaierMethods\ToM_200ns_20130611.par |
| **customer:** | RKI ZBS 2\| ZBS 6 |
| **measurement date/time:** | 2013-06-14T08:27:14.015+02:00 |
| **path to MS file:** | C:\Users\LaschP\Documents\MATLAB\Microbe MS testdata\ring trial RKI spectra\Sample_01\0_H4\1\1SLin |

##### Identification results: analysis of score ranking list

| No. | Genus/Species | Score | Log Score | UniProt Identifier | No. of strains in DB |
|---|---|---|---|---|---|
| 1 | Burkholderia pseudomallei | 832.7202 | 2.9205 | 28450 | 20 |
| 2 | Burkholderia mallei | 27.8945 | 1.4455 | 13373 | 33 |
| 3 | n/a | 0 | -Inf | | |

##### Score ranking list: best matches with test spectrum

| No. | Genus/Species/Strain | Score | Log Score | Spectrum identifier | Customer, or UniprotKB link |
|---|---|---|---|---|---|
| 1 | Burkholderia pseudomallei PITT 521 041206RR1059 | 782.9115 | 2.8937 | dbspec-03-Mar-2023-12-52-34.948 | Dr. Tomaso (FLI Jena) |
| 2 | Burkholderia pseudomallei A335-1 | 728.6319 | 2.8625 | dbspec-03-Mar-2023-12-51-15.574 | RKI ZBS2 |
| 3 | Burkholderia pseudomallei 03 04448 251109RR8920 | 718.5748 | 2.8565 | dbspec-03-Mar-2023-12-50-25.2 | Dr. Tomaso (FLI Jena) |
| 4 | Burkholderia pseudomallei Heckeshorn 041206RR1061 | 691.0637 | 2.8395 | dbspec-03-Mar-2023-12-51-45.996 | Dr. Tomaso (FLI Jena) |
| 5 | Burkholderia pseudomallei NCTC 1688 041206RR1062 | 584.1842 | 2.7665 | dbspec-03-Mar-2023-12-52-10.62 | Dr. Tomaso (FLI Jena) |
| 6 | Burkholderia mallei ATCC 23344 300102RR0118 | 582.9704 | 2.7656 | dbspec-03-Mar-2023-12-41-29.653 | Dr. Tomaso (FLI Jena) |
| 7 | Burkholderia pseudomallei 03 04450 060406RR0740 | 581.2193 | 2.7643 | dbspec-03-Mar-2023-12-50-45.809 | Dr. Tomaso (FLI Jena) |
| 8 | Burkholderia pseudomallei 3463 | 569.9087 | 2.7558 | dbspec-03-Mar-2023-12-51-39.871 | RKI ZBS6 |
| 9 | Burkholderia mallei BfR 242 041206RR1050 | 562.7146 | 2.7503 | dbspec-03-Mar-2023-12-42-38.496 | Dr. Tomaso (FLI Jena) |
| 10 | Burkholderia mallei Zagreb 080304RR0090 | 558.3551 | 2.7469 | dbspec-03-Mar-2023-12-40-32.655 | Dr. Tomaso (FLI Jena) |
| 11 | Burkholderia mallei NCTC 10230 041206RR1052 | 557.128 | 2.746 | dbspec-03-Mar-2023-12-46-16.366 | Dr. Tomaso (FLI Jena) |
| 12 | Burkholderia mallei BfR M2 041206RR1057 | 557.0078 | 2.7459 | dbspec-03-Mar-2023-12-44-57.446 | Dr. Tomaso (FLI Jena) |
| 13 | Burkholderia mallei Bogor 211101RR0419 | 556.4357 | 2.7454 | dbspec-03-Mar-2023-12-39-20.813 | Dr. Tomaso (FLI Jena) |
| 14 | Burkholderia mallei BfR 34 281002RR0518 | 556.2381 | 2.7453 | dbspec-03-Mar-2023-12-43-53.666 | Dr. Tomaso (FLI Jena) |
| 15 | Burkholderia mallei BfR M1 290103RR0043 | 555.2716 | 2.7445 | dbspec-03-Mar-2023-12-44-39.196 | Dr. Tomaso (FLI Jena) |
| 16 | Burkholderia pseudomallei CCUG 15648 | 553.2692 | 2.7429 | dbspec-03-Mar-2023-12-51-24.84 | RKI ZBS6 |
| 17 | Burkholderia mallei BfR 237 061102RR0551 | 551.5972 | 2.7416 | dbspec-03-Mar-2023-12-42-14.512 | Dr. Tomaso (FLI Jena) |
| 18 | Burkholderia mallei Dubai 7 240609RR5318 | 550.9204 | 2.7411 | dbspec-03-Mar-2023-12-39-40.093 | Dr. Tomaso (FLI Jena) |
| 19 | Burkholderia mallei ATCC 23344 251109RR8925 | 550.4647 | 2.7407 | dbspec-03-Mar-2023-12-41-0.185 | Dr. Tomaso (FLI Jena) |
| 20 | Burkholderia pseudomallei PITT 5691 041206RR1064 | 550.0876 | 2.7404 | dbspec-03-Mar-2023-12-52-51.791 | Dr. Tomaso (FLI Jena) |

---

#### # 3 - IDENTIFICATION REPORT FOR MALDI-TOF MS SPECTRUM *'2013_06_13_Maren_Stämmler_0002'*

##### Metadata of actual MALDI-ToF test spectrum

| | |
|---|---|
| **genus / species / strain:** | RKI MALDI sample 02, QUANDHIP EQAE ring trial, strains provided by RKI \| ZBS 2 |

| | |
|---|---|
| **file id:** | 2013_06_13_Maren_Stämmler_0002 |
| **type:** | Measurement 01 |
| **Bruker ID:** | B1409699-0EFE-4C28-AAD4-FB638EF4774B |
| **NCBI ID (primary):** | 119857 |
| **NCBI ID (secondary):** | 119857 |
| **growth time:** | Optimal growth time between 24 - 72h |
| **growth temperature:** | 37°C |
| **growth conditions:** | Optimal aerobic or microaerophilic conditions |
| **growth medium:** | Heart cysteine agar (HCA) |
| **sample treatment:** | Sample mixed with 20 mkL 100Â§ TFA, final TFA conc. approx. 80 perc.; approx. 30 Min treatment time; diluted 1:10 (vol); mixed with 1:1 HCCA\|TA2(A) |
| **spores:** | No |
| **concentration:** | Pellet produced by centrifugation (1 x 5 Min, 15,000 rpm) of a 500 mkL cell suspension, gamma ray irradiated (30 kGy) |
| **extra info:** | Francisella tularensis ssp. holarctica Ft 32: microbial preparation by ZBS 2\| preparation for MALDI-ToF MS: M. Stämmler |
| **calibration standard:** | linear calibration using Escherichia coli DSM 3871 |
| **measurement method:** | D:\Methods\flexControlMethods\MaierMethods\ToM_200ns_20130611.par |
| **customer:** | RKI ZBS 2\| ZBS 6 |
| **measurement date/time:** | 2013-06-13T14:04:41.578+02:00 |
| **path to MS file:** | C:\Users\LaschP\Documents\MATLAB\Microbe MS testdata\ring trial RKI spectra\Sample_02\0_H5\1\1SLin |

#### Identification results: analysis of score ranking list

| No. | Genus/Species | Score | Log Score | UniProt Identifier | No. of strains in DB |
|---|---|---|---|---|---|
| 1 | Francisella tularensis ssp. holarctica | 880.1908 | 2.9446 | 119857 | 9 |
| 2 | Francisella tularensis ssp. tularensis | 1.872 | 0.27231 | 119856 | 1 |
| 3 | Francisella tularensis ssp. mediasiatica | 0.89877 | -0.046353 | 135248 | 3 |
| 4 | Francisella tularensis ssp. novicida | 0.020134 | -1.6961 | 264 | 4 |
| 5 | Morganella morganii ssp. sibonii | 0.00095824 | -3.0185 | 180435 | 2 |

#### Score ranking list: best matches with test spectrum

| No. | Genus/Species/Strain | Score | Log Score | Spectrum identifier | Customer, or UniprotKB link |
|---|---|---|---|---|---|
| 1 | Francisella tularensis ssp. holarctica 06T0001 | 830.3116 | 2.9192 | dbspec-03-Mar-2023-13-13-18.249 | Dr. Tomaso (FLI Jena) |
| 2 | Francisella tularensis ssp. holarctica A 1935-1 | 737.4997 | 2.8678 | dbspec-03-Mar-2023-13-13-54.92 | RKI ZBS2\|ZBS6 |
| 3 | Francisella tularensis ssp. holarctica Gaisky | 716.9505 | 2.8555 | dbspec-03-Mar-2023-13-14-37.388 | Dr. Tomaso (FLI Jena) |
| 4 | Francisella tularensis ssp. holarctica A 1935-3 | 709.8789 | 2.8512 | dbspec-03-Mar-2023-13-14-6.748 | RKI ZBS2\|ZBS6 |
| 5 | Francisella tularensis ssp. holarctica W 2903 | 688.1642 | 2.8377 | dbspec-03-Mar-2023-13-16-16.057 | Dr. Tomaso (FLI Jena) |
| 6 | Francisella tularensis ssp. holarctica Ft32, FSC 200 | 684.7799 | 2.8356 | dbspec-03-Mar-2023-13-14-25.403 | RKI ZBS2\|ZBS6 |
| 7 | Francisella tularensis ssp. holarctica LVS | 654.9771 | 2.8162 | dbspec-03-Mar-2023-13-15-8.59 | Dr. Tomaso (FLI Jena) |
| 8 | Francisella tularensis ssp. holarctica W 1468 | 615.0531 | 2.7889 | dbspec-03-Mar-2023-13-15-53.136 | Dr. Tomaso (FLI Jena) |
| 9 | Francisella tularensis ssp. holarctica A466-1 | 551.0347 | 2.7412 | dbspec-03-Mar-2023-13-14-18.935 | RKI ZBS2\|ZBS6 |
| 10 | Francisella tularensis ssp. tularensis FSC 237 | 459.9171 | 2.6627 | dbspec-03-Mar-2023-13-19-13.038 | Dr. Tomaso (FLI Jena) |
| 11 | Francisella tularensis ssp. mediasiatica Ft31, FSC 148 | 453.4018 | 2.6565 | dbspec-03-Mar-2023-13-17-43.993 | RKI ZBS2\|ZBS6 |
| 12 | Francisella tularensis ssp. mediasiatica F063 | 332.5212 | 2.5218 | dbspec-03-Mar-2023-13-16-42.26 | Dr. Tomaso (FLI Jena) |
| 13 | Francisella tularensis ssp. mediasiatica F064 | 331.2949 | 2.5202 | dbspec-03-Mar-2023-13-17-12.087 | Dr. Tomaso (FLI Jena) |
| 14 | Francisella tularensis ssp. novicida F048 | 287.4028 | 2.4585 | dbspec-03-Mar-2023-13-17-53.133 | Dr. Tomaso (FLI Jena) |
| 15 | Francisella tularensis ssp. novicida Ft26 | 284.6501 | 2.4543 | dbspec-03-Mar-2023-13-19-6.522 | Dr. Tomaso (FLI Jena) |
| 16 | Morganella morganii ssp. sibonii DSM 6675 | 256.751 | 2.4095 | dbspec-03-Mar-2023-13-25-29.858 | RKI ZBS6 |
| 17 | Francisella tularensis ssp. novicida F059 | 256.2705 | 2.4087 | dbspec-03-Mar-2023-13-18-26.961 | Dr. Tomaso (FLI Jena) |
| 18 | Klebsiella aerogenes NM 20 | 239.8525 | 2.3799 | dbspec-03-Mar-2023-13-20-43.239 | RKI ZBS6 |
| 19 | Klebsiella oxytoca CB4063 | 238.4676 | 2.3774 | dbspec-03-Mar-2023-13-21-15.707 | RKI ZBS2\|ZBS6 |
| 20 | Morganella morganii ssp. morganii DSM 30164 | 221.0526 | 2.3445 | dbspec-03-Mar-2023-13-25-20.342 | RKI ZBS6 |

---

### # 4 - IDENTIFICATION REPORT FOR MALDI-TOF MS SPECTRUM *'2013_06_14_Maren_Stämmler_0002'*

#### Metadata of actual MALDI-ToF test spectrum

| | |
|---|---|
| **genus / species / strain:** | RKI MALDI sample 02, QUANDHIP EQAE ring trial, strains provided by RKI \| ZBS 2 |
| **file id:** | 2013_06_14_Maren_Stämmler_0002 |
| **type:** | Measurement 02 |
| **Bruker ID:** | 4F9079BF-2237-4052-A736-259FD2ABA28A |
| **NCBI ID (primary):** | 119857 |
| **NCBI ID (secondary):** | 119857 |
| **growth time:** | Optimal growth time between 24 - 72h |
| **growth temperature:** | 37°C |
| **growth conditions:** | Optimal aerobic or microaerophilic conditions |
| **growth medium:** | Heart cysteine agar (HCA) |
| **sample treatment:** | Sample mixed with 20 mkL 100Â§ TFA, final TFA conc. approx. 80 perc.; approx. 30 Min treatment time; diluted 1:10 (vol); mixed with 1:1 HCCA\|TA2(A) |
| **spores:** | No |
| **concentration:** | Pellet produced by centrifugation (1 x 5 Min, 15,000 rpm) of a 500 mkL cell suspension, gamma ray irradiated (30 kGy) |
| **extra info:** | Francisella tularensis ssp. holarctica Ft 32: microbial preparation by ZBS 2\| preparation for MALDI-ToF MS: M. Stämmler |
| **calibration standard:** | linear calibration using Escherichia coli DSM 3871 |
| **measurement method:** | D:\Methods\flexControlMethods\MaierMethods\ToM_200ns_20130611.par |
| **customer:** | RKI ZBS 2\| ZBS 6 |

| | |
|---|---|
| **measurement date/time:** | 2013-06-14T08:33:44.531+02:00 |
| **path to MS file:** | C:\Users\LaschP\Documents\MATLAB\Microbe MS testdata\ring trial RKI spectra\Sample_02\0_H6\1\1SLin |

#### Identification results: analysis of score ranking list

| No. | Genus/Species | Score | Log Score | UniProt Identifier | No. of strains in DB |
|---|---|---|---|---|---|
| 1 | Francisella tularensis ssp. holarctica | 880.8784 | 2.9449 | 119857 | 9 |
| 2 | Francisella tularensis ssp. tularensis | 1.9367 | 0.28707 | 119856 | 1 |
| 3 | Francisella tularensis ssp. mediasiatica | 0.87553 | -0.057731 | 135248 | 3 |
| 4 | Francisella tularensis ssp. novicida | 0.022062 | -1.6564 | 264 | 4 |
| 5 | Morganella morganii ssp. sibonii | 0.0010417 | -2.9823 | 180435 | 2 |

#### Score ranking list: best matches with test spectrum

| No. | Genus/Species/Strain | Score | Log Score | Spectrum identifier | Customer, or UniprotKB link |
|---|---|---|---|---|---|
| 1 | Francisella tularensis ssp. holarctica 06T0001 | 799.4063 | 2.9028 | dbspec-03-Mar-2023-13-13-18.249 | Dr. Tomaso (FLI Jena) |
| 2 | Francisella tularensis ssp. holarctica A 1935-3 | 766.2873 | 2.8844 | dbspec-03-Mar-2023-13-14-6.748 | RKI ZBS2\|ZBS6 |
| 3 | Francisella tularensis ssp. holarctica A 1935-1 | 737.678 | 2.8679 | dbspec-03-Mar-2023-13-13-54.92 | RKI ZBS2\|ZBS6 |
| 4 | Francisella tularensis ssp. holarctica Gaisky | 713.9006 | 2.8536 | dbspec-03-Mar-2023-13-14-37.388 | Dr. Tomaso (FLI Jena) |
| 5 | Francisella tularensis ssp. holarctica Ft32, FSC 200 | 713.0865 | 2.8531 | dbspec-03-Mar-2023-13-14-25.403 | RKI ZBS2\|ZBS6 |
| 6 | Francisella tularensis ssp. holarctica W 2903 | 680.2516 | 2.8327 | dbspec-03-Mar-2023-13-16-16.057 | Dr. Tomaso (FLI Jena) |
| 7 | Francisella tularensis ssp. holarctica LVS | 650.9548 | 2.8136 | dbspec-03-Mar-2023-13-15-8.59 | Dr. Tomaso (FLI Jena) |
| 8 | Francisella tularensis ssp. holarctica W 1468 | 641.7365 | 2.8074 | dbspec-03-Mar-2023-13-15-53.136 | Dr. Tomaso (FLI Jena) |
| 9 | Francisella tularensis ssp. holarctica A466-1 | 581.6996 | 2.7647 | dbspec-03-Mar-2023-13-14-18.935 | RKI ZBS2\|ZBS6 |
| 10 | Francisella tularensis ssp. tularensis FSC 237 | 484.8172 | 2.6856 | dbspec-03-Mar-2023-13-19-13.038 | Dr. Tomaso (FLI Jena) |
| 11 | Francisella tularensis ssp. mediasiatica Ft31, FSC 148 | 424.0259 | 2.6274 | dbspec-03-Mar-2023-13-17-43.993 | RKI ZBS2\|ZBS6 |
| 12 | Francisella tularensis ssp. mediasiatica F063 | 339.7939 | 2.5312 | dbspec-03-Mar-2023-13-16-42.26 | Dr. Tomaso (FLI Jena) |
| 13 | Francisella tularensis ssp. mediasiatica F064 | 339.5853 | 2.5309 | dbspec-03-Mar-2023-13-17-12.087 | Dr. Tomaso (FLI Jena) |
| 14 | Francisella tularensis ssp. novicida Ft26 | 325.6324 | 2.5127 | dbspec-03-Mar-2023-13-19-6.522 | Dr. Tomaso (FLI Jena) |
| 15 | Francisella tularensis ssp. novicida F059 | 286.4787 | 2.4571 | dbspec-03-Mar-2023-13-18-26.961 | Dr. Tomaso (FLI Jena) |
| 16 | Morganella morganii ssp. sibonii DSM 6675 | 278.7286 | 2.4452 | dbspec-03-Mar-2023-13-25-29.858 | RKI ZBS6 |
| 17 | Francisella tularensis ssp. novicida F048 | 262.6836 | 2.4194 | dbspec-03-Mar-2023-13-17-53.133 | Dr. Tomaso (FLI Jena) |
| 18 | Klebsiella aerogenes NM 20 | 259.8792 | 2.4148 | dbspec-03-Mar-2023-13-20-43.239 | RKI ZBS6 |
| 19 | Francisella hispaniensis DSM 22475, CCUG 58020 | 252.581 | 2.4024 | dbspec-03-Mar-2023-13-12-3.875 | RKI ZBS6 |
| 20 | Morganella morganii ssp. morganii DSM 30164 | 242.634 | 2.385 | dbspec-03-Mar-2023-13-25-20.342 | RKI ZBS6 |

---

### # 5 - IDENTIFICATION REPORT FOR MALDI-TOF MS SPECTRUM *'2013_06_13_Maren_Stämmler_0003'*

#### Metadata of actual MALDI-ToF test spectrum

| | |
|---|---|
| **genus / species / strain:** | RKI MALDI sample 03, QUANDHIP EQAE ring trial, strains provided by RKI \| ZBS 2 |
| **file id:** | 2013_06_13_Maren_Stämmler_0003 |
| **type:** | Measurement 01 |
| **Bruker ID:** | 76182EF3-AD43-4157-A768-0D20D4A2FDF6 |
| **NCBI ID (primary):** | 36855 |
| **NCBI ID (secondary):** | 36855 |
| **growth time:** | Optimal growth time between 24 - 72h |
| **growth temperature:** | 37°C |
| **growth conditions:** | Optimal aerobic or microaerophilic conditions |
| **growth medium:** | Columbia blood agar (Oxoid), 2nd passage on TSA or Caso agar, harvested by the 2nd passage |
| **sample treatment:** | Sample mixed with 20 mkL 100Â§ TFA, final TFA conc. approx. 80 perc.; approx. 30 Min treatment time; diluted 1:10 (vol); mixed with 1:1 HCCA\|TA2(A) |
| **spores:** | No |
| **concentration:** | Pellet produced by centrifugation (1 x 5 Min, 15,000 rpm) of a 500 mkL cell suspension, gamma ray irradiated (30 kGy) |
| **extra info:** | Brucella canis A138-11: microbial preparation by ZBS 2\| preparation for MALDI-ToF MS: M. Stämmler |
| **calibration standard:** | linear calibration using Escherichia coli DSM 3871 |
| **measurement method:** | D:\Methods\flexControlMethods\MaierMethods\ToM_200ns_20130611.par |
| **customer:** | RKI ZBS 2\| ZBS 6 |
| **measurement date/time:** | 2013-06-13T14:11:32.906+02:00 |
| **path to MS file:** | C:\Users\LaschP\Documents\MATLAB\Microbe MS testdata\ring trial RKI spectra\Sample_03\0_H7\1\1SLin |

#### Identification results: analysis of score ranking list

| No. | Genus/Species | Score | Log Score | UniProt Identifier | No. of strains in DB |
|---|---|---|---|---|---|
| 1 | Brucella canis | 696.7732 | 2.8431 | 36855 | 5 |
| 2 | Brucella abortus | 212.2509 | 2.3268 | 235 | 16 |
| 3 | Brucella melitensis | 3.1687 | 0.50088 | 29459 | 9 |
| 4 | Brucella neotomae | 0.0014446 | -2.8403 | 29460 | 3 |
| 5 | Brucella ovis | 0.00017317 | -3.7615 | 236 | 4 |

#### Score ranking list: best matches with test spectrum

| No. | Genus/Species/Strain | Score | Log Score | Spectrum identifier | Customer, or UniprotKB link |
|---|---|---|---|---|---|
| 1 | Brucella canis A183-5 | 801.1668 | 2.9037 | dbspec-03-Mar-2023-12-17-3.16 | RKI ZBS2\|ZBS6 |
| 2 | Brucella canis A446-3 | 728.401 | 2.8624 | dbspec-03-Mar-2023-12-17-9.159 | RKI ZBS2 |
| 3 | Brucella abortus A104-10 | 720.9118 | 2.8579 | dbspec-03-Mar-2023-12-14-18.585 | RKI ZBS2 |
| 4 | Brucella canis R 6 66 03RB0214 3 | 708.3006 | 2.8502 | dbspec-03-Mar-2023-12-17-14.847 | Dr. Tomaso (FLI Jena) |
| 5 | Brucella abortus 9 Strain C68 07RB1203 | 635.1573 | 2.8029 | dbspec-03-Mar-2023-12-13-50.945 | Dr. Tomaso (FLI Jena) |
| 6 | Brucella abortus 5 Strain B3196 07RB1201 | 634.8456 | 2.8027 | dbspec-03-Mar-2023-12-13-0.259 | Dr. Tomaso (FLI Jena) |
| 7 | Brucella abortus 2 Strain 86 8 59 | 629.974 | 2.7993 | dbspec-03-Mar-2023-12-12-15.385 | Dr. Tomaso (FLI Jena) |
| 8 | Brucella canis 04 1 001 29 | 629.2533 | 2.7988 | dbspec-03-Mar-2023-12-16-35.926 | Dr. Tomaso (FLI Jena) |
| 9 | Brucella melitensis A146-13 | 622.1862 | 2.7939 | dbspec-03-Mar-2023-12-21-12.87 | RKI ZBS2 |
| 10 | Brucella abortus S19 | 605.3688 | 2.782 | dbspec-03-Mar-2023-12-14-44.397 | RKI FG14 |
| 11 | Brucella abortus 6 Strain 870 07RB1202 | 604.5702 | 2.7814 | dbspec-03-Mar-2023-12-13-27.164 | Dr. Tomaso (FLI Jena) |
| 12 | Brucella abortus 01 1 001 1 | 603.6456 | 2.7808 | dbspec-03-Mar-2023-12-11-7.98 | Dr. Tomaso (FLI Jena) |
| 13 | Brucella abortus Ring trial A-269 06 A | 599.2489 | 2.7776 | dbspec-03-Mar-2023-12-14-16.194 | RKI ZBS2\|ZBS6 |
| 14 | Brucella canis (BM385) | 589.5199 | 2.7705 | dbspec-03-Mar-2023-12-16-11.379 | TNO Netherlands |
| 15 | Brucella melitensis biovar 2 (BM410) | 583.276 | 2.7659 | dbspec-03-Mar-2023-12-22-36.774 | TNO Netherlands |
| 16 | Brucella neotomae A148-7 | 580.8429 | 2.7641 | dbspec-03-Mar-2023-12-24-20.374 | RKI ZBS2\|ZBS6 |
| 17 | Brucella ovis 63 290 03RB0213 3 | 576.426 | 2.7607 | dbspec-03-Mar-2023-12-25-42.185 | Dr. Tomaso (FLI Jena) |
| 18 | Brucella abortus S19 delta mglA3.14 | 574.9268 | 2.7596 | dbspec-03-Mar-2023-12-14-50.428 | RKI FG14 |
| 19 | Brucella sp. A 1755_25 | 567.501 | 2.754 | dbspec-03-Mar-2023-12-27-17.183 | RKI ZBS6 |
| 20 | Brucella melitensis 02 3 022 17 | 565.8506 | 2.7527 | dbspec-03-Mar-2023-12-20-44.98 | Dr. Tomaso (FLI Jena) |

---

### # 6 - IDENTIFICATION REPORT FOR MALDI-TOF MS SPECTRUM *'2013_06_14_Maren_Stämmler_0003'*

#### Metadata of actual MALDI-ToF test spectrum

| | |
|---|---|
| **genus / species / strain:** | RKI MALDI sample 03, QUANDHIP EQAE ring trial, strains provided by RKI \| ZBS 2 |
| **file id:** | 2013_06_14_Maren_Stämmler_0003 |
| **type:** | Measurement 02 |
| **Bruker ID:** | 22710A07-8E7C-49C6-8EB2-00460D2E50DB |
| **NCBI ID (primary):** | 36855 |
| **NCBI ID (secondary):** | 36855 |
| **growth time:** | Optimal growth time between 24 - 72h |
| **growth temperature:** | 37°C |
| **growth conditions:** | Optimal aerobic or microaerophilic conditions |
| **growth medium:** | Columbia blood agar (Oxoid), 2nd passage on TSA or Caso agar, harvested by the 2nd passage |
| **sample treatment:** | Sample mixed with 20 mkL 100Â§ TFA, final TFA conc. approx. 80 perc.; approx. 30 Min treatment time; diluted 1:10 (vol); mixed with 1:1 HCCA\|TA2(A) |
| **spores:** | No |
| **concentration:** | Pellet produced by centrifugation (1 x 5 Min, 15,000 rpm) of a 500 mkL cell suspension, gamma ray irradiated (30 kGy) |
| **extra info:** | Brucella canis A138-11: microbial preparation by ZBS 2\| preparation for MALDI-ToF MS: M. Stämmler |
| **calibration standard:** | linear calibration using Escherichia coli DSM 3871 |
| **measurement method:** | D:\Methods\flexControlMethods\MaierMethods\ToM_200ns_20130611.par |
| **customer:** | RKI ZBS 2\| ZBS 6 |
| **measurement date/time:** | 2013-06-14T08:42:40.468+02:00 |
| **path to MS file:** | C:\Users\LaschP\Documents\MATLAB\Microbe MS testdata\ring trial RKI spectra\Sample_03\0_H8\1\1SLin |

#### Identification results: analysis of score ranking list

| No. | Genus/Species | Score | Log Score | UniProt Identifier | No. of strains in DB |
|---|---|---|---|---|---|
| 1 | Brucella canis | 616.4006 | 2.7899 | 36855 | 5 |
| 2 | Brucella abortus | 290.0613 | 2.4625 | 235 | 16 |
| 3 | Brucella melitensis | 6.6668 | 0.82392 | 29459 | 9 |
| 4 | Brucella neotomae | 0.10192 | -0.99176 | 29460 | 3 |
| 5 | Brucella sp. | 0.00015691 | -3.8043 | 52132 | 9 |

#### Score ranking list: best matches with test spectrum

| No. | Genus/Species/Strain | Score | Log Score | Spectrum identifier | Customer, or UniprotKB link |
|---|---|---|---|---|---|
| 1 | Brucella canis A183-5 | 801.9215 | 2.9041 | dbspec-03-Mar-2023-12-17-3.16 | RKI ZBS2\|ZBS6 |
| 2 | Brucella abortus A104-10 | 747.8337 | 2.8738 | dbspec-03-Mar-2023-12-14-18.585 | RKI ZBS2 |
| 3 | Brucella canis A446-3 | 728.294 | 2.8623 | dbspec-03-Mar-2023-12-17-9.159 | RKI ZBS2 |
| 4 | Brucella canis R 6 66 03RB0214 3 | 711.3406 | 2.8521 | dbspec-03-Mar-2023-12-17-14.847 | Dr. Tomaso (FLI Jena) |
| 5 | Brucella abortus 9 Strain C68 07RB1203 | 662.5882 | 2.8212 | dbspec-03-Mar-2023-12-13-50.945 | Dr. Tomaso (FLI Jena) |
| 6 | Brucella abortus 5 Strain B3196 07RB1201 | 662.2602 | 2.821 | dbspec-03-Mar-2023-12-13-0.259 | Dr. Tomaso (FLI Jena) |
| 7 | Brucella canis 04 1 001 29 | 658.5238 | 2.8186 | dbspec-03-Mar-2023-12-16-35.926 | Dr. Tomaso (FLI Jena) |
| 8 | Brucella melitensis A146-13 | 652.0402 | 2.8143 | dbspec-03-Mar-2023-12-21-12.87 | RKI ZBS2 |
| 9 | Brucella abortus S19 | 634.631 | 2.8025 | dbspec-03-Mar-2023-12-14-44.397 | RKI FG14 |

| | | | | | |
|---|---|---|---|---|---|
| 10 | Brucella abortus 6 Strain 870 07RB1202 | 631.8799 | 2.8006 | dbspec-03-Mar-2023-12-13-27.164 | Dr. Tomaso (FLI Jena) |
| 11 | Brucella abortus 01 1 001 1 | 631.3795 | 2.8003 | dbspec-03-Mar-2023-12-11-7.98 | Dr. Tomaso (FLI Jena) |
| 12 | Brucella canis (BM385) | 618.3226 | 2.7912 | dbspec-03-Mar-2023-12-16-11.379 | TNO Netherlands |
| 13 | Brucella neotomae A148-7 | 608.7357 | 2.7844 | dbspec-03-Mar-2023-12-24-20.374 | RKI ZBS2\|ZBS6 |
| 14 | Brucella abortus S19 delta mglA3.14 | 604.7419 | 2.7816 | dbspec-03-Mar-2023-12-14-50.428 | RKI FG14 |
| 15 | Brucella abortus Ring trial A-269 06 A | 600.2398 | 2.7783 | dbspec-03-Mar-2023-12-14-16.194 | RKI ZBS2\|ZBS6 |
| 16 | Brucella abortus 2 Strain 86 8 59 | 600.0188 | 2.7782 | dbspec-03-Mar-2023-12-12-15.385 | Dr. Tomaso (FLI Jena) |
| 17 | Brucella sp. A 1755_25 | 596.046 | 2.7753 | dbspec-03-Mar-2023-12-27-17.183 | RKI ZBS6 |
| 18 | Brucella melitensis 02 3 022 17 | 595.316 | 2.7747 | dbspec-03-Mar-2023-12-20-44.98 | Dr. Tomaso (FLI Jena) |
| 19 | Brucella abortus 1 EQADeBa 09RB8922 | 594.6852 | 2.7743 | dbspec-03-Mar-2023-12-11-53.229 | Dr. Tomaso (FLI Jena) |
| 20 | Brucella sp. A 1755_28 | 593.5357 | 2.7734 | dbspec-03-Mar-2023-12-27-39.807 | RKI ZBS6 |

#### # 7 - IDENTIFICATION REPORT FOR MALDI-TOF MS SPECTRUM *'2013_06_17_Maren_Stämmler_0001'*

##### Metadata of actual MALDI-ToF test spectrum

| | |
|---|---|
| **genus / species / strain:** | RKI MALDI sample 04, QUANDHIP EQAE ring trial, strains provided by RKI \| ZBS 2 |
| **file id:** | 2013_06_17_Maren_Stämmler_0001 |
| **type:** | Measurement 01 |
| **Bruker ID:** | 7DEDDBEA-CB5D-49B9-B905-F8942788B5EC |
| **NCBI ID (primary):** | 1392 |
| **NCBI ID (secondary):** | 1392 |
| **growth time:** | Optimal growth time between 24 - 72h |
| **growth temperature:** | 37°C |
| **growth conditions:** | Optimal aerobic or microaerophilic conditions |
| **growth medium:** | Columbia blood agar (Oxoid), 2nd passage on TSA or Caso agar, harvested by the 2nd passage |
| **sample treatment:** | Sample mixed with 20 mkL 100Â§ TFA, final TFA conc. approx. 80 perc.; approx. 30 Min treatment time; diluted 1:10 (vol); mixed with 1:1 HCCA\|TA2(A) |
| **spores:** | No |
| **concentration:** | Pellet produced by centrifugation (1 x 5 Min, 15,000 rpm) of a 500 mkL cell suspension, gamma ray irradiated (30 kGy) |
| **extra info:** | Bacillus anthracis AMES: microbial preparation by ZBS 2\| preparation for MALDI-ToF MS: M. Stämmler |
| **calibration standard:** | linear calibration using Escherichia coli DSM 3871 |
| **measurement method:** | D:\Methods\flexControlMethods\MaierMethods\ToM_200ns_20130611.par |
| **customer:** | RKI ZBS 2\| ZBS 6 |
| **measurement date/time:** | 2013-06-17T09:21:01.468+02:00 |
| **path to MS file:** | C:\Users\LaschP\Documents\MATLAB\Microbe MS testdata\ring trial RKI spectra\Sample_04\0_E19\1\1SLin |

##### Identification results: analysis of score ranking list

| No. | Genus/Species | Score | Log Score | UniProt Identifier | No. of strains in DB |
|---|---|---|---|---|---|
| 1 | Bacillus anthracis | 870.3009 | 2.9397 | 1392 | 130 |
| 2 | Bacillus cereus | 0.0012764 | -2.894 | 1392.1 | 6 |
| 3 | n/a | 0 | -Inf | | |

##### Score ranking list: best matches with test spectrum

| No. | Genus/Species/Strain | Score | Log Score | Spectrum identifier | Customer, or UniprotKB link |
|---|---|---|---|---|---|
| 1 | Bacillus anthracis unknown origin (A18, Beyer) | 756.1898 | 2.8786 | dbspec-03-Mar-2023-11-11-19.059 | RKI ZBS6, Dr. Beyer (Uni Hohenheim) |
| 2 | Bacillus anthracis unknown origin (A61, Beyer) | 755.8394 | 2.8784 | dbspec-03-Mar-2023-11-13-26.283 | RKI ZBS6, Dr. Beyer (Uni Hohenheim) |
| 3 | Bacillus anthracis unknown origin (A27, Beyer) | 726.6941 | 2.8614 | dbspec-03-Mar-2023-11-11-45.794 | RKI ZBS6, Dr. Beyer (Uni Hohenheim) |
| 4 | Bacillus anthracis unknown origin (A7, Beyer) | 724.9825 | 2.8603 | dbspec-03-Mar-2023-11-13-54.559 | RKI ZBS6, Dr. Beyer (Uni Hohenheim) |
| 5 | Bacillus anthracis unknown origin (A10, Beyer) | 698.519 | 2.8442 | dbspec-03-Mar-2023-11-9-18.763 | RKI ZBS6, Dr. Beyer (Uni Hohenheim) |
| 6 | Bacillus anthracis unknown origin (A5, Beyer) | 698.011 | 2.8439 | dbspec-03-Mar-2023-11-13-7.803 | RKI ZBS6, Dr. Beyer (Uni Hohenheim) |
| 7 | Bacillus anthracis unknown origin (A9, Beyer) | 696.8295 | 2.8431 | dbspec-03-Mar-2023-11-14-41.907 | RKI ZBS6, Dr. Beyer (Uni Hohenheim) |
| 8 | Bacillus anthracis unknown origin (A35, Beyer) | 695.7209 | 2.8424 | dbspec-03-Mar-2023-11-12-18.584 | RKI ZBS6, Dr. Beyer (Uni Hohenheim) |
| 9 | Bacillus anthracis ATCC 4229 (RKI) | 694.8551 | 2.8419 | dbspec-03-Mar-2023-11-6-56.498 | RKI ZBS2 |
| 10 | Bacillus anthracis unknown origin (A63, Beyer) | 694.7324 | 2.8418 | dbspec-03-Mar-2023-11-13-35.513 | RKI ZBS6, Dr. Beyer (Uni Hohenheim) |
| 11 | Bacillus anthracis unknown origin (A121, Beyer) | 694.3252 | 2.8416 | dbspec-03-Mar-2023-11-10-40.859 | RKI ZBS6, Dr. Beyer (Uni Hohenheim) |
| 12 | Bacillus anthracis unknown origin, black (A89, Beyer) | 692.9598 | 2.8407 | dbspec-03-Mar-2023-11-14-55.859 | RKI ZBS6, Dr. Beyer (Uni Hohenheim) |
| 13 | Bacillus anthracis Sterne vaccine (A118, Beyer) | 691.3421 | 2.8397 | dbspec-03-Mar-2023-11-7-39.84 | RKI ZBS6, Dr. Beyer (Uni Hohenheim) |
| 14 | Bacillus anthracis unknown origin (A62, Beyer) | 690.1126 | 2.8389 | dbspec-03-Mar-2023-11-13-30.361 | RKI ZBS6, Dr. Beyer (Uni Hohenheim) |
| 15 | Bacillus anthracis unknown origin (A73, Beyer) | 664.9096 | 2.8228 | dbspec-03-Mar-2023-11-13-59.372 | RKI ZBS6, Dr. Beyer (Uni Hohenheim) |
| 16 | Bacillus cereus DSM 8438 (B248, Beyer) | 664.757 | 2.8227 | dbspec-03-Mar-2023-11-21-17.457 | RKI ZBS6, Dr. Beyer (Uni Hohenheim) |
| 17 | Bacillus anthracis unknown origin (A3, Beyer) | 662.7567 | 2.8214 | dbspec-03-Mar-2023-11-11-59.272 | RKI ZBS6, Dr. Beyer (Uni Hohenheim) |
| 18 | Bacillus anthracis ATCC 14185 (A2, Beyer) | 662.4046 | 2.8211 | dbspec-03-Mar-2023-11-6-45.935 | RKI ZBS6, Dr. Beyer (Uni Hohenheim) |
| 19 | Bacillus anthracis unknown origin (A106, Beyer) | 661.4475 | 2.8205 | dbspec-03-Mar-2023-11-9-41.083 | RKI ZBS6, Dr. Beyer (Uni Hohenheim) |
| 20 | Bacillus anthracis unknown origin (A84, Beyer) | 661.3295 | 2.8204 | dbspec-03-Mar-2023-11-14-27.907 | RKI ZBS6, Dr. Beyer (Uni Hohenheim) |

#### # 8 - IDENTIFICATION REPORT FOR MALDI-TOF MS SPECTRUM *'2013_06_14_Maren_Stämmler_0004'*

#### Metadata of actual MALDI-ToF test spectrum

| | |
|---|---|
| **genus / species / strain:** | RKI MALDI sample 04, QUANDHIP EQAE ring trial, strains provided by RKI \| ZBS 2 |
| **file id:** | 2013_06_14_Maren_Stämmler_0004 |
| **type:** | Measurement 02 |
| **Bruker ID:** | 0AFF63F5-F910-4924-A353-4FAA9835969D |
| **NCBI ID (primary):** | 1392 |
| **NCBI ID (secondary):** | 1392 |
| **growth time:** | Optimal growth time between 24 - 72h |
| **growth temperature:** | 37°C |
| **growth conditions:** | Optimal aerobic or microaerophilic conditions |
| **growth medium:** | Columbia blood agar (Oxoid), 2nd passage on TSA or Caso agar, harvested by the 2nd passage |
| **sample treatment:** | Sample mixed with 25 mkL 100Â§ TFA, final TFA conc. approx. 80 perc.; 10 mkL 80 perc. TFA added; approx. 30 Min treatment time; diluted 1:10 (vol); mixed with 1:1 HCCA\|TA2(A) |
| **spores:** | No |
| **concentration:** | Pellet produced by centrifugation (1 x 5 Min, 15,000 rpm) of a 500 mkL cell suspension, gamma ray irradiated (30 kGy) |
| **extra info:** | Bacillus anthracis AMES: microbial preparation by ZBS 2\| preparation for MALDI-ToF MS: M. Stämmler |
| **calibration standard:** | linear calibration using Escherichia coli DSM 3871 |
| **measurement method:** | D:\Methods\flexControlMethods\MaierMethods\ToM_200ns_20130611.par |
| **customer:** | RKI ZBS 2\| ZBS 6 |
| **measurement date/time:** | 2013-06-14T08:52:26.953+02:00 |
| **path to MS file:** | C:\Users\LaschP\Documents\MATLAB\Microbe MS testdata\ring trial RKI spectra\Sample_04\0_H10\1\1SLin |

#### Identification results: analysis of score ranking list

| No. | Genus/Species | Score | Log Score | UniProt Identifier | No. of strains in DB |
|---|---|---|---|---|---|
| 1 | Bacillus anthracis | 867.6172 | 2.9383 | 1392 | 130 |
| 2 | Bacillus cereus | 14.0644 | 1.1481 | 1392.1 | 6 |
| 3 | n/a | 0 | -Inf | | |

#### Score ranking list: best matches with test spectrum

| No. | Genus/Species/Strain | Score | Log Score | Spectrum identifier | Customer, or UniprotKB link |
|---|---|---|---|---|---|
| 1 | Bacillus anthracis unknown origin (A10, Beyer) | 786.5829 | 2.8957 | dbspec-03-Mar-2023-11-9-18.763 | RKI ZBS6, Dr. Beyer (Uni Hohenheim) |
| 2 | Bacillus anthracis unknown origin (A7, Beyer) | 750.2069 | 2.8752 | dbspec-03-Mar-2023-11-13-54.559 | RKI ZBS6, Dr. Beyer (Uni Hohenheim) |
| 3 | Bacillus anthracis unknown origin (A61, Beyer) | 749.3855 | 2.8747 | dbspec-03-Mar-2023-11-13-26.283 | RKI ZBS6, Dr. Beyer (Uni Hohenheim) |
| 4 | Bacillus anthracis Sterne vaccine (A1, Beyer) | 748.625 | 2.8743 | dbspec-03-Mar-2023-11-7-35.242 | RKI ZBS6, Dr. Beyer (Uni Hohenheim) |
| 5 | Bacillus anthracis unknown origin (A5, Beyer) | 727.7025 | 2.862 | dbspec-03-Mar-2023-11-13-7.803 | RKI ZBS6, Dr. Beyer (Uni Hohenheim) |
| 6 | Bacillus anthracis unknown origin (A18, Beyer) | 720.716 | 2.8578 | dbspec-03-Mar-2023-11-11-19.059 | RKI ZBS6, Dr. Beyer (Uni Hohenheim) |
| 7 | Bacillus cereus DSM 8438 (B248, Beyer) | 719.2673 | 2.8569 | dbspec-03-Mar-2023-11-21-17.457 | RKI ZBS6, Dr. Beyer (Uni Hohenheim) |
| 8 | Bacillus anthracis ATCC 4229 (RKI) | 717.3501 | 2.8557 | dbspec-03-Mar-2023-11-6-56.498 | RKI ZBS2 |
| 9 | Bacillus anthracis unknown origin (A28, Beyer) | 699.3673 | 2.8447 | dbspec-03-Mar-2023-11-11-49.856 | RKI ZBS6, Dr. Beyer (Uni Hohenheim) |
| 10 | Bacillus anthracis unknown origin (A36, Beyer) | 697.5034 | 2.8435 | dbspec-03-Mar-2023-11-12-23.443 | RKI ZBS6, Dr. Beyer (Uni Hohenheim) |
| 11 | Bacillus anthracis unknown origin (A9, Beyer) | 694.6064 | 2.8417 | dbspec-03-Mar-2023-11-14-41.907 | RKI ZBS6, Dr. Beyer (Uni Hohenheim) |
| 12 | Bacillus anthracis unknown origin (A73, Beyer) | 693.8598 | 2.8413 | dbspec-03-Mar-2023-11-13-59.372 | RKI ZBS6, Dr. Beyer (Uni Hohenheim) |
| 13 | Bacillus anthracis unknown origin (A27, Beyer) | 691.4017 | 2.8397 | dbspec-03-Mar-2023-11-11-45.794 | RKI ZBS6, Dr. Beyer (Uni Hohenheim) |
| 14 | Bacillus anthracis unknown origin (A63, Beyer) | 691.2405 | 2.8396 | dbspec-03-Mar-2023-11-13-35.513 | RKI ZBS6, Dr. Beyer (Uni Hohenheim) |
| 15 | Bacillus anthracis unknown origin (A121, Beyer) | 689.2194 | 2.8384 | dbspec-03-Mar-2023-11-10-40.859 | RKI ZBS6, Dr. Beyer (Uni Hohenheim) |
| 16 | Bacillus anthracis unknown origin (A6, Beyer) | 688.8601 | 2.8381 | dbspec-03-Mar-2023-11-13-17.158 | RKI ZBS6, Dr. Beyer (Uni Hohenheim) |
| 17 | Bacillus anthracis Sterne vaccine (A118, Beyer) | 687.2482 | 2.8371 | dbspec-03-Mar-2023-11-7-39.84 | RKI ZBS6, Dr. Beyer (Uni Hohenheim) |
| 18 | Bacillus anthracis unknown origin (A35, Beyer) | 666.7254 | 2.8239 | dbspec-03-Mar-2023-11-12-18.584 | RKI ZBS6, Dr. Beyer (Uni Hohenheim) |
| 19 | Bacillus anthracis unknown origin (A3, Beyer) | 666.5561 | 2.8238 | dbspec-03-Mar-2023-11-11-59.272 | RKI ZBS6, Dr. Beyer (Uni Hohenheim) |
| 20 | Bacillus anthracis unknown origin (A16, Beyer) | 666.2856 | 2.8237 | dbspec-03-Mar-2023-11-11-14.107 | RKI ZBS6, Dr. Beyer (Uni Hohenheim) |

---

### # 9 - IDENTIFICATION REPORT FOR MALDI-TOF MS SPECTRUM *'2013_06_13_Maren_Stämmler_0004'*

#### Metadata of actual MALDI-ToF test spectrum

| | |
|---|---|
| **genus / species / strain:** | RKI MALDI sample 04, QUANDHIP EQAE ring trial, strains provided by RKI \| ZBS 2 |
| **file id:** | 2013_06_13_Maren_Stämmler_0004 |
| **type:** | Measurement 03 |
| **Bruker ID:** | 27965397-8164-4463-BB40-0FD314995834 |
| **NCBI ID (primary):** | 1392 |
| **NCBI ID (secondary):** | 1392 |
| **growth time:** | Optimal growth time between 24 - 72h |
| **growth temperature:** | 37°C |
| **growth conditions:** | Optimal aerobic or microaerophilic conditions |
| **growth medium:** | Columbia blood agar (Oxoid), 2nd passage on TSA or Caso agar, harvested by the 2nd passage |
| **sample treatment:** | Sample mixed with 25 mkL 100Â§ TFA, final TFA conc. approx. 80 perc.; 10 mkL 80 perc. TFA added; approx. 30 Min treatment time; diluted 1:10 (vol); mixed with 1:1 HCCA\|TA2(A) |
| **spores:** | No |
| **concentration:** | Pellet produced by centrifugation (1 x 5 Min, 15,000 rpm) of a 500 mkL cell suspension, gamma ray irradiated (30 kGy) |
| **extra info:** | Bacillus anthracis AMES: microbial preparation by ZBS 2\| preparation for MALDI-ToF MS: M. Stämmler |
| **calibration standard:** | linear calibration using Escherichia coli DSM 3871 |

| | |
|---|---|
| **measurement method:** | D:\Methods\flexControlMethods\MaierMethods\ToM_200ns_20130611.par |
| **customer:** | RKI ZBS 2\| ZBS 6 |
| **measurement date/time:** | 2013-06-13T14:19:29.671+02:00 |
| **path to MS file:** | C:\Users\LaschP\Documents\MATLAB\Microbe MS testdata\ring trial RKI spectra\Sample_04\0_H9\1\1SLin |

##### Identification results: analysis of score ranking list

| No. | Genus/Species | Score | Log Score | UniProt Identifier | No. of strains in DB |
|---|---|---|---|---|---|
| 1 | Bacillus anthracis | 847.6911 | 2.9282 | 1392 | 130 |
| 2 | Bacillus cereus | 3.3578 | 0.52605 | 1392.1 | 6 |
| 3 | Bacillus cereus s.l. | 0.001351 | -2.8693 | 86661 | 22 |
| 4 | Bacillus cereus | 1.1832e-05 | -4.9269 | 1396 | 102 |

##### Score ranking list: best matches with test spectrum

| No. | Genus/Species/Strain | Score | Log Score | Spectrum identifier | Customer, or UniprotKB link |
|---|---|---|---|---|---|
| 1 | Bacillus anthracis ATCC 4229 (RKI) | 754.6104 | 2.8777 | dbspec-03-Mar-2023-11-6-56.498 | RKI ZBS2 |
| 2 | Bacillus anthracis unknown origin (A27, Beyer) | 692.4448 | 2.8404 | dbspec-03-Mar-2023-11-11-45.794 | RKI ZBS6, Dr. Beyer (Uni Hohenheim) |
| 3 | Bacillus anthracis unknown origin (A61, Beyer) | 691.2708 | 2.8396 | dbspec-03-Mar-2023-11-13-26.283 | RKI ZBS6, Dr. Beyer (Uni Hohenheim) |
| 4 | Bacillus anthracis unknown origin (A10, Beyer) | 662.9844 | 2.8215 | dbspec-03-Mar-2023-11-9-18.763 | RKI ZBS6, Dr. Beyer (Uni Hohenheim) |
| 5 | Bacillus anthracis unknown origin (A9, Beyer) | 661.4401 | 2.8205 | dbspec-03-Mar-2023-11-14-41.907 | RKI ZBS6, Dr. Beyer (Uni Hohenheim) |
| 6 | Bacillus anthracis unknown origin (A7, Beyer) | 660.7224 | 2.82 | dbspec-03-Mar-2023-11-13-54.559 | RKI ZBS6, Dr. Beyer (Uni Hohenheim) |
| 7 | Bacillus anthracis unknown origin (A121, Beyer) | 660.2319 | 2.8197 | dbspec-03-Mar-2023-11-10-40.859 | RKI ZBS6, Dr. Beyer (Uni Hohenheim) |
| 8 | Bacillus anthracis unknown origin (A63, Beyer) | 659.5261 | 2.8192 | dbspec-03-Mar-2023-11-13-35.513 | RKI ZBS6, Dr. Beyer (Uni Hohenheim) |
| 9 | Bacillus cereus DSM 8438 (B248, Beyer) | 659.4425 | 2.8192 | dbspec-03-Mar-2023-11-21-17.457 | RKI ZBS6, Dr. Beyer (Uni Hohenheim) |
| 10 | Bacillus anthracis unknown origin (A18, Beyer) | 657.7954 | 2.8181 | dbspec-03-Mar-2023-11-11-19.059 | RKI ZBS6, Dr. Beyer (Uni Hohenheim) |
| 11 | Bacillus anthracis Stamatin vaccine (A15, Beyer) | 653.396 | 2.8152 | dbspec-03-Mar-2023-11-7-26.93 | RKI ZBS6, Dr. Beyer (Uni Hohenheim) |
| 12 | Bacillus anthracis unknown origin (A28, Beyer) | 634.6703 | 2.8025 | dbspec-03-Mar-2023-11-11-49.856 | RKI ZBS6, Dr. Beyer (Uni Hohenheim) |
| 13 | Bacillus anthracis unknown origin (A73, Beyer) | 630.5422 | 2.7997 | dbspec-03-Mar-2023-11-13-59.372 | RKI ZBS6, Dr. Beyer (Uni Hohenheim) |
| 14 | Bacillus anthracis unknown origin (A62, Beyer) | 628.292 | 2.7982 | dbspec-03-Mar-2023-11-13-30.361 | RKI ZBS6, Dr. Beyer (Uni Hohenheim) |
| 15 | Bacillus anthracis unknown origin (A107, Beyer) | 624.2742 | 2.7954 | dbspec-03-Mar-2023-11-9-45.489 | RKI ZBS6, Dr. Beyer (Uni Hohenheim) |
| 16 | Bacillus cereus s.l. CD 3-1c | 620.4967 | 2.7927 | dbspec-03-Mar-2023-11-26-3.052 | RKI ZBS6 \| Dr. Thanh Tam (IPPR, Hanoi, Vietnam) |
| 17 | Bacillus cereus s.l. CD 3-1a | 615.397 | 2.7892 | dbspec-03-Mar-2023-11-25-52.443 | RKI ZBS6 \| Dr. Thanh Tam (IPPR, Hanoi, Vietnam) |
| 18 | Bacillus cereus DSM 2302 (B192, Beyer) | 603.9321 | 2.781 | dbspec-03-Mar-2023-11-18-47.867 | RKI ZBS6, Dr. Beyer (Uni Hohenheim) |
| 19 | Bacillus anthracis unknown origin (A5, Beyer) | 602.0537 | 2.7796 | dbspec-03-Mar-2023-11-13-7.803 | RKI ZBS6, Dr. Beyer (Uni Hohenheim) |
| 20 | Bacillus anthracis unknown origin (A35, Beyer) | 599.1749 | 2.7776 | dbspec-03-Mar-2023-11-12-18.584 | RKI ZBS6, Dr. Beyer (Uni Hohenheim) |

---

#### # 10 - IDENTIFICATION REPORT FOR MALDI-TOF MS SPECTRUM *'2013_06_13_Maren_Stämmler_0005'*

##### Metadata of actual MALDI-ToF test spectrum

| | |
|---|---|
| **genus / species / strain:** | RKI MALDI sample 05, QUANDHIP EQAE ring trial, strains provided by RKI \| ZBS 2 |
| **file id:** | 2013_06_13_Maren_Stämmler_0005 |
| **type:** | Measurement 01 |
| **Bruker ID:** | DB87AC50-0ED5-4ACA-B2F9-4B34DE757E1B |
| **NCBI ID (primary):** | 529 |
| **NCBI ID (secondary):** | 529 |
| **growth time:** | Optimal growth time between 24 - 72h |
| **growth temperature:** | 37°C |
| **growth conditions:** | Optimal aerobic or microaerophilic conditions |
| **growth medium:** | Columbia blood agar (Oxoid), 2nd passage on TSA or Caso agar, harvested by the 2nd passage |
| **sample treatment:** | Sample mixed with 20 mkL 100Â§ TFA, final TFA conc. approx. 80 perc.; approx. 30 Min treatment time; diluted 1:10 (vol); mixed with 1:1 HCCA\|TA2(A) |
| **spores:** | No |
| **concentration:** | Pellet produced by centrifugation (1 x 5 Min, 15,000 rpm) of a 500 mkL cell suspension, gamma ray irradiated (30 kGy) |
| **extra info:** | Ochrobactrum anthropi A-148-11: microbial preparation by ZBS 2\| preparation for MALDI-ToF MS: M. Stämmler |
| **calibration standard:** | linear calibration using Escherichia coli DSM 3871 |
| **measurement method:** | D:\Methods\flexControlMethods\MaierMethods\ToM_200ns_20130611.par |
| **customer:** | RKI ZBS 2\| ZBS 6 |
| **measurement date/time:** | 2013-06-13T14:25:30.359+02:00 |
| **path to MS file:** | C:\Users\LaschP\Documents\MATLAB\Microbe MS testdata\ring trial RKI spectra\Sample_05\0_H11\1\1SLin |

##### Identification results: analysis of score ranking list

| No. | Genus/Species | Score | Log Score | UniProt Identifier | No. of strains in DB |
|---|---|---|---|---|---|
| 1 | Ochrobactrum anthropi | 828.7715 | 2.9184 | 529 | 2 |
| 2 | Brucella intermedia | 128.5477 | 2.1091 | 94625 | 1 |
| 3 | Burkholderia plantarii | 50.2247 | 1.7009 | 41899 | 2 |
| 4 | Ralstonia mannitolilytica | 27.9507 | 1.4464 | 105219 | 1 |

| | | | | | |
|---|---|---|---|---|---|
| 5 | Brucella vulpis | 14.1418 | 1.1505 | 981386 | 1 |

##### Score ranking list: best matches with test spectrum

| No. | Genus/Species/Strain | Score | Log Score | Spectrum identifier | Customer, or UniprotKB link |
|---|---|---|---|---|---|
| 1 | Ochrobactrum anthropi DSM 20150, ATCC 11425 | 697.8836 | 2.8438 | dbspec-03-Mar-2023-13-26-27.357 | RKI ZBS6 |
| 2 | Ochrobactrum anthropi Ring trial A-269 08 A | 638.1946 | 2.805 | dbspec-03-Mar-2023-13-26-25.107 | RKI ZBS2\|ZBS6 |
| 3 | Brucella intermedia A1309 | 347.4749 | 2.5409 | dbspec-03-Mar-2023-12-19-39.95 | RKI ZBS6 |
| 4 | Burkholderia plantarii LMG 9035 | 225.5588 | 2.3533 | dbspec-03-Mar-2023-12-50-19.622 | RKI ZBS2\|ZBS6 |
| 5 | Ralstonia mannitolilytica DSM 17512, LMG 6866 | 218.5235 | 2.3395 | dbspec-03-Mar-2023-13-34-9.843 | RKI ZBS6 |
| 6 | Brucella vulpis DSM 101715 | 202.7911 | 2.307 | dbspec-03-Mar-2023-12-30-19.241 | RKI ZBS6 |
| 7 | Brucella abortus S19 | 195.6302 | 2.2914 | dbspec-03-Mar-2023-12-14-44.397 | RKI FG14 |
| 8 | Brucella sp. 83-13 | 179.2401 | 2.2534 | dbspec-03-Mar-2023-12-27-0.761 | RKI FG14 |
| 9 | Brucella suis biovar 1 (BM431) | 176.8586 | 2.2476 | dbspec-03-Mar-2023-12-28-37.337 | TNO Netherlands |
| 10 | Brucella canis (BM385) | 176.0257 | 2.2456 | dbspec-03-Mar-2023-12-16-11.379 | TNO Netherlands |
| 11 | Streptococcus parasanguinis B 9056 | 168.7668 | 2.2273 | dbspec-03-Mar-2023-13-45-42.182 | RKI ZBS6 |
| 12 | Brucella abortus S19 delta mglA3.14 | 166.5756 | 2.2216 | dbspec-03-Mar-2023-12-14-50.428 | RKI FG14 |
| 13 | Burkholderia plantarii LMG 10907 | 166.228 | 2.2207 | dbspec-03-Mar-2023-12-50-16.357 | RKI ZBS2\|ZBS6 |
| 14 | Brucella abortus 1 EQADeBa 09RB8922 | 166.0785 | 2.2203 | dbspec-03-Mar-2023-12-11-53.229 | Dr. Tomaso (FLI Jena) |
| 15 | Inquilinus limosus DSM 16000, LMG 20952, CCUG 45653 | 163.6593 | 2.2139 | dbspec-03-Mar-2023-13-20-12.005 | RKI ZBS6 |
| 16 | Brucella abortus NCTC 10503 07RB1200, ATCC 23451 | 160.4895 | 2.2054 | dbspec-03-Mar-2023-12-14-24.554 | Dr. Tomaso (FLI Jena) |
| 17 | Vibrio vulnificus A171-4 | 158.0477 | 2.1988 | dbspec-03-Mar-2023-13-52-49.641 | RKI ZBS2\|ZBS6 |
| 18 | Bacillus paralicheniformis EZ15-08 | 157.9801 | 2.1986 | dbspec-03-Mar-2023-11-47-29.432 | RKI ZBS6 \|Prof. Gao (Nanjing, China) |
| 19 | Staphylococcus delphini Referenzstamm | 154.9283 | 2.1901 | dbspec-03-Mar-2023-13-41-49.031 | RKI ZBS6 |
| 20 | Stenotrophomonas maltophilia Sm 132 | 154.3386 | 2.1885 | dbspec-03-Mar-2023-13-44-46.84 | RKI ZBS6 |

---

#### # 11 - IDENTIFICATION REPORT FOR MALDI-TOF MS SPECTRUM *'2013_06_14_Maren_Stämmler_0005'*

##### Metadata of actual MALDI-ToF test spectrum

| | |
|---|---|
| **genus / species / strain:** | RKI MALDI sample 05, QUANDHIP EQAE ring trial, strains provided by RKI \| ZBS 2 |
| **file id:** | 2013_06_14_Maren_Stämmler_0005 |
| **type:** | Measurement 02 |
| **Bruker ID:** | B8441899-48FF-4802-B5F8-4491FED91EF2 |
| **NCBI ID (primary):** | 529 |
| **NCBI ID (secondary):** | 529 |
| **growth time:** | Optimal growth time between 24 - 72h |
| **growth temperature:** | 37°C |
| **growth conditions:** | Optimal aerobic or microaerophilic conditions |
| **growth medium:** | Columbia blood agar (Oxoid), 2nd passage on TSA or Caso agar, harvested by the 2nd passage |
| **sample treatment:** | Sample mixed with 20 mkL 100Â§ TFA, final TFA conc. approx. 80 perc.; approx. 30 Min treatment time; diluted 1:10 (vol); mixed with 1:1 HCCA\|TA2(A) |
| **spores:** | No |
| **concentration:** | Pellet produced by centrifugation (1 x 5 Min, 15,000 rpm) of a 500 mkL cell suspension, gamma ray irradiated (30 kGy) |
| **extra info:** | Ochrobactrum anthropi A-148-11: microbial preparation by ZBS 2\| preparation for MALDI-ToF MS: M. Stämmler |
| **calibration standard:** | linear calibration using Escherichia coli DSM 3871 |
| **measurement method:** | D:\Methods\flexControlMethods\MaierMethods\ToM_200ns_20130611.par |
| **customer:** | RKI ZBS 2\| ZBS 6 |
| **measurement date/time:** | 2013-06-14T09:00:58.531+02:00 |
| **path to MS file:** | C:\Users\LaschP\Documents\MATLAB\Microbe MS testdata\ring trial RKI spectra\Sample_05\0_H12\1\1SLin |

##### Identification results: analysis of score ranking list

| No. | Genus/Species | Score | Log Score | UniProt Identifier | No. of strains in DB |
|---|---|---|---|---|---|
| 1 | Ochrobactrum anthropi | 827.607 | 2.9178 | 529 | 2 |
| 2 | Brucella intermedia | 128.348 | 2.1084 | 94625 | 1 |
| 3 | Burkholderia plantarii | 49.6714 | 1.6961 | 41899 | 2 |
| 4 | Brucella sp. | 16.8519 | 1.2266 | 52132 | 9 |
| 5 | Brucella vulpis | 14.1219 | 1.1499 | 981386 | 1 |

##### Score ranking list: best matches with test spectrum

| No. | Genus/Species/Strain | Score | Log Score | Spectrum identifier | Customer, or UniprotKB link |
|---|---|---|---|---|---|
| 1 | Ochrobactrum anthropi DSM 20150, ATCC 11425 | 696.7335 | 2.8431 | dbspec-03-Mar-2023-13-26-27.357 | RKI ZBS6 |
| 2 | Ochrobactrum anthropi Ring trial A-269 08 A | 635.4443 | 2.8031 | dbspec-03-Mar-2023-13-26-25.107 | RKI ZBS2\|ZBS6 |
| 3 | Brucella intermedia A1309 | 346.8032 | 2.5401 | dbspec-03-Mar-2023-12-19-39.95 | RKI ZBS6 |
| 4 | Burkholderia plantarii LMG 9035 | 223.5468 | 2.3494 | dbspec-03-Mar-2023-12-50-19.622 | RKI ZBS2\|ZBS6 |
| 5 | Brucella sp. 83-13 | 207.7896 | 2.3176 | dbspec-03-Mar-2023-12-27-0.761 | RKI FG14 |
| 6 | Brucella vulpis DSM 101715 | 202.5897 | 2.3066 | dbspec-03-Mar-2023-12-30-19.241 | RKI ZBS6 |
| 7 | Brucella abortus S19 | 195.6019 | 2.2914 | dbspec-03-Mar-2023-12-14-44.397 | RKI FG14 |

| | | | | | |
|---|---|---|---|---|---|
| 8 | Brucella suis biovar 1 (BM431) | 175.892 | 2.2452 | dbspec-03-Mar-2023-12-28-37.337 | TNO Netherlands |
| 9 | Brucella canis (BM385) | 175.6695 | 2.2447 | dbspec-03-Mar-2023-12-16-11.379 | TNO Netherlands |
| 10 | Streptococcus parasanguinis B 9056 | 167.4394 | 2.2239 | dbspec-03-Mar-2023-13-45-42.182 | RKI ZBS6 |
| 11 | Brucella abortus S19 delta mglA3.14 | 166.3675 | 2.2211 | dbspec-03-Mar-2023-12-14-50.428 | RKI FG14 |
| 12 | Brucella abortus 1 EQADeBa 09RB8922 | 165.9038 | 2.2199 | dbspec-03-Mar-2023-12-11-53.229 | Dr. Tomaso (FLI Jena) |
| 13 | Burkholderia plantarii LMG 10907 | 165.1248 | 2.2178 | dbspec-03-Mar-2023-12-50-16.357 | RKI ZBS2\|ZBS6 |
| 14 | Inquilinus limosus DSM 16000, LMG 20952, CCUG 45653 | 161.7731 | 2.2089 | dbspec-03-Mar-2023-13-20-12.005 | RKI ZBS6 |
| 15 | Brucella abortus NCTC 10503 07RB1200, ATCC 23451 | 161.5372 | 2.2083 | dbspec-03-Mar-2023-12-14-24.554 | Dr. Tomaso (FLI Jena) |
| 16 | Ralstonia mannitolilytica DSM 17512, LMG 6866 | 158.4631 | 2.1999 | dbspec-03-Mar-2023-13-34-9.843 | RKI ZBS6 |
| 17 | Stenotrophomonas maltophilia Sm 132 | 154.9842 | 2.1903 | dbspec-03-Mar-2023-13-44-46.84 | RKI ZBS6 |
| 18 | Staphylococcus delphini Referenzstamm | 154.0659 | 2.1877 | dbspec-03-Mar-2023-13-41-49.031 | RKI ZBS6 |
| 19 | Staphylococcus pseudintermedius DSM 21284, CCUG 49543, LMG 22219, CIP 108864 | 152.3643 | 2.1829 | dbspec-03-Mar-2023-13-44-32.168 | RKI ZBS6 |
| 20 | Stenotrophomonas maltophilia Sm 20\|06 | 151.9338 | 2.1817 | dbspec-03-Mar-2023-13-44-48.652 | RKI ZBS6 |

---

#### # 12 - IDENTIFICATION REPORT FOR MALDI-TOF MS SPECTRUM *'2013_06_13_Maren_Stämmler_0006'*

##### Metadata of actual MALDI-ToF test spectrum

| | |
|---|---|
| **genus / species / strain:** | RKI MALDI sample 06, QUANDHIP EQAE ring trial, strains provided by RKI \| ZBS 2 |
| **file id:** | 2013_06_13_Maren_Stämmler_0006 |
| **type:** | Measurement 01 |
| **Bruker ID:** | A91F9B76-08DA-40E6-9154-5F7CE8EF3789 |
| **NCBI ID (primary):** | 633 |
| **NCBI ID (secondary):** | 633 |
| **growth time:** | Optimal growth time between 24 - 72h |
| **growth temperature:** | 37°C |
| **growth conditions:** | Optimal aerobic or microaerophilic conditions |
| **growth medium:** | Columbia blood agar (Oxoid), 2nd passage on TSA or Caso agar, harvested by the 2nd passage |
| **sample treatment:** | Sample mixed with 20 mkL 100Â§ TFA, final TFA conc. approx. 80 perc.; approx. 30 Min treatment time; diluted 1:10 (vol); mixed with 1:1 HCCA\|TA2(A) |
| **spores:** | No |
| **concentration:** | Pellet produced by centrifugation (1 x 5 Min, 15,000 rpm) of a 500 mkL cell suspension, gamma ray irradiated (30 kGy) |
| **extra info:** | Yersinia pseudotuberculosis III: microbial preparation by ZBS 2\| preparation for MALDI-ToF MS: M. Stämmler |
| **calibration standard:** | linear calibration using Escherichia coli DSM 3871 |
| **measurement method:** | D:\Methods\flexControlMethods\MaierMethods\ToM_200ns_20130611.par |
| **customer:** | RKI ZBS 2\| ZBS 6 |
| **measurement date/time:** | 2013-06-13T14:36:26.328+02:00 |
| **path to MS file:** | C:\Users\LaschP\Documents\MATLAB\Microbe MS testdata\ring trial RKI spectra\Sample_06\0_H13\1\1SLin |

##### Identification results: analysis of score ranking list

| No. | Genus/Species | Score | Log Score | UniProt Identifier | No. of strains in DB |
|---|---|---|---|---|---|
| 1 | Yersinia pseudotuberculosis | 748.7865 | 2.8744 | 633 | 24 |
| 2 | Yersinia similis | 67.6672 | 1.8304 | 367190 | 2 |
| 3 | Yersinia pestis | 44.6106 | 1.6494 | 632 | 11 |

##### Score ranking list: best matches with test spectrum

| No. | Genus/Species/Strain | Score | Log Score | Spectrum identifier | Customer, or UniprotKB link |
|---|---|---|---|---|---|
| 1 | Yersinia pseudotuberculosis 29827 | 698.9029 | 2.8444 | dbspec-03-Mar-2023-14-1-24.021 | RKI ZBS6 |
| 2 | Yersinia pseudotuberculosis VI | 694.5313 | 2.8417 | dbspec-03-Mar-2023-14-1-50.63 | RKI ZBS6 |
| 3 | Yersinia pseudotuberculosis 25743 | 674.0433 | 2.8287 | dbspec-03-Mar-2023-14-1-1.334 | RKI ZBS6 |
| 4 | Yersinia pseudotuberculosis 29490 | 673.706 | 2.8285 | dbspec-03-Mar-2023-14-1-20.802 | RKI ZBS6 |
| 5 | Yersinia similis DSM 18211, LMG 23763, CCUG 52882 | 663.5998 | 2.8219 | dbspec-03-Mar-2023-14-2-21.16 | CVUA Stuttgart |
| 6 | Yersinia pestis NCTC 2868 | 657.2594 | 2.8177 | dbspec-03-Mar-2023-13-59-28.93 | Dr. Drevinek (Sujchbo, Czech Republic) |
| 7 | Yersinia pestis NCTC 10030 | 648.935 | 2.8122 | dbspec-03-Mar-2023-13-58-55.571 | Dr. Drevinek (Sujchbo, Czech Republic) |
| 8 | Yersinia pestis NCTC 5923, ATCC 19428 | 646.7817 | 2.8108 | dbspec-03-Mar-2023-13-59-54.507 | Dr. Drevinek (Sujchbo, Czech Republic) |
| 9 | Yersinia pseudotuberculosis J9 | 642.9471 | 2.8082 | dbspec-03-Mar-2023-14-1-27.115 | RKI ZBS6 |
| 10 | Yersinia pseudotuberculosis 27707 | 640.6081 | 2.8066 | dbspec-03-Mar-2023-14-1-11.209 | RKI ZBS6 |
| 11 | Yersinia pestis NCTC 570 | 638.0201 | 2.8048 | dbspec-03-Mar-2023-13-59-40.07 | Dr. Drevinek (Sujchbo, Czech Republic) |
| 12 | Yersinia pestis NCTC 2028 | 633.0711 | 2.8015 | dbspec-03-Mar-2023-13-59-18.102 | Dr. Drevinek (Sujchbo, Czech Republic) |
| 13 | Yersinia pseudotuberculosis DSM 22972 | 626.721 | 2.7971 | dbspec-03-Mar-2023-14-1-34.286 | RKI ZBS6 |
| 14 | Yersinia pseudotuberculosis 04PA01423 | 614.4752 | 2.7885 | dbspec-03-Mar-2023-14-0-31.741 | RKI ZBS6 |
| 15 | Yersinia pseudotuberculosis 07PW12234 | 613.2194 | 2.7876 | dbspec-03-Mar-2023-14-0-54.053 | RKI ZBS6 |
| 16 | Yersinia pseudotuberculosis 27705 | 612.9517 | 2.7874 | dbspec-03-Mar-2023-14-1-7.881 | RKI ZBS6 |
| 17 | Yersinia pseudotuberculosis 25858 | 612.5346 | 2.7871 | dbspec-03-Mar-2023-14-1-4.677 | RKI ZBS6 |
| 18 | Yersinia similis IMB 4354 | 605.7913 | 2.7823 | dbspec-03-Mar-2023-14-2-37.113 | Prof. Fuchs (ZIEL, Munich) |
| 19 | Yersinia pseudotuberculosis 06PW40285 | 601.3232 | 2.7791 | dbspec-03-Mar-2023-14-0-47.272 | RKI ZBS6 |

| | | | | | |
|---|---|---|---|---|---|
| 20 | Yersinia pestis NCTC 10029 | 592.7404 | 2.7729 | dbspec-03-Mar-2023-13-58-45.946 | Dr. Drevinek (Sujchbo, Czech Republic) |

---

#### # 13 - IDENTIFICATION REPORT FOR MALDI-TOF MS SPECTRUM *'2013_06_14_Maren_Stämmler_0006'*

##### Metadata of actual MALDI-ToF test spectrum

| | |
|---|---|
| **genus / species / strain:** | RKI MALDI sample 06, QUANDHIP EQAE ring trial, strains provided by RKI \| ZBS 2 |
| **file id:** | 2013_06_14_Maren_Stämmler_0006 |
| **type:** | Measurement 02 |
| **Bruker ID:** | 0D79F381-905B-4F07-8C28-4E0E02288EDC |
| **NCBI ID (primary):** | 633 |
| **NCBI ID (secondary):** | 633 |
| **growth time:** | Optimal growth time between 24 - 72h |
| **growth temperature:** | 37°C |
| **growth conditions:** | Optimal aerobic or microaerophilic conditions |
| **growth medium:** | Columbia blood agar (Oxoid), 2nd passage on TSA or Caso agar, harvested by the 2nd passage |
| **sample treatment:** | Sample mixed with 20 mkL 100Â§ TFA, final TFA conc. approx. 80 perc.; approx. 30 Min treatment time; diluted 1:10 (vol); mixed with 1:1 HCCA\|TA2(A) |
| **spores:** | No |
| **concentration:** | Pellet produced by centrifugation (1 x 5 Min, 15,000 rpm) of a 500 mkL cell suspension, gamma ray irradiated (30 kGy) |
| **extra info:** | Yersinia pseudotuberculosis III: microbial preparation by ZBS 2\| preparation for MALDI-ToF MS: M. Stämmler |
| **calibration standard:** | linear calibration using Escherichia coli DSM 3871 |
| **measurement method:** | D:\Methods\flexControlMethods\MaierMethods\ToM_200ns_20130611.par |
| **customer:** | RKI ZBS 2\| ZBS 6 |
| **measurement date/time:** | 2013-06-14T09:22:37.468+02:00 |
| **path to MS file:** | C:\Users\LaschP\Documents\MATLAB\Microbe MS testdata\ring trial RKI spectra\Sample_06\0_H14\1\1SLin |

##### Identification results: analysis of score ranking list

| No. | Genus/Species | Score | Log Score | UniProt Identifier | No. of strains in DB |
|---|---|---|---|---|---|
| 1 | Yersinia pseudotuberculosis | 743.7506 | 2.8714 | 633 | 24 |
| 2 | Yersinia similis | 67.7773 | 1.8311 | 367190 | 2 |
| 3 | Yersinia pestis | 44.0243 | 1.6437 | 632 | 11 |

##### Score ranking list: best matches with test spectrum

| No. | Genus/Species/Strain | Score | Log Score | Spectrum identifier | Customer, or UniprotKB link |
|---|---|---|---|---|---|
| 1 | Yersinia pseudotuberculosis VI | 692.9056 | 2.8407 | dbspec-03-Mar-2023-14-1-50.63 | RKI ZBS6 |
| 2 | Yersinia pseudotuberculosis 25743 | 672.1942 | 2.8275 | dbspec-03-Mar-2023-14-1-1.334 | RKI ZBS6 |
| 3 | Yersinia pseudotuberculosis 29490 | 672.0967 | 2.8274 | dbspec-03-Mar-2023-14-1-20.802 | RKI ZBS6 |
| 4 | Yersinia pseudotuberculosis 29827 | 667.2003 | 2.8243 | dbspec-03-Mar-2023-14-1-24.021 | RKI ZBS6 |
| 5 | Yersinia similis DSM 18211, LMG 23763, CCUG 52882 | 665.6468 | 2.8232 | dbspec-03-Mar-2023-14-2-21.16 | CVUA Stuttgart |
| 6 | Yersinia pestis NCTC 2868 | 656.0374 | 2.8169 | dbspec-03-Mar-2023-13-59-28.93 | Dr. Drevinek (Sujchbo, Czech Republic) |
| 7 | Yersinia pestis NCTC 10030 | 647.3814 | 2.8112 | dbspec-03-Mar-2023-13-58-55.571 | Dr. Drevinek (Sujchbo, Czech Republic) |
| 8 | Yersinia pestis NCTC 5923, ATCC 19428 | 644.8415 | 2.8095 | dbspec-03-Mar-2023-13-59-54.507 | Dr. Drevinek (Sujchbo, Czech Republic) |
| 9 | Yersinia pseudotuberculosis 27705 | 644.5328 | 2.8092 | dbspec-03-Mar-2023-14-1-7.881 | RKI ZBS6 |
| 10 | Yersinia pseudotuberculosis J9 | 641.2641 | 2.807 | dbspec-03-Mar-2023-14-1-27.115 | RKI ZBS6 |
| 11 | Yersinia pseudotuberculosis 27707 | 639.0125 | 2.8055 | dbspec-03-Mar-2023-14-1-11.209 | RKI ZBS6 |
| 12 | Yersinia pestis NCTC 570 | 636.4798 | 2.8038 | dbspec-03-Mar-2023-13-59-40.07 | Dr. Drevinek (Sujchbo, Czech Republic) |
| 13 | Yersinia pestis NCTC 2028 | 631.244 | 2.8002 | dbspec-03-Mar-2023-13-59-18.102 | Dr. Drevinek (Sujchbo, Czech Republic) |
| 14 | Yersinia pseudotuberculosis 06PW40285 | 628.7718 | 2.7985 | dbspec-03-Mar-2023-14-0-47.272 | RKI ZBS6 |
| 15 | Yersinia pseudotuberculosis DSM 22972 | 627.9707 | 2.7979 | dbspec-03-Mar-2023-14-1-34.286 | RKI ZBS6 |
| 16 | Yersinia pestis NCTC 10029 | 621.3112 | 2.7933 | dbspec-03-Mar-2023-13-58-45.946 | Dr. Drevinek (Sujchbo, Czech Republic) |
| 17 | Yersinia pseudotuberculosis 04PA01423 | 612.8093 | 2.7873 | dbspec-03-Mar-2023-14-0-31.741 | RKI ZBS6 |
| 18 | Yersinia pseudotuberculosis 07PW12234 | 611.1162 | 2.7861 | dbspec-03-Mar-2023-14-0-54.053 | RKI ZBS6 |
| 19 | Yersinia pseudotuberculosis 25858 | 610.775 | 2.7859 | dbspec-03-Mar-2023-14-1-4.677 | RKI ZBS6 |
| 20 | Yersinia similis IMB 4354 | 606.712 | 2.783 | dbspec-03-Mar-2023-14-2-37.113 | Prof. Fuchs (ZIEL, Munich) |

---

#### # 14 - IDENTIFICATION REPORT FOR MALDI-TOF MS SPECTRUM *'2013_06_13_Maren_Stämmler_0007'*

##### Metadata of actual MALDI-ToF test spectrum

| | |
|---|---|
| **genus / species / strain:** | RKI MALDI sample 07, QUANDHIP EQAE ring trial, strains provided by RKI \| ZBS 2 |
| **file id:** | 2013_06_13_Maren_Stämmler_0007 |
| **type:** | Measurement 01 |

| | |
|---|---|
| **Bruker ID:** | 9E2A8556-22E3-4402-90AF-098AB16AC043 |
| **NCBI ID (primary):** | 13373 |
| **NCBI ID (secondary):** | 13373 |
| **growth time:** | Optimal growth time between 24 - 72h |
| **growth temperature:** | 37°C |
| **growth conditions:** | Optimal aerobic or microaerophilic conditions |
| **growth medium:** | Columbia blood agar (Oxoid), 2nd passage on TSA or Caso agar, harvested by the 2nd passage |
| **sample treatment:** | Sample mixed with 20 mkL 100Â§ TFA, final TFA conc. approx. 80 perc.; approx. 30 Min treatment time; diluted 1:10 (vol); mixed with 1:1 HCCA\|TA2(A) |
| **spores:** | No |
| **concentration:** | Pellet produced by centrifugation (1 x 5 Min, 15,000 rpm) of a 500 mkL cell suspension, gamma ray irradiated (30 kGy) |
| **extra info:** | Burkholderia mallei A106-3: microbial preparation by ZBS 2\| preparation for MALDI-ToF MS: M. Stämmler |
| **calibration standard:** | linear calibration using Escherichia coli DSM 3871 |
| **measurement method:** | D:\Methods\flexControlMethods\MaierMethods\ToM_200ns_20130611.par |
| **customer:** | RKI ZBS 2\| ZBS 6 |
| **measurement date/time:** | 2013-06-13T14:39:43.625+02:00 |
| **path to MS file:** | C:\Users\LaschP\Documents\MATLAB\Microbe MS testdata\ring trial RKI spectra\Sample_07\0_H15\1\1SLin |

#### Identification results: analysis of score ranking list

| No. | Genus/Species | Score | Log Score | UniProt Identifier | No. of strains in DB |
|---|---|---|---|---|---|
| 1 | Burkholderia mallei | 707.4652 | 2.8497 | 13373 | 33 |
| 2 | Burkholderia pseudomallei | 83.7394 | 1.9229 | 28450 | 20 |
| 3 | n/a | 0 | -Inf | | |

#### Score ranking list: best matches with test spectrum

| No. | Genus/Species/Strain | Score | Log Score | Spectrum identifier | Customer, or UniprotKB link |
|---|---|---|---|---|---|
| 1 | Burkholderia mallei Dubai 7 | 626.6822 | 2.797 | dbspec-03-Mar-2023-12-45-34.492 | Dr. Tomaso (FLI Jena) |
| 2 | Burkholderia mallei Dubai 7 240609RR5318 | 621.166 | 2.7932 | dbspec-03-Mar-2023-12-39-40.093 | Dr. Tomaso (FLI Jena) |
| 3 | Burkholderia mallei ATCC 23344 251109RR8925 | 620.2422 | 2.7926 | dbspec-03-Mar-2023-12-41-0.185 | Dr. Tomaso (FLI Jena) |
| 4 | Burkholderia pseudomallei PITT 5691 041206RR1064 | 606.988 | 2.7832 | dbspec-03-Mar-2023-12-52-51.791 | Dr. Tomaso (FLI Jena) |
| 5 | Burkholderia mallei Mukteswar 290103RR0041 | 596.4066 | 2.7755 | dbspec-03-Mar-2023-12-40-9.218 | Dr. Tomaso (FLI Jena) |
| 6 | Burkholderia mallei ATCC 23344 300102RR0118 | 590.4928 | 2.7712 | dbspec-03-Mar-2023-12-41-29.653 | Dr. Tomaso (FLI Jena) |
| 7 | Burkholderia pseudomallei 03 04450 060406RR0740 | 589.1215 | 2.7702 | dbspec-03-Mar-2023-12-50-45.809 | Dr. Tomaso (FLI Jena) |
| 8 | Burkholderia mallei 3708 | 579.1518 | 2.7628 | dbspec-03-Mar-2023-12-45-32.914 | RKI ZBS6 |
| 9 | Burkholderia mallei BfR 237 061102RR0551 | 563.2867 | 2.7507 | dbspec-03-Mar-2023-12-42-14.512 | Dr. Tomaso (FLI Jena) |
| 10 | Burkholderia mallei NCTC 10260 041206RR1051 | 561.1409 | 2.7491 | dbspec-03-Mar-2023-12-47-0.642 | Dr. Tomaso (FLI Jena) |
| 11 | Burkholderia mallei BfR 34 281002RR0518 | 561.1126 | 2.7491 | dbspec-03-Mar-2023-12-43-53.666 | Dr. Tomaso (FLI Jena) |
| 12 | Burkholderia mallei BfR M1 290103RR0043 | 559.2992 | 2.7476 | dbspec-03-Mar-2023-12-44-39.196 | Dr. Tomaso (FLI Jena) |
| 13 | Burkholderia mallei Zagreb 080304RR0090 | 559.2127 | 2.7476 | dbspec-03-Mar-2023-12-40-32.655 | Dr. Tomaso (FLI Jena) |
| 14 | Burkholderia mallei BfR M2 041206RR1057 | 558.3277 | 2.7469 | dbspec-03-Mar-2023-12-44-57.446 | Dr. Tomaso (FLI Jena) |
| 15 | Burkholderia mallei type strain | 557.0162 | 2.7459 | dbspec-03-Mar-2023-12-48-34.734 | Dr. Tomaso (FLI Jena) |
| 16 | Burkholderia pseudomallei NCTC 1688 041206RR1062 | 540.4352 | 2.7327 | dbspec-03-Mar-2023-12-52-10.62 | Dr. Tomaso (FLI Jena) |
| 17 | Burkholderia pseudomallei ring trial A-269 01 A | 530.6296 | 2.7248 | dbspec-03-Mar-2023-12-50-22.872 | RKI ZBS2\|ZBS6 |
| 18 | Burkholderia mallei BfR 235 300102RR0104 | 528.5485 | 2.7231 | dbspec-03-Mar-2023-12-41-52.122 | Dr. Tomaso (FLI Jena) |
| 19 | Burkholderia mallei BfR M1 040203RR0053 | 528.2781 | 2.7229 | dbspec-03-Mar-2023-12-44-15.009 | Dr. Tomaso (FLI Jena) |
| 20 | Burkholderia mallei type strain 041206RR1054 | 528.2603 | 2.7228 | dbspec-03-Mar-2023-12-48-13.235 | Dr. Tomaso (FLI Jena) |

---

### # 15 - IDENTIFICATION REPORT FOR MALDI-TOF MS SPECTRUM *'2013_06_14_Maren_Stämmler_0007'*

#### Metadata of actual MALDI-ToF test spectrum

| | |
|---|---|
| **genus / species / strain:** | RKI MALDI sample 07, QUANDHIP EQAE ring trial, strains provided by RKI \| ZBS 2 |
| **file id:** | 2013_06_14_Maren_Stämmler_0007 |
| **type:** | Measurement 02 |
| **Bruker ID:** | 92FCB042-DB6D-4DCC-9783-56CB2B43BCD5 |
| **NCBI ID (primary):** | 13373 |
| **NCBI ID (secondary):** | 13373 |
| **growth time:** | Optimal growth time between 24 - 72h |
| **growth temperature:** | 37°C |
| **growth conditions:** | Optimal aerobic or microaerophilic conditions |
| **growth medium:** | Columbia blood agar (Oxoid), 2nd passage on TSA or Caso agar, harvested by the 2nd passage |
| **sample treatment:** | Sample mixed with 20 mkL 100Â§ TFA, final TFA conc. approx. 80 perc.; approx. 30 Min treatment time; diluted 1:10 (vol); mixed with 1:1 HCCA\|TA2(A) |
| **spores:** | No |
| **concentration:** | Pellet produced by centrifugation (1 x 5 Min, 15,000 rpm) of a 500 mkL cell suspension, gamma ray irradiated (30 kGy) |
| **extra info:** | Burkholderia mallei A106-3: microbial preparation by ZBS 2\| preparation for MALDI-ToF MS: M. Stämmler |
| **calibration standard:** | linear calibration using Escherichia coli DSM 3871 |
| **measurement method:** | D:\Methods\flexControlMethods\MaierMethods\ToM_200ns_20130611.par |
| **customer:** | RKI ZBS 2\| ZBS 6 |
| **measurement date/time:** | 2013-06-14T09:30:51.015+02:00 |
| **path to MS file:** | C:\Users\LaschP\Documents\MATLAB\Microbe MS testdata\ring trial RKI spectra\Sample_07\0_H16\1\1SLin |

#### Identification results: analysis of score ranking list

| No. | Genus/Species | Score | Log Score | UniProt Identifier | No. of strains in DB |
|---|---|---|---|---|---|
| 1 | Burkholderia mallei | 482.5109 | 2.6835 | 13373 | 33 |
| 2 | Burkholderia pseudomallei | 324.3962 | 2.5111 | 28450 | 20 |
| 3 | Burkholderia thailandensis | 5.6981e-10 | -9.2443 | 57975 | 15 |

#### Score ranking list: best matches with test spectrum

| No. | Genus/Species/Strain | Score | Log Score | Spectrum identifier | Customer, or UniprotKB link |
|---|---|---|---|---|---|
| 1 | Burkholderia pseudomallei PITT 5691 041206RR1064 | 660.2042 | 2.8197 | dbspec-03-Mar-2023-12-52-51.791 | Dr. Tomaso (FLI Jena) |
| 2 | Burkholderia mallei Dubai 7 | 655.4306 | 2.8165 | dbspec-03-Mar-2023-12-45-34.492 | Dr. Tomaso (FLI Jena) |
| 3 | Burkholderia mallei Dubai 7 240609RR5318 | 617.8711 | 2.7909 | dbspec-03-Mar-2023-12-39-40.093 | Dr. Tomaso (FLI Jena) |
| 4 | Burkholderia mallei ATCC 23344 251109RR8925 | 616.3707 | 2.7898 | dbspec-03-Mar-2023-12-41-0.185 | Dr. Tomaso (FLI Jena) |
| 5 | Burkholderia mallei type strain | 613.1832 | 2.7876 | dbspec-03-Mar-2023-12-48-34.734 | Dr. Tomaso (FLI Jena) |
| 6 | Burkholderia mallei Mukteswar 290103RR0041 | 592.6552 | 2.7728 | dbspec-03-Mar-2023-12-40-9.218 | Dr. Tomaso (FLI Jena) |
| 7 | Burkholderia mallei ATCC 23344 300102RR0118 | 589.1399 | 2.7702 | dbspec-03-Mar-2023-12-41-29.653 | Dr. Tomaso (FLI Jena) |
| 8 | Burkholderia mallei 3708 | 575.2289 | 2.7598 | dbspec-03-Mar-2023-12-45-32.914 | RKI ZBS6 |
| 9 | Burkholderia mallei BfR 237 061102RR0551 | 558.9882 | 2.7474 | dbspec-03-Mar-2023-12-42-14.512 | Dr. Tomaso (FLI Jena) |
| 10 | Burkholderia pseudomallei ring trial A-269 01 A | 556.7376 | 2.7457 | dbspec-03-Mar-2023-12-50-22.872 | RKI ZBS2\|ZBS6 |
| 11 | Burkholderia mallei BfR 34 281002RR0518 | 556.6912 | 2.7456 | dbspec-03-Mar-2023-12-43-53.666 | Dr. Tomaso (FLI Jena) |
| 12 | Burkholderia mallei NCTC 10260 041206RR1051 | 556.5506 | 2.7455 | dbspec-03-Mar-2023-12-47-0.642 | Dr. Tomaso (FLI Jena) |
| 13 | Burkholderia mallei BfR M1 290103RR0043 | 556.2202 | 2.7452 | dbspec-03-Mar-2023-12-44-39.196 | Dr. Tomaso (FLI Jena) |
| 14 | Burkholderia mallei Zagreb 080304RR0090 | 555.8616 | 2.745 | dbspec-03-Mar-2023-12-40-32.655 | Dr. Tomaso (FLI Jena) |
| 15 | Burkholderia pseudomallei 03 04450 060406RR0740 | 555.4464 | 2.7446 | dbspec-03-Mar-2023-12-50-45.809 | Dr. Tomaso (FLI Jena) |
| 16 | Burkholderia mallei BfR M1 040203RR0053 | 555.1852 | 2.7444 | dbspec-03-Mar-2023-12-44-15.009 | Dr. Tomaso (FLI Jena) |
| 17 | Burkholderia mallei Zagreb | 554.8834 | 2.7442 | dbspec-03-Mar-2023-12-47-48.61 | Dr. Tomaso (FLI Jena) |
| 18 | Burkholderia mallei BfR M2 041206RR1057 | 554.6351 | 2.744 | dbspec-03-Mar-2023-12-44-57.446 | Dr. Tomaso (FLI Jena) |
| 19 | Burkholderia mallei ATCC 23344 | 550.4419 | 2.7407 | dbspec-03-Mar-2023-12-40-53.201 | RKI ZBS6 |
| 20 | Burkholderia thailandensis E143 | 549.8897 | 2.7403 | dbspec-03-Mar-2023-12-58-28.784 | RKI ZBS6 |

---

### # 16 - IDENTIFICATION REPORT FOR MALDI-TOF MS SPECTRUM *'2013_06_17_Maren_Stämmler_0002'*

#### Metadata of actual MALDI-ToF test spectrum

| | |
|---|---|
| **genus / species / strain:** | RKI MALDI sample 08, QUANDHIP EQAE ring trial, strains provided by RKI \| ZBS 2 |
| **file id:** | 2013_06_17_Maren_Stämmler_0002 |
| **type:** | Measurement 01 |
| **Bruker ID:** | 8111BE2B-254B-4723-A815-87E8BF9DAC5F |
| **NCBI ID (primary):** | 57975 |
| **NCBI ID (secondary):** | 57975 |
| **growth time:** | Optimal growth time between 24 - 72h |
| **growth temperature:** | 37°C |
| **growth conditions:** | Optimal aerobic or microaerophilic conditions |
| **growth medium:** | Columbia blood agar (Oxoid), 2nd passage on TSA or Caso agar, harvested by the 2nd passage |
| **sample treatment:** | Sample mixed with 30 mkL 100Â§ TFA, final TFA conc. approx. 80 perc.; 10 mkL 80 perc. TFA added; approx. 30 Min treatment time; diluted 1:10 (vol); mixed with 1:1 HCCA\|TA2(A) |
| **spores:** | No |
| **concentration:** | Pellet produced by centrifugation (1 x 5 Min, 15,000 rpm) of a 500 mkL cell suspension, gamma ray irradiated (30 kGy) |
| **extra info:** | Burkholderia thailandensis E125: microbial preparation by ZBS 2\| preparation for MALDI-ToF MS: M. Stämmler |
| **calibration standard:** | linear calibration using Escherichia coli DSM 3871 |
| **measurement method:** | D:\Methods\flexControlMethods\MaierMethods\ToM_200ns_20130611.par |
| **customer:** | RKI ZBS 2\| ZBS 6 |
| **measurement date/time:** | 2013-06-17T09:23:57.843+02:00 |
| **path to MS file:** | C:\Users\LaschP\Documents\MATLAB\Microbe MS testdata\ring trial RKI spectra\Sample_08\0_E20\1\1SLin |

#### Identification results: analysis of score ranking list

| No. | Genus/Species | Score | Log Score | UniProt Identifier | No. of strains in DB |
|---|---|---|---|---|---|
| 1 | Burkholderia thailandensis | 798.7223 | 2.9024 | 57975 | 15 |
| 2 | Burkholderia oklahomensis | 0.00089287 | -3.0492 | 342113 | 6 |
| 3 | Burkholderia pseudomallei | 0.00012428 | -3.9056 | 28450 | 20 |

#### Score ranking list: best matches with test spectrum

| No. | Genus/Species/Strain | Score | Log Score | Spectrum identifier | Customer, or UniprotKB link |
|---|---|---|---|---|---|
| 1 | Burkholderia thailandensis E125 | 654.6707 | 2.816 | dbspec-03-Mar-2023-12-58-16.128 | RKI ZBS6 |
| 2 | Burkholderia thailandensis DSM 13276 | 620.4202 | 2.7927 | dbspec-03-Mar-2023-12-58-1.034 | RKI ZBS6 |

| | | | | | |
|---|---|---|---|---|---|
| 3 | Burkholderia thailandensis E184 | 617.61 | 2.7907 | dbspec-03-Mar-2023-12-58-38.737 | RKI ZBS6 |
| 4 | Burkholderia thailandensis E131 | 617.3327 | 2.7905 | dbspec-03-Mar-2023-12-58-25.487 | RKI ZBS6 |
| 5 | Burkholderia thailandensis E207 | 590.156 | 2.771 | dbspec-03-Mar-2023-12-58-45.393 | RKI ZBS6 |
| 6 | Burkholderia thailandensis E067 | 588.6388 | 2.7698 | dbspec-03-Mar-2023-12-58-12.971 | RKI ZBS6 |
| 7 | Burkholderia thailandensis E202 | 588.1626 | 2.7695 | dbspec-03-Mar-2023-12-58-42.033 | RKI ZBS6 |
| 8 | Burkholderia thailandensis E163 | 586.1668 | 2.768 | dbspec-03-Mar-2023-12-58-35.549 | RKI ZBS6 |
| 9 | Burkholderia thailandensis E049 | 583.3243 | 2.7659 | dbspec-03-Mar-2023-12-58-6.394 | RKI ZBS6 |
| 10 | Burkholderia thailandensis E143 | 556.6173 | 2.7456 | dbspec-03-Mar-2023-12-58-28.784 | RKI ZBS6 |
| 11 | Burkholderia thailandensis E153 | 552.8257 | 2.7426 | dbspec-03-Mar-2023-12-58-32.096 | RKI ZBS6 |
| 12 | Burkholderia thailandensis 090804RR0288 135 | 516.4125 | 2.713 | dbspec-03-Mar-2023-12-57-39.41 | Dr. Tomaso (FLI Jena) |
| 13 | Burkholderia thailandensis LMG 20219 | 511.6803 | 2.709 | dbspec-03-Mar-2023-12-58-48.596 | RKI ZBS2\|ZBS6 |
| 14 | Burkholderia thailandensis E058 | 490.458 | 2.6906 | dbspec-03-Mar-2023-12-58-9.659 | RKI ZBS6 |
| 15 | Burkholderia thailandensis ring trial A-269 07 A | 426.4398 | 2.6299 | dbspec-03-Mar-2023-12-57-59.206 | RKI ZBS2\|ZBS6 |
| 16 | Burkholderia oklahomensis LMG 23620 | 376.2324 | 2.5755 | dbspec-03-Mar-2023-12-50-9.373 | RKI ZBS2\|ZBS6 |
| 17 | Burkholderia pseudomallei A335-1 | 374.8907 | 2.5739 | dbspec-03-Mar-2023-12-51-15.574 | RKI ZBS2 |
| 18 | Burkholderia pseudomallei 4845 | 350.939 | 2.5452 | dbspec-03-Mar-2023-12-51-41.371 | RKI ZBS6 |
| 19 | Burkholderia pseudomallei PITT 5691 041206RR1064 | 347.5468 | 2.541 | dbspec-03-Mar-2023-12-52-51.791 | Dr. Tomaso (FLI Jena) |
| 20 | Burkholderia oklahomensis DSM 21774 | 344.8196 | 2.5376 | dbspec-03-Mar-2023-12-49-58.576 | RKI ZBS6 |

---

#### # 17 - IDENTIFICATION REPORT FOR MALDI-TOF MS SPECTRUM *'2013_06_13_Maren_Stämmler_0008'*

##### Metadata of actual MALDI-ToF test spectrum

| | |
|---|---|
| **genus / species / strain:** | RKI MALDI sample 08, QUANDHIP EQAE ring trial, strains provided by RKI \| ZBS 2 |
| **file id:** | 2013_06_13_Maren_Stämmler_0008 |
| **type:** | Measurement 02 |
| **Bruker ID:** | 70156A73-5473-4C86-A9AA-061FA9ACF803 |
| **NCBI ID (primary):** | 57975 |
| **NCBI ID (secondary):** | 57975 |
| **growth time:** | Optimal growth time between 24 - 72h |
| **growth temperature:** | 37°C |
| **growth conditions:** | Optimal aerobic or microaerophilic conditions |
| **growth medium:** | Columbia blood agar (Oxoid), 2nd passage on TSA or Caso agar, harvested by the 2nd passage |
| **sample treatment:** | Sample mixed with 20 mkL 100Â§ TFA, final TFA conc. approx. 80 perc.; approx. 30 Min treatment time; diluted 1:10 (vol); mixed with 1:1 HCCA\|TA2(A) |
| **spores:** | No |
| **concentration:** | Pellet produced by centrifugation (1 x 5 Min, 15,000 rpm) of a 500 mkL cell suspension, gamma ray irradiated (30 kGy) |
| **extra info:** | Burkholderia thailandensis E125: microbial preparation by ZBS 2\| preparation for MALDI-ToF MS: M. Stämmler |
| **calibration standard:** | linear calibration using Escherichia coli DSM 3871 |
| **measurement method:** | D:\Methods\flexControlMethods\MaierMethods\ToM_200ns_20130611.par |
| **customer:** | RKI ZBS 2\| ZBS 6 |
| **measurement date/time:** | 2013-06-13T14:48:34.328+02:00 |
| **path to MS file:** | C:\Users\LaschP\Documents\MATLAB\Microbe MS testdata\ring trial RKI spectra\Sample_08\0_H17\1\1SLin |

##### Identification results: analysis of score ranking list

| No. | Genus/Species | Score | Log Score | UniProt Identifier | No. of strains in DB |
|---|---|---|---|---|---|
| 1 | Burkholderia thailandensis | 879.2215 | 2.9441 | 57975 | 15 |
| 2 | Burkholderia pseudomallei | 0.0011122 | -2.9538 | 28450 | 20 |
| 3 | Burkholderia mallei | 9.8068e-06 | -5.0085 | 13373 | 33 |
| 4 | Burkholderia oklahomensis | 2.6284e-07 | -6.5803 | 342113 | 6 |

##### Score ranking list: best matches with test spectrum

| No. | Genus/Species/Strain | Score | Log Score | Spectrum identifier | Customer, or UniprotKB link |
|---|---|---|---|---|---|
| 1 | Burkholderia thailandensis DSM 13276 | 783.8194 | 2.8942 | dbspec-03-Mar-2023-12-58-1.034 | RKI ZBS6 |
| 2 | Burkholderia thailandensis E184 | 752.5735 | 2.8765 | dbspec-03-Mar-2023-12-58-38.737 | RKI ZBS6 |
| 3 | Burkholderia thailandensis E131 | 750.8429 | 2.8755 | dbspec-03-Mar-2023-12-58-25.487 | RKI ZBS6 |
| 4 | Burkholderia thailandensis E067 | 722.7416 | 2.859 | dbspec-03-Mar-2023-12-58-12.971 | RKI ZBS6 |
| 5 | Burkholderia thailandensis E153 | 720.3183 | 2.8575 | dbspec-03-Mar-2023-12-58-32.096 | RKI ZBS6 |
| 6 | Burkholderia thailandensis E125 | 720.2077 | 2.8575 | dbspec-03-Mar-2023-12-58-16.128 | RKI ZBS6 |
| 7 | Burkholderia thailandensis E049 | 712.9565 | 2.8531 | dbspec-03-Mar-2023-12-58-6.394 | RKI ZBS6 |
| 8 | Burkholderia thailandensis E207 | 692.3583 | 2.8403 | dbspec-03-Mar-2023-12-58-45.393 | RKI ZBS6 |
| 9 | Burkholderia thailandensis E163 | 691.9648 | 2.8401 | dbspec-03-Mar-2023-12-58-35.549 | RKI ZBS6 |
| 10 | Burkholderia thailandensis E143 | 689.619 | 2.8386 | dbspec-03-Mar-2023-12-58-28.784 | RKI ZBS6 |
| 11 | Burkholderia thailandensis E202 | 661.6497 | 2.8206 | dbspec-03-Mar-2023-12-58-42.033 | RKI ZBS6 |
| 12 | Burkholderia thailandensis E058 | 628.6624 | 2.7984 | dbspec-03-Mar-2023-12-58-9.659 | RKI ZBS6 |
| 13 | Burkholderia thailandensis 090804RR0288 135 | 619.8471 | 2.7923 | dbspec-03-Mar-2023-12-57-39.41 | Dr. Tomaso (FLI Jena) |
| 14 | Burkholderia thailandensis LMG 20219 | 574.592 | 2.7594 | dbspec-03-Mar-2023-12-58-48.596 | RKI ZBS2\|ZBS6 |
| 15 | Burkholderia thailandensis ring trial A-269 07 A | 529.755 | 2.7241 | dbspec-03-Mar-2023-12-57-59.206 | RKI ZBS2\|ZBS6 |
| 16 | Burkholderia pseudomallei PITT 5691 041206RR1064 | 449.418 | 2.6527 | dbspec-03-Mar-2023-12-52-51.791 | Dr. Tomaso (FLI Jena) |
| 17 | Burkholderia pseudomallei PITT 521 041206RR1059 | 445.9618 | 2.6493 | dbspec-03-Mar-2023-12-52-34.948 | Dr. Tomaso (FLI Jena) |

| | | | | | |
|---|---|---|---|---|---|
| 18 | Burkholderia mallei BfR M2 041206RR1057 | 443.8691 | 2.6473 | dbspec-03-Mar-2023-12-44-57.446 | Dr. Tomaso (FLI Jena) |
| 19 | Burkholderia oklahomensis DSM 21774 | 442.8163 | 2.6462 | dbspec-03-Mar-2023-12-49-58.576 | RKI ZBS6 |
| 20 | Burkholderia mallei ATCC 23344 251109RR8925 | 441.4933 | 2.6449 | dbspec-03-Mar-2023-12-41-0.185 | Dr. Tomaso (FLI Jena) |

#### # 18 - IDENTIFICATION REPORT FOR MALDI-TOF MS SPECTRUM *'2013_06_14_Maren_Stämmler_0008'*

##### Metadata of actual MALDI-ToF test spectrum

| | |
|---|---|
| **genus / species / strain:** | RKI MALDI sample 08, QUANDHIP EQAE ring trial, strains provided by RKI \| ZBS 2 |
| **file id:** | 2013_06_14_Maren_Stämmler_0008 |
| **type:** | Measurement 03 |
| **Bruker ID:** | BE203424-ED2C-4A25-BA68-10D562636EDE |
| **NCBI ID (primary):** | 57975 |
| **NCBI ID (secondary):** | 57975 |
| **growth time:** | Optimal growth time between 24 - 72h |
| **growth temperature:** | 37°C |
| **growth conditions:** | Optimal aerobic or microaerophilic conditions |
| **growth medium:** | Columbia blood agar (Oxoid), 2nd passage on TSA or Caso agar, harvested by the 2nd passage |
| **sample treatment:** | Sample mixed with 20 mkL 100Â§ TFA, final TFA conc. approx. 80 perc.; approx. 30 Min treatment time; diluted 1:10 (vol); mixed with 1:1 HCCA\|TA2(A) |
| **spores:** | No |
| **concentration:** | Pellet produced by centrifugation (1 x 5 Min, 15,000 rpm) of a 500 mkL cell suspension, gamma ray irradiated (30 kGy) |
| **extra info:** | Burkholderia thailandensis E125: microbial preparation by ZBS 2\| preparation for MALDI-ToF MS: M. Stämmler |
| **calibration standard:** | linear calibration using Escherichia coli DSM 3871 |
| **measurement method:** | D:\Methods\flexControlMethods\MaierMethods\ToM_200ns_20130611.par |
| **customer:** | RKI ZBS 2\| ZBS 6 |
| **measurement date/time:** | 2013-06-14T09:36:55.281+02:00 |
| **path to MS file:** | C:\Users\LaschP\Documents\MATLAB\Microbe MS testdata\ring trial RKI spectra\Sample_08\0_H18\1\1SLin |

##### Identification results: analysis of score ranking list

| No. | Genus/Species | Score | Log Score | UniProt Identifier | No. of strains in DB |
|---|---|---|---|---|---|
| 1 | Burkholderia thailandensis | 878.4209 | 2.9437 | 57975 | 15 |
| 2 | Burkholderia mallei | 0.0011567 | -2.9368 | 13373 | 33 |
| 3 | Burkholderia pseudomallei | 1.0227e-05 | -4.9902 | 28450 | 20 |
| 4 | Burkholderia oklahomensis | 5.3019e-10 | -9.2756 | 342113 | 6 |

##### Score ranking list: best matches with test spectrum

| No. | Genus/Species/Strain | Score | Log Score | Spectrum identifier | Customer, or UniprotKB link |
|---|---|---|---|---|---|
| 1 | Burkholderia thailandensis E131 | 774.5827 | 2.8891 | dbspec-03-Mar-2023-12-58-25.487 | RKI ZBS6 |
| 2 | Burkholderia thailandensis E125 | 748.6123 | 2.8743 | dbspec-03-Mar-2023-12-58-16.128 | RKI ZBS6 |
| 3 | Burkholderia thailandensis E184 | 746.4627 | 2.873 | dbspec-03-Mar-2023-12-58-38.737 | RKI ZBS6 |
| 4 | Burkholderia thailandensis E067 | 746.4608 | 2.873 | dbspec-03-Mar-2023-12-58-12.971 | RKI ZBS6 |
| 5 | Burkholderia thailandensis DSM 13276 | 744.8663 | 2.8721 | dbspec-03-Mar-2023-12-58-1.034 | RKI ZBS6 |
| 6 | Burkholderia thailandensis E202 | 717.0958 | 2.8556 | dbspec-03-Mar-2023-12-58-42.033 | RKI ZBS6 |
| 7 | Burkholderia thailandensis E163 | 716.438 | 2.8552 | dbspec-03-Mar-2023-12-58-35.549 | RKI ZBS6 |
| 8 | Burkholderia thailandensis E153 | 715.144 | 2.8544 | dbspec-03-Mar-2023-12-58-32.096 | RKI ZBS6 |
| 9 | Burkholderia thailandensis E049 | 711.9137 | 2.8524 | dbspec-03-Mar-2023-12-58-6.394 | RKI ZBS6 |
| 10 | Burkholderia thailandensis E207 | 682.1171 | 2.8339 | dbspec-03-Mar-2023-12-58-45.393 | RKI ZBS6 |
| 11 | Burkholderia thailandensis E058 | 652.0314 | 2.8143 | dbspec-03-Mar-2023-12-58-9.659 | RKI ZBS6 |
| 12 | Burkholderia thailandensis E143 | 651.5391 | 2.8139 | dbspec-03-Mar-2023-12-58-28.784 | RKI ZBS6 |
| 13 | Burkholderia thailandensis 090804RR0288 135 | 646.341 | 2.8105 | dbspec-03-Mar-2023-12-57-39.41 | Dr. Tomaso (FLI Jena) |
| 14 | Burkholderia thailandensis LMG 20219 | 600.4017 | 2.7784 | dbspec-03-Mar-2023-12-58-48.596 | RKI ZBS2\|ZBS6 |
| 15 | Burkholderia thailandensis ring trial A-269 07 A | 556.2438 | 2.7453 | dbspec-03-Mar-2023-12-57-59.206 | RKI ZBS2\|ZBS6 |
| 16 | Burkholderia mallei ATCC 23344 251109RR8925 | 476.6687 | 2.6782 | dbspec-03-Mar-2023-12-41-0.185 | Dr. Tomaso (FLI Jena) |
| 17 | Burkholderia mallei BfR M2 041206RR1057 | 476.6457 | 2.6782 | dbspec-03-Mar-2023-12-44-57.446 | Dr. Tomaso (FLI Jena) |
| 18 | Burkholderia pseudomallei PITT 521 041206RR1059 | 473.2051 | 2.675 | dbspec-03-Mar-2023-12-52-34.948 | Dr. Tomaso (FLI Jena) |
| 19 | Burkholderia mallei Dubai 7 240609RR5318 | 466.6165 | 2.669 | dbspec-03-Mar-2023-12-39-40.093 | Dr. Tomaso (FLI Jena) |
| 20 | Burkholderia oklahomensis DSM 21774 | 464.9486 | 2.6674 | dbspec-03-Mar-2023-12-49-58.576 | RKI ZBS6 |

#### # 19 - IDENTIFICATION REPORT FOR MALDI-TOF MS SPECTRUM *'2013_06_17_Maren_Stämmler_0003'*

##### Metadata of actual MALDI-ToF test spectrum

| | |
|---|---|
| **genus / species / strain:** | RKI MALDI sample 09, QUANDHIP EQAE ring trial, strains provided by RKI \| ZBS 2 |
| **file id:** | 2013_06_17_Maren_Stämmler_0003 |
| **type:** | Measurement 01 |
| **Bruker ID:** | 8EEA47D8-C42B-4229-9813-6DBBF397112D |
| **NCBI ID (primary):** | 632 |

| | |
|---|---|
| **NCBI ID (secondary):** | 632 |
| **growth time:** | Optimal growth time between 24 - 72h |
| **growth temperature:** | 37°C |
| **growth conditions:** | Optimal aerobic or microaerophilic conditions |
| **growth medium:** | Columbia blood agar (Oxoid), 2nd passage on TSA or Caso agar, harvested by the 2nd passage |
| **sample treatment:** | Sample mixed with 30 mkL 100Â§ TFA, final TFA conc. approx. 80 perc.; approx. 30 Min treatment time; diluted 1:10 (vol); mixed with 1:1 HCCA\|TA2(A) |
| **spores:** | No |
| **concentration:** | Pellet produced by centrifugation (1 x 5 Min, 15,000 rpm) of a 500 mkL cell suspension, gamma ray irradiated (30 kGy) |
| **extra info:** | Yersinia pestis A106-2: microbial preparation by ZBS 2\| preparation for MALDI-ToF MS: M. Stämmler |
| **calibration standard:** | linear calibration using Escherichia coli DSM 3871 |
| **measurement method:** | D:\Methods\flexControlMethods\MaierMethods\ToM_200ns_20130611.par |
| **customer:** | RKI ZBS 2\| ZBS 6 |
| **measurement date/time:** | 2013-06-17T09:29:00.578+02:00 |
| **path to MS file:** | C:\Users\LaschP\Documents\MATLAB\Microbe MS testdata\ring trial RKI spectra\Sample_09\0_E21\1\1SLin |

#### Identification results: analysis of score ranking list

| No. | Genus/Species | Score | Log Score | UniProt Identifier | No. of strains in DB |
|---|---|---|---|---|---|
| 1 | Yersinia pestis | 396.4475 | 2.5982 | 632 | 11 |
| 2 | Yersinia pseudotuberculosis | 72.5794 | 1.8608 | 633 | 24 |
| 3 | Bacillus anthracis | 5.8273 | 0.76547 | 1392 | 130 |
| 4 | Yersinia enterocolitica | 0.6458 | -0.18991 | 630 | 56 |
| 5 | Yersinia similis | 0.16473 | -0.78322 | 367190 | 2 |

#### Score ranking list: best matches with test spectrum

| No. | Genus/Species/Strain | Score | Log Score | Spectrum identifier | Customer, or UniprotKB link |
|---|---|---|---|---|---|
| 1 | Yersinia pestis O3-01501 | 320.4904 | 2.5058 | dbspec-03-Mar-2023-14-0-6.944 | RKI ZBS6 |
| 2 | Yersinia pestis O3-01500 | 313.1871 | 2.4958 | dbspec-03-Mar-2023-14-0-3.616 | RKI ZBS6 |
| 3 | Yersinia pestis CCUG EV 76, CCUG 32133 | 304.2455 | 2.4832 | dbspec-03-Mar-2023-13-58-44.446 | RKI ZBS6 |
| 4 | Yersinia pseudotuberculosis Ring trial A-269 04 B | 251.2555 | 2.4001 | dbspec-03-Mar-2023-14-1-30.489 | RKI ZBS2\|ZBS6 |
| 5 | Yersinia pseudotuberculosis Typ3 P- INV+ | 242.3905 | 2.3845 | dbspec-03-Mar-2023-14-1-47.411 | RKI ZBS6 |
| 6 | Yersinia pseudotuberculosis 07PW12234 | 235.387 | 2.3718 | dbspec-03-Mar-2023-14-0-54.053 | RKI ZBS6 |
| 7 | Bacillus anthracis Sterne vaccine (A123, Beyer) | 232.4975 | 2.3664 | dbspec-03-Mar-2023-11-7-43.918 | RKI ZBS6, Dr. Beyer (Uni Hohenheim) |
| 8 | Yersinia pseudotuberculosis 04PA01423 | 231.5902 | 2.3647 | dbspec-03-Mar-2023-14-0-31.741 | RKI ZBS6 |
| 9 | Yersinia pseudotuberculosis J9 | 227.0398 | 2.3561 | dbspec-03-Mar-2023-14-1-27.115 | RKI ZBS6 |
| 10 | Yersinia enterocolitica Ring trial A-269 06 B | 226.8887 | 2.3558 | dbspec-03-Mar-2023-13-54-5.952 | RKI ZBS2\|ZBS6 |
| 11 | Yersinia pseudotuberculosis 27705 | 226.8203 | 2.3557 | dbspec-03-Mar-2023-14-1-7.881 | RKI ZBS6 |
| 12 | Yersinia similis DSM 18211, LMG 23763, CCUG 52882 | 226.4196 | 2.3549 | dbspec-03-Mar-2023-14-2-21.16 | CVUA Stuttgart |
| 13 | Yersinia pseudotuberculosis 27707 | 225.8674 | 2.3539 | dbspec-03-Mar-2023-14-1-11.209 | RKI ZBS6 |
| 14 | Yersinia pseudotuberculosis 25858 | 224.8508 | 2.3519 | dbspec-03-Mar-2023-14-1-4.677 | RKI ZBS6 |
| 15 | Yersinia pseudotuberculosis 25743 | 223.861 | 2.35 | dbspec-03-Mar-2023-14-1-1.334 | RKI ZBS6 |
| 16 | Yersinia pseudotuberculosis CNCTC 22\|90 | 223.8274 | 2.3499 | dbspec-03-Mar-2023-14-1-32.942 | RKI ZBS6 |
| 17 | Yersinia pseudotuberculosis 07WI00989 | 223.6974 | 2.3497 | dbspec-03-Mar-2023-14-0-57.99 | RKI ZBS6 |
| 18 | Yersinia pseudotuberculosis 29490 | 219.8012 | 2.342 | dbspec-03-Mar-2023-14-1-20.802 | RKI ZBS6 |
| 19 | Yersinia enterocolitica O:3 63\|1 | 215.0014 | 2.3324 | dbspec-03-Mar-2023-13-54-52.279 | RKI ZBS6 |
| 20 | Yersinia pestis RV 3 | 214.3698 | 2.3312 | dbspec-03-Mar-2023-14-0-30.334 | RKI ZBS6 |

---

### # 20 - IDENTIFICATION REPORT FOR MALDI-TOF MS SPECTRUM *'2013_06_13_Maren_Stämmler_0009'*

#### Metadata of actual MALDI-ToF test spectrum

| | |
|---|---|
| **genus / species / strain:** | RKI MALDI sample 09, QUANDHIP EQAE ring trial, strains provided by RKI \| ZBS 2 |
| **file id:** | 2013_06_13_Maren_Stämmler_0009 |
| **type:** | Measurement 02 |
| **Bruker ID:** | 70E47FB9-207D-4ABA-83A5-5463D6F150AA |
| **NCBI ID (primary):** | 632 |
| **NCBI ID (secondary):** | 632 |
| **growth time:** | Optimal growth time between 24 - 72h |
| **growth temperature:** | 37°C |
| **growth conditions:** | Optimal aerobic or microaerophilic conditions |
| **growth medium:** | Columbia blood agar (Oxoid), 2nd passage on TSA or Caso agar, harvested by the 2nd passage |
| **sample treatment:** | Sample mixed with 15 mkL 100Â§ TFA, final TFA conc. approx. 80 perc.; approx. 30 Min treatment time; diluted 1:10 (vol); mixed with 1:1 HCCA\|TA2(A) |
| **spores:** | No |
| **concentration:** | Pellet produced by centrifugation (1 x 5 Min, 15,000 rpm) of a 500 mkL cell suspension, gamma ray irradiated (30 kGy) |
| **extra info:** | Yersinia pestis A106-2: microbial preparation by ZBS 2\| preparation for MALDI-ToF MS: M. Stämmler |
| **calibration standard:** | linear calibration using Escherichia coli DSM 3871 |
| **measurement method:** | D:\Methods\flexControlMethods\MaierMethods\ToM_200ns_20130611.par |
| **customer:** | RKI ZBS 2\| ZBS 6 |
| **measurement date/time:** | 2013-06-13T14:57:12.250+02:00 |
| **path to MS file:** | C:\Users\LaschP\Documents\MATLAB\Microbe MS testdata\ring trial RKI spectra\Sample_09\0_H19\1\1SLin |

#### Identification results: analysis of score ranking list

| No. | Genus/Species | Score | Log Score | UniProt Identifier | No. of strains in DB |
|---|---|---|---|---|---|
| 1 | Yersinia pestis | 432.1668 | 2.6357 | 632 | 11 |
| 2 | Yersinia similis | 73.5492 | 1.8666 | 367190 | 2 |
| 3 | Yersinia pseudotuberculosis | 48.6377 | 1.687 | 633 | 24 |
| 4 | Yersinia enterocolitica | 3.7501 | 0.57404 | 630 | 56 |
| 5 | Yersinia pestis | 0.0032925 | -2.4825 | 632.1 | 8 |

#### Score ranking list: best matches with test spectrum

| No. | Genus/Species/Strain | Score | Log Score | Spectrum identifier | Customer, or UniprotKB link |
|---|---|---|---|---|---|
| 1 | Yersinia pestis O3-01501 | 362.173 | 2.5589 | dbspec-03-Mar-2023-14-0-6.944 | RKI ZBS6 |
| 2 | Yersinia pestis CCUG EV 76, CCUG 32133 | 344.455 | 2.5371 | dbspec-03-Mar-2023-13-58-44.446 | RKI ZBS6 |
| 3 | Yersinia pestis O3-01500 | 325.4395 | 2.5125 | dbspec-03-Mar-2023-14-0-3.616 | RKI ZBS6 |
| 4 | Yersinia similis DSM 18211, LMG 23763, CCUG 52882 | 314.6701 | 2.4979 | dbspec-03-Mar-2023-14-2-21.16 | CVUA Stuttgart |
| 5 | Yersinia pseudotuberculosis Ring trial A-269 04 B | 308.1834 | 2.4888 | dbspec-03-Mar-2023-14-1-30.489 | RKI ZBS2\|ZBS6 |
| 6 | Yersinia pseudotuberculosis Typ3 P- INV+ | 299.9801 | 2.4771 | dbspec-03-Mar-2023-14-1-47.411 | RKI ZBS6 |
| 7 | Yersinia pseudotuberculosis 04PA01423 | 289.6316 | 2.4618 | dbspec-03-Mar-2023-14-0-31.741 | RKI ZBS6 |
| 8 | Yersinia enterocolitica Ring trial A-269 06 B | 288.0197 | 2.4594 | dbspec-03-Mar-2023-13-54-5.952 | RKI ZBS2\|ZBS6 |
| 9 | Yersinia similis IMB 4354 | 285.5257 | 2.4556 | dbspec-03-Mar-2023-14-2-37.113 | Prof. Fuchs (ZIEL, Munich) |
| 10 | Yersinia pseudotuberculosis 27705 | 284.7028 | 2.4544 | dbspec-03-Mar-2023-14-1-7.881 | RKI ZBS6 |
| 11 | Yersinia pseudotuberculosis 25743 | 281.9385 | 2.4502 | dbspec-03-Mar-2023-14-1-1.334 | RKI ZBS6 |
| 12 | Yersinia pseudotuberculosis 07WI00989 | 281.6204 | 2.4497 | dbspec-03-Mar-2023-14-0-57.99 | RKI ZBS6 |
| 13 | Yersinia pestis NCTC 5923, ATCC 19428 | 276.3519 | 2.4415 | dbspec-03-Mar-2023-13-59-54.507 | Dr. Drevinek (Sujchbo, Czech Republic) |
| 14 | Yersinia pseudotuberculosis DSM 22972 | 268.0584 | 2.4282 | dbspec-03-Mar-2023-14-1-34.286 | RKI ZBS6 |
| 15 | Yersinia pestis O3-01502 | 263.9271 | 2.4215 | dbspec-03-Mar-2023-14-0-10.147 | RKI ZBS6 |
| 16 | Yersinia pseudotuberculosis 07PW12234 | 263.4473 | 2.4207 | dbspec-03-Mar-2023-14-0-54.053 | RKI ZBS6 |
| 17 | Yersinia pestis NCTC 10029 | 259.4877 | 2.4141 | dbspec-03-Mar-2023-13-58-45.946 | Dr. Drevinek (Sujchbo, Czech Republic) |
| 18 | Yersinia pseudotuberculosis J9 | 256.5179 | 2.4091 | dbspec-03-Mar-2023-14-1-27.115 | RKI ZBS6 |
| 19 | Yersinia pestis NCTC 2868 | 253.9289 | 2.4047 | dbspec-03-Mar-2023-13-59-28.93 | Dr. Drevinek (Sujchbo, Czech Republic) |
| 20 | Yersinia pseudotuberculosis 27707 | 253.4823 | 2.4039 | dbspec-03-Mar-2023-14-1-11.209 | RKI ZBS6 |

### # 21 - IDENTIFICATION REPORT FOR MALDI-TOF MS SPECTRUM *'2013_06_14_Maren_Stämmler_0009'*

#### Metadata of actual MALDI-ToF test spectrum

| | |
|---|---|
| **genus / species / strain:** | RKI MALDI sample 09, QUANDHIP EQAE ring trial, strains provided by RKI \| ZBS 2 |
| **file id:** | 2013_06_14_Maren_Stämmler_0009 |
| **type:** | Measurement 03 |
| **Bruker ID:** | 144E04DA-E6C7-4A7F-BD46-568E8747ADE2 |
| **NCBI ID (primary):** | 632 |
| **NCBI ID (secondary):** | 632 |
| **growth time:** | Optimal growth time between 24 - 72h |
| **growth temperature:** | 37°C |
| **growth conditions:** | Optimal aerobic or microaerophilic conditions |
| **growth medium:** | Columbia blood agar (Oxoid), 2nd passage on TSA or Caso agar, harvested by the 2nd passage |
| **sample treatment:** | Sample mixed with 15 mkL 100Â§ TFA, final TFA conc. approx. 80 perc.; approx. 30 Min treatment time; diluted 1:10 (vol); mixed with 1:1 HCCA\|TA2(A) |
| **spores:** | No |
| **concentration:** | Pellet produced by centrifugation (1 x 5 Min, 15,000 rpm) of a 500 mkL cell suspension, gamma ray irradiated (30 kGy) |
| **extra info:** | Yersinia pestis A106-2: microbial preparation by ZBS 2\| preparation for MALDI-ToF MS: M. Stämmler |
| **calibration standard:** | linear calibration using Escherichia coli DSM 3871 |
| **measurement method:** | D:\Methods\flexControlMethods\MaierMethods\ToM_200ns_20130611.par |
| **customer:** | RKI ZBS 2\| ZBS 6 |
| **measurement date/time:** | 2013-06-14T09:44:17.781+02:00 |
| **path to MS file:** | C:\Users\LaschP\Documents\MATLAB\Microbe MS testdata\ring trial RKI spectra\Sample_09\0_H20\1\1SLin |

#### Identification results: analysis of score ranking list

| No. | Genus/Species | Score | Log Score | UniProt Identifier | No. of strains in DB |
|---|---|---|---|---|---|
| 1 | Yersinia similis | 378.7337 | 2.5783 | 367190 | 2 |
| 2 | Yersinia pestis | 240.1012 | 2.3804 | 632 | 11 |
| 3 | Yersinia pseudotuberculosis | 114.5129 | 2.0589 | 633 | 24 |
| 4 | Yersinia enterocolitica | 2.1766 | 0.33778 | 630 | 56 |

#### Score ranking list: best matches with test spectrum

| No. | Genus/Species/Strain | Score | Log Score | Spectrum identifier | Customer, or UniprotKB link |
|---|---|---|---|---|---|
| 1 | Yersinia similis DSM 18211, LMG 23763, CCUG 52882 | 413.3943 | 2.6164 | dbspec-03-Mar-2023-14-2-21.16 | CVUA Stuttgart |
| 2 | Yersinia pestis CCUG EV 76, CCUG 32133 | 402.0335 | 2.6043 | dbspec-03-Mar-2023-13-58-44.446 | RKI ZBS6 |
| 3 | Yersinia pestis O3-01501 | 387.6835 | 2.5885 | dbspec-03-Mar-2023-14-0-6.944 | RKI ZBS6 |
| 4 | Yersinia pseudotuberculosis 27705 | 379.6871 | 2.5794 | dbspec-03-Mar-2023-14-1-7.881 | RKI ZBS6 |
| 5 | Yersinia pseudotuberculosis 25743 | 377.7477 | 2.5772 | dbspec-03-Mar-2023-14-1-1.334 | RKI ZBS6 |
| 6 | Yersinia pseudotuberculosis DSM 22972 | 364.8882 | 2.5622 | dbspec-03-Mar-2023-14-1-34.286 | RKI ZBS6 |
| 7 | Yersinia pseudotuberculosis 04PA01423 | 356.4687 | 2.552 | dbspec-03-Mar-2023-14-0-31.741 | RKI ZBS6 |
| 8 | Yersinia pseudotuberculosis 07PW12234 | 355.9207 | 2.5514 | dbspec-03-Mar-2023-14-0-54.053 | RKI ZBS6 |
| 9 | Yersinia enterocolitica Ring trial A-269 06 B | 353.2161 | 2.548 | dbspec-03-Mar-2023-13-54-5.952 | RKI ZBS2\|ZBS6 |
| 10 | Yersinia pseudotuberculosis J9 | 352.1583 | 2.5467 | dbspec-03-Mar-2023-14-1-27.115 | RKI ZBS6 |
| 11 | Yersinia similis IMB 4354 | 351.383 | 2.5458 | dbspec-03-Mar-2023-14-2-37.113 | Prof. Fuchs (ZIEL, Munich) |
| 12 | Yersinia pseudotuberculosis 07WI00989 | 350.2523 | 2.5444 | dbspec-03-Mar-2023-14-0-57.99 | RKI ZBS6 |
| 13 | Yersinia pestis O3-01500 | 349.0799 | 2.5429 | dbspec-03-Mar-2023-14-0-3.616 | RKI ZBS6 |
| 14 | Yersinia pseudotuberculosis 25858 | 348.9879 | 2.5428 | dbspec-03-Mar-2023-14-1-4.677 | RKI ZBS6 |
| 15 | Yersinia pseudotuberculosis 27707 | 348.9077 | 2.5427 | dbspec-03-Mar-2023-14-1-11.209 | RKI ZBS6 |
| 16 | Yersinia pseudotuberculosis Ring trial A-269 04 B | 345.9818 | 2.5391 | dbspec-03-Mar-2023-14-1-30.489 | RKI ZBS2\|ZBS6 |
| 17 | Yersinia pseudotuberculosis 29490 | 345.728 | 2.5387 | dbspec-03-Mar-2023-14-1-20.802 | RKI ZBS6 |
| 18 | Yersinia pseudotuberculosis 28928 | 345.4515 | 2.5384 | dbspec-03-Mar-2023-14-1-17.646 | RKI ZBS6 |
| 19 | Yersinia pseudotuberculosis DSM 8992, ATCC 29833, NCTC 10275, CIP 55.85 | 344.3363 | 2.537 | dbspec-03-Mar-2023-14-1-38.192 | RKI ZBS6 |
| 20 | Yersinia pestis NCTC 10029 | 343.6126 | 2.5361 | dbspec-03-Mar-2023-13-58-45.946 | Dr. Drevinek (Sujchbo, Czech Republic) |

---

### # 22 - IDENTIFICATION REPORT FOR MALDI-TOF MS SPECTRUM *'2013_06_17_Maren_Stämmler_0004'*

#### Metadata of actual MALDI-ToF test spectrum

| | |
|---|---|
| **genus / species / strain:** | RKI MALDI sample 10, QUANDHIP EQAE ring trial, strains provided by RKI \| ZBS 2 |
| **file id:** | 2013_06_17_Maren_Stämmler_0004 |
| **type:** | Measurement 01 |
| **Bruker ID:** | BF74B46D-3F52-4707-A3D6-5DC37A88756C |
| **NCBI ID (primary):** | 1428 |
| **NCBI ID (secondary):** | 1428 |
| **growth time:** | Optimal growth time between 24 - 72h |
| **growth temperature:** | 37°C |
| **growth conditions:** | Optimal aerobic or microaerophilic conditions |
| **growth medium:** | Columbia blood agar (Oxoid), 2nd passage on TSA or Caso agar, harvested by the 2nd passage |
| **sample treatment:** | Sample mixed with 80 mkL 100Â§ TFA, final TFA conc. approx. 80 perc.; approx. 30 Min treatment time; diluted 1:10 (vol); mixed with 1:1 HCCA\|TA2(A) |
| **spores:** | No |
| **concentration:** | Pellet produced by centrifugation (1 x 5 Min, 15,000 rpm) of a 500 mkL cell suspension, gamma ray irradiated (30 kGy) |
| **extra info:** | Bacillus thuringiensis DSM 350: microbial preparation by ZBS 2\| preparation for MALDI-ToF MS: M. Stämmler |
| **calibration standard:** | linear calibration using Escherichia coli DSM 3871 |
| **measurement method:** | D:\Methods\flexControlMethods\MaierMethods\ToM_200ns_20130611.par |
| **customer:** | RKI ZBS 2\| ZBS 6 |
| **measurement date/time:** | 2013-06-17T09:39:08.984+02:00 |
| **path to MS file:** | C:\Users\LaschP\Documents\MATLAB\Microbe MS testdata\ring trial RKI spectra\Sample_10\0_E22\1\1SLin |

#### Identification results: analysis of score ranking list

| No. | Genus/Species | Score | Log Score | UniProt Identifier | No. of strains in DB |
|---|---|---|---|---|---|
| 1 | Bacillus thuringiensis | 763.1728 | 2.8826 | 1428 | 20 |
| 2 | Bacillus cereus | 5.7675 | 0.76099 | 1396 | 102 |
| 3 | Bacillus anthracis | 1.7004 | 0.23056 | 1392 | 130 |
| 4 | Bacillus cereus s.l. | 0.066653 | -1.1762 | 86661 | 22 |
| 5 | Bacillus paramycoides | 0.030854 | -1.5107 | 2026194 | 1 |

#### Score ranking list: best matches with test spectrum

| No. | Genus/Species/Strain | Score | Log Score | Spectrum identifier | Customer, or UniprotKB link |
|---|---|---|---|---|---|
| 1 | Bacillus thuringiensis DSM 2046 (B188, Beyer) | 647.5043 | 2.8112 | dbspec-03-Mar-2023-11-32-48.217 | RKI ZBS6, Dr. Beyer (Uni Hohenheim) |
| 2 | Bacillus thuringiensis DSM 350 | 589.807 | 2.7707 | dbspec-03-Mar-2023-11-32-58.982 | RKI ZBS6 |
| 3 | Bacillus thuringiensis DSM 2046 | 555.3713 | 2.7446 | dbspec-03-Mar-2023-11-32-52.982 | RKI ZBS6 |
| 4 | Bacillus thuringiensis DSM 5815 | 525.3354 | 2.7204 | dbspec-03-Mar-2023-11-33-6.482 | RKI ZBS6 |
| 5 | Bacillus thuringiensis DSM 6890 | 519.1873 | 2.7153 | dbspec-03-Mar-2023-11-33-39.231 | RKI ZBS6 |
| 6 | Bacillus thuringiensis DSM 2046 (WS 2734) | 507.4934 | 2.7054 | dbspec-03-Mar-2023-11-35-1.151 | RKI ZBS6, Prof. Ehling-Schulz |
| 7 | Bacillus thuringiensis WS 2621 | 507.454 | 2.7054 | dbspec-03-Mar-2023-11-34-58.479 | RKI ZBS6, Prof. Ehling-Schulz |
| 8 | Bacillus cereus DSM 31 (RKI) | 481.9454 | 2.683 | dbspec-03-Mar-2023-11-18-56.351 | RKI ZBS6 |

| | | | | | |
|---|---|---|---|---|---|
| 9 | Bacillus thuringiensis DSM 350 (WIS-St. Nr. 315) | 465.2278 | 2.6677 | dbspec-03-Mar-2023-11-34-55.917 | RKI ZBS6 \|WIS (Dr. Niederwoehrmeier, Munster) |
| 10 | Bacillus anthracis unknown origin (A18, Beyer) | 453.7965 | 2.6569 | dbspec-03-Mar-2023-11-11-19.059 | RKI ZBS6, Dr. Beyer (Uni Hohenheim) |
| 11 | Bacillus anthracis unknown origin (A62, Beyer) | 449.2558 | 2.6525 | dbspec-03-Mar-2023-11-13-30.361 | RKI ZBS6, Dr. Beyer (Uni Hohenheim) |
| 12 | Bacillus cereus DSM 31 ATCC 14579 (B69, Beyer) | 441.9005 | 2.6453 | dbspec-03-Mar-2023-11-19-7.21 | RKI ZBS6, Dr. Beyer (Uni Hohenheim) |
| 13 | Bacillus cereus s.l. CD 3-1a | 441.7345 | 2.6452 | dbspec-03-Mar-2023-11-25-52.443 | RKI ZBS6 \| Dr. Thanh Tam (IPPR, Hanoi, Vietnam) |
| 14 | Bacillus paramycoides LMG 28876 | 434.7144 | 2.6382 | dbspec-03-Mar-2023-11-31-50.89 | RKI ZBS6 |
| 15 | Bacillus anthracis unknown origin (A5, Beyer) | 430.4381 | 2.6339 | dbspec-03-Mar-2023-11-13-7.803 | RKI ZBS6, Dr. Beyer (Uni Hohenheim) |
| 16 | Bacillus anthracis unknown origin (A28, Beyer) | 428.4303 | 2.6319 | dbspec-03-Mar-2023-11-11-49.856 | RKI ZBS6, Dr. Beyer (Uni Hohenheim) |
| 17 | Bacillus anthracis unknown origin (A9, Beyer) | 427.5194 | 2.631 | dbspec-03-Mar-2023-11-14-41.907 | RKI ZBS6, Dr. Beyer (Uni Hohenheim) |
| 18 | Bacillus anthracis unknown origin (A63, Beyer) | 426.1573 | 2.6296 | dbspec-03-Mar-2023-11-13-35.513 | RKI ZBS6, Dr. Beyer (Uni Hohenheim) |
| 19 | Bacillus anthracis ATCC 4229 (RKI) | 425.7851 | 2.6292 | dbspec-03-Mar-2023-11-6-56.498 | RKI ZBS2 |
| 20 | Bacillus anthracis unknown origin (A10, Beyer) | 425.7629 | 2.6292 | dbspec-03-Mar-2023-11-9-18.763 | RKI ZBS6, Dr. Beyer (Uni Hohenheim) |

---

#### # 23 - IDENTIFICATION REPORT FOR MALDI-TOF MS SPECTRUM *'2013_06_13_Maren_Stämmler_0010'*

##### Metadata of actual MALDI-ToF test spectrum

| | |
|---|---|
| **genus / species / strain:** | RKI MALDI sample 10, QUANDHIP EQAE ring trial, strains provided by RKI \| ZBS 2 |
| **file id:** | 2013_06_13_Maren_Stämmler_0010 |
| **type:** | Measurement 02 |
| **Bruker ID:** | 31336184-977D-4562-99B6-1F28627C6218 |
| **NCBI ID (primary):** | 1428 |
| **NCBI ID (secondary):** | 1428 |
| **growth time:** | Optimal growth time between 24 - 72h |
| **growth temperature:** | 37°C |
| **growth conditions:** | Optimal aerobic or microaerophilic conditions |
| **growth medium:** | Columbia blood agar (Oxoid), 2nd passage on TSA or Caso agar, harvested by the 2nd passage |
| **sample treatment:** | Sample mixed with 25 mkL 100Â§ TFA, final TFA conc. approx. 80 perc.; approx. 30 Min treatment time; diluted 1:10 (vol); mixed with 1:1 HCCA\|TA2(A) |
| **spores:** | No |
| **concentration:** | Pellet produced by centrifugation (1 x 5 Min, 15,000 rpm) of a 500 mkL cell suspension, gamma ray irradiated (30 kGy) |
| **extra info:** | Bacillus thuringiensis DSM 350: microbial preparation by ZBS 2\| preparation for MALDI-ToF MS: M. Stämmler |
| **calibration standard:** | linear calibration using Escherichia coli DSM 3871 |
| **measurement method:** | D:\Methods\flexControlMethods\MaierMethods\ToM_200ns_20130611.par |
| **customer:** | RKI ZBS 2\| ZBS 6 |
| **measurement date/time:** | 2013-06-13T15:05:30.796+02:00 |
| **path to MS file:** | C:\Users\LaschP\Documents\MATLAB\Microbe MS testdata\ring trial RKI spectra\Sample_10\0_H21\1\1SLin |

##### Identification results: analysis of score ranking list

| No. | Genus/Species | Score | Log Score | UniProt Identifier | No. of strains in DB |
|---|---|---|---|---|---|
| 1 | Bacillus thuringiensis | 789.288 | 2.8972 | 1428 | 20 |
| 2 | Bacillus cereus | 5.5789 | 0.74655 | 1396 | 102 |
| 3 | Bacillus cereus | 1.273 | 0.10483 | 1392.1 | 6 |
| 4 | Bacillus wiedmannii | 0.068291 | -1.1656 | 1890302 | 8 |
| 5 | Bacillus cereus s.l. | 0.020365 | -1.6911 | 86661 | 22 |

##### Score ranking list: best matches with test spectrum

| No. | Genus/Species/Strain | Score | Log Score | Spectrum identifier | Customer, or UniprotKB link |
|---|---|---|---|---|---|
| 1 | Bacillus thuringiensis DSM 2046 (B188, Beyer) | 649.2907 | 2.8124 | dbspec-03-Mar-2023-11-32-48.217 | RKI ZBS6, Dr. Beyer (Uni Hohenheim) |
| 2 | Bacillus thuringiensis DSM 6890 | 648.8551 | 2.8121 | dbspec-03-Mar-2023-11-33-39.231 | RKI ZBS6 |
| 3 | Bacillus thuringiensis DSM 350 | 628.0844 | 2.798 | dbspec-03-Mar-2023-11-32-58.982 | RKI ZBS6 |
| 4 | Bacillus thuringiensis DSM 2046 | 587.6428 | 2.7691 | dbspec-03-Mar-2023-11-32-52.982 | RKI ZBS6 |
| 5 | Bacillus thuringiensis DSM 5815 | 558.1047 | 2.7467 | dbspec-03-Mar-2023-11-33-6.482 | RKI ZBS6 |
| 6 | Bacillus thuringiensis WS 2621 | 516.1626 | 2.7128 | dbspec-03-Mar-2023-11-34-58.479 | RKI ZBS6, Prof. Ehling-Schulz |
| 7 | Bacillus thuringiensis DSM 2046 (WS 2734) | 514.4983 | 2.7114 | dbspec-03-Mar-2023-11-35-1.151 | RKI ZBS6, Prof. Ehling-Schulz |
| 8 | Bacillus cereus DSM 31 (RKI) | 482.4743 | 2.6835 | dbspec-03-Mar-2023-11-18-56.351 | RKI ZBS6 |
| 9 | Bacillus thuringiensis NMTD81 | 482.1891 | 2.6832 | dbspec-03-Mar-2023-11-34-42.214 | RKI ZBS6 \|Prof. Gao (Nanjing, China) |
| 10 | Bacillus cereus DSM 8438 (B248, Beyer) | 478.186 | 2.6796 | dbspec-03-Mar-2023-11-21-17.457 | RKI ZBS6, Dr. Beyer (Uni Hohenheim) |
| 11 | Bacillus thuringiensis GBSC29 (02) | 470.3387 | 2.6724 | dbspec-03-Mar-2023-11-34-14.402 | RKI ZBS6 \|Prof. Gao (Nanjing, China) |
| 12 | Bacillus thuringiensis DSM 5815 | 453.3026 | 2.6564 | dbspec-03-Mar-2023-11-33-10.357 | RKI ZBS6 |
| 13 | Bacillus wiedmannii GBSC45 | 453.2103 | 2.6563 | dbspec-03-Mar-2023-11-38-17.366 | RKI ZBS6 \|Prof. Gao (Nanjing, China) |
| 14 | Bacillus cereus s.l. unknown origin (B340, Beyer) | 452.525 | 2.6556 | dbspec-03-Mar-2023-11-27-11.535 | RKI ZBS6, Dr. Beyer (Uni Hohenheim) |
| 15 | Bacillus anthracis unknown origin (A62, Beyer) | 449.0012 | 2.6522 | dbspec-03-Mar-2023-11-13-30.361 | RKI ZBS6, Dr. Beyer (Uni Hohenheim) |
| 16 | Bacillus thuringiensis EZ05-05 | 448.0084 | 2.6513 | dbspec-03-Mar-2023-11-33-50.606 | RKI ZBS6 \|Prof. Gao (Nanjing, China) |
| 17 | Bacillus toyonensis NMTD92 | 447.5297 | 2.6508 | dbspec-03-Mar-2023-11-35-59.275 | RKI ZBS6 \|Prof. Gao (Nanjing, China) |
| 18 | Bacillus wiedmannii GBSC29 (01) | 444.5912 | 2.648 | dbspec-03-Mar-2023-11-38-1.069 | RKI ZBS6 \|Prof. Gao (Nanjing, China) |
| 19 | Bacillus thuringiensis DSM 350 (WIS-St. Nr. 315) | 436.8324 | 2.6403 | dbspec-03-Mar-2023-11-34-55.917 | RKI ZBS6 \|WIS (Dr. Niederwoehrmeier, Munster) |

| | | | | | |
|---|---|---|---|---|---|
| 20 | Bacillus toyonensis unknown origin (B260, Beyer) | 429.2994 | 2.6328 | dbspec-03-Mar-2023-11-36-40.243 | RKI ZBS6 \| Dr. Beyer (Uni Hohenheim) |

---

#### # 24 - IDENTIFICATION REPORT FOR MALDI-TOF MS SPECTRUM *'2013_06_14_Maren_Stämmler_0010'*

##### Metadata of actual MALDI-ToF test spectrum

| | |
|---|---|
| **genus / species / strain:** | RKI MALDI sample 10, QUANDHIP EQAE ring trial, strains provided by RKI \| ZBS 2 |
| **file id:** | 2013_06_14_Maren_Stämmler_0010 |
| **type:** | Measurement 03 |
| **Bruker ID:** | FCF8970F-FA71-4773-839A-DB9AB0101324 |
| **NCBI ID (primary):** | 1428 |
| **NCBI ID (secondary):** | 1428 |
| **growth time:** | Optimal growth time between 24 - 72h |
| **growth temperature:** | 37°C |
| **growth conditions:** | Optimal aerobic or microaerophilic conditions |
| **growth medium:** | Columbia blood agar (Oxoid), 2nd passage on TSA or Caso agar, harvested by the 2nd passage |
| **sample treatment:** | Sample mixed with 25 mkL 100Â§ TFA, final TFA conc. approx. 80 perc.; approx. 30 Min treatment time; diluted 1:10 (vol); mixed with 1:1 HCCA\|TA2(A) |
| **spores:** | No |
| **concentration:** | Pellet produced by centrifugation (1 x 5 Min, 15,000 rpm) of a 500 mkL cell suspension, gamma ray irradiated (30 kGy) |
| **extra info:** | Bacillus thuringiensis DSM 350: microbial preparation by ZBS 2\| preparation for MALDI-ToF MS: M. Stämmler |
| **calibration standard:** | linear calibration using Escherichia coli DSM 3871 |
| **measurement method:** | D:\Methods\flexControlMethods\MaierMethods\ToM_200ns_20130611.par |
| **customer:** | RKI ZBS 2\| ZBS 6 |
| **measurement date/time:** | 2013-06-14T09:51:48.343+02:00 |
| **path to MS file:** | C:\Users\LaschP\Documents\MATLAB\Microbe MS testdata\ring trial RKI spectra\Sample_10\0_H22\1\1SLin |

##### Identification results: analysis of score ranking list

| No. | Genus/Species | Score | Log Score | UniProt Identifier | No. of strains in DB |
|---|---|---|---|---|---|
| 1 | Bacillus thuringiensis | 738.8467 | 2.8686 | 1428 | 20 |
| 2 | Bacillus anthracis | 3.0192 | 0.4799 | 1392 | 130 |
| 3 | Bacillus toyonensis | 1.1155 | 0.047451 | 155322 | 16 |
| 4 | Bacillus wiedmannii | 0.00089513 | -3.0481 | 1890302 | 8 |
| 5 | Bacillus cereus | 4.4341e-10 | -9.3532 | 1396 | 102 |

##### Score ranking list: best matches with test spectrum

| No. | Genus/Species/Strain | Score | Log Score | Spectrum identifier | Customer, or UniprotKB link |
|---|---|---|---|---|---|
| 1 | Bacillus thuringiensis DSM 350 | 613.0602 | 2.7875 | dbspec-03-Mar-2023-11-32-58.982 | RKI ZBS6 |
| 2 | Bacillus thuringiensis DSM 6890 | 544.7107 | 2.7362 | dbspec-03-Mar-2023-11-33-39.231 | RKI ZBS6 |
| 3 | Bacillus thuringiensis DSM 2046 (B188, Beyer) | 539.4652 | 2.732 | dbspec-03-Mar-2023-11-32-48.217 | RKI ZBS6, Dr. Beyer (Uni Hohenheim) |
| 4 | Bacillus thuringiensis DSM 2046 | 480.3774 | 2.6816 | dbspec-03-Mar-2023-11-32-52.982 | RKI ZBS6 |
| 5 | Bacillus thuringiensis WS 2621 | 476.5257 | 2.6781 | dbspec-03-Mar-2023-11-34-58.479 | RKI ZBS6, Prof. Ehling-Schulz |
| 6 | Bacillus thuringiensis DSM 2046 (WS 2734) | 470.8335 | 2.6729 | dbspec-03-Mar-2023-11-35-1.151 | RKI ZBS6, Prof. Ehling-Schulz |
| 7 | Bacillus thuringiensis DSM 5815 | 453.506 | 2.6566 | dbspec-03-Mar-2023-11-33-6.482 | RKI ZBS6 |
| 8 | Bacillus thuringiensis NMTD81 | 441.5666 | 2.645 | dbspec-03-Mar-2023-11-34-42.214 | RKI ZBS6 \|Prof. Gao (Nanjing, China) |
| 9 | Bacillus anthracis unknown origin (A10, Beyer) | 417.5577 | 2.6207 | dbspec-03-Mar-2023-11-9-18.763 | RKI ZBS6, Dr. Beyer (Uni Hohenheim) |
| 10 | Bacillus toyonensis NMTD92 | 411.6986 | 2.6146 | dbspec-03-Mar-2023-11-35-59.275 | RKI ZBS6 \|Prof. Gao (Nanjing, China) |
| 11 | Bacillus anthracis unknown origin (A62, Beyer) | 408.1329 | 2.6108 | dbspec-03-Mar-2023-11-13-30.361 | RKI ZBS6, Dr. Beyer (Uni Hohenheim) |
| 12 | Bacillus thuringiensis GBSC29 (02) | 400.3971 | 2.6025 | dbspec-03-Mar-2023-11-34-14.402 | RKI ZBS6 \|Prof. Gao (Nanjing, China) |
| 13 | Bacillus anthracis unknown origin (A36, Beyer) | 390.1251 | 2.5912 | dbspec-03-Mar-2023-11-12-23.443 | RKI ZBS6, Dr. Beyer (Uni Hohenheim) |
| 14 | Bacillus anthracis unknown origin, blue (A88, Beyer) | 385.4542 | 2.586 | dbspec-03-Mar-2023-11-15-0.156 | RKI ZBS6, Dr. Beyer (Uni Hohenheim) |
| 15 | Bacillus anthracis unknown origin, black (A88, Beyer) | 385.3935 | 2.5859 | dbspec-03-Mar-2023-11-14-51.156 | RKI ZBS6, Dr. Beyer (Uni Hohenheim) |
| 16 | Bacillus wiedmannii GBSC45 | 384.9526 | 2.5854 | dbspec-03-Mar-2023-11-38-17.366 | RKI ZBS6 \|Prof. Gao (Nanjing, China) |
| 17 | Bacillus thuringiensis DSM 5815 | 381.0798 | 2.581 | dbspec-03-Mar-2023-11-33-10.357 | RKI ZBS6 |
| 18 | Bacillus wiedmannii GBSC29 (01) | 379.6447 | 2.5794 | dbspec-03-Mar-2023-11-38-1.069 | RKI ZBS6 \|Prof. Gao (Nanjing, China) |
| 19 | Bacillus thuringiensis EZ05-05 | 377.8397 | 2.5773 | dbspec-03-Mar-2023-11-33-50.606 | RKI ZBS6 \|Prof. Gao (Nanjing, China) |
| 20 | Bacillus cereus DSM 31 (RKI) | 376.7529 | 2.5761 | dbspec-03-Mar-2023-11-18-56.351 | RKI ZBS6 |

---
